# Niche partitioning and limited mobility characterise Middle Pleistocene kangaroos from eastern Australia

**DOI:** 10.1101/2025.10.30.685489

**Authors:** Christopher Laurikainen Gaete, Scott Hocknull, Clement P. Bataille, Andrew M. Lorrey, Katarina M. Mikac, Rochelle Lawrence, Anthony Dosseto

## Abstract

Australia’s Quaternary fossil record is characterised by a high diversity of macropodid taxa. Based on fossil faunal assemblages, it has been hypothesised many macropodids lived in sympatry during the Pleistocene, however, local geographic and dietary overlap is equivocal due to taphonomic uncertainty. Modern macropodid species rarely exhibit sympatry, suggesting anthropogenic or environmental changes may have disrupted these communities. Using Sr and C isotopes, we reconstruct foraging ranges and dietary preferences of several fossil macropodid lineages recovered in Middle Pleistocene cave deposits, at Mount Etna Caves, central eastern Queensland, Australia. Our results show that most macropodids, baring a single *Petrogale* potentially dispersing > 60 km, had limited foraging ranges and remained within 15 km of the fossil site. Moderate to large scale dispersal in individual *Petrogale* mirrors male-biased dispersal observed in some modern *Petrogale* populations indicating some individuals have the propensity to move between isolated colonies when corridors for dispersal are present. Smaller macropodids show dietary preferences similar to modern counterparts, while *Protemnodon* exhibit a division between C_3_/C_4_ intake, potentially indicating species-level differences. The analysis of geographic range and diet of this fossil assemblage reveals that macropodids partitioned on the landscape around the cave with a C_3_-dominant community – comprised of *Protemnodon, Petrogale,* and *Thylogale –* to the northwest, and a C_4_-dominant grassland community – comprised of *Notamacropus* and isolated members of *Protemnodon* and *Petrogale* – to the south. Therefore, we conclude, that although faunal assemblages alone suggest a larger number of macropodids living in sympatry, isotopic proxies uncover complex habitat partitioning between C_3_ and C_4_ environments around Mount Etna Caves.

## 1. Introduction

Kangaroos and wallabies (Macropodidae) have been a prominent part of Sahul (Australia and New Guinea) mammal fauna throughout much of the late Cenozoic (Kerr et al., 2024, Prideaux, 2004). During the Quaternary, macropodids exhibited remarkable diversity, including giant forms far exceeding the size of the largest extant species, such as the red kangaroo (*Osphranter rufus*; Prideaux, 2004, Hocknull et al., 2020, Kerr et al., 2024, Helgen et al., 2006). Despite their prominence in the fossil record and their ecological significance, research into the life histories of extinct macropodids remains limited, with palaeoecological studies often focused on iconic northern hemisphere megafauna species such as mammoths, bison, reindeer and smilodons (Wooller et al., 2021, Funck et al., 2021, Price et al., 2017b, Feranec and DeSantis, 2014). In contrast, ecology of extant macropodids is better understood with an observed diversification in body size (Telfer and Bowman, 2006), behaviour (Laws and Goldizen, 2003) and morphological structure (Warburton et al., 2011) allowing the group to colonise most environment, central arid deserts to coastal and highland rainforests s available across the continent of Sahul (Australia and New Guinea). Despite this diversity in behaviour, many modern macropodids reside in highly fragmented habitats with significant anthropogenic modifications making it unlikely that their observed behaviour and life history traits reflect those under ‘natural’ conditions (Heise-Pavlov, 2017, Newell, 1999b, Newell, 1999a, Procter-Gray, 1986). To address this, it is critical to reconstruct macropodid ecology prior to human arrival – particularly their foraging strategies, dietary preferences, and community structure – to establish baseline life history traits and better understand the evolutionary and ecological dynamics that shaped their past distributions (Dickman and Calver, 2023, Pires et al., 2018, Berzaghi et al., 2018).

Mount Etna Caves, located 27 km north of Rockhampton in central eastern Queensland (23°09’24" S, 150°26’58" E; Hocknull, 2005b), offers a unique opportunity to study diet, foraging ranges, and community dynamics across a range of macropodid taxa well before the arrival of humans on the Australian continent. The site comprises a series of limestone karst cave systems formed within isolated late Devonian limestone blocks, that preserve diverse vertebrate fossil assemblages that span middle to late Pleistocene (Deer, 2011, Hocknull, 2009, Hocknull et al., 2007, Hocknull, 2005b). Today, the region experiences a humid subtropical climate situated within the Brigalow Belt North Bioregion (Sprent and Sprent, 1970). Local vegetation is dominated by a semi-evergreen vine thicket (SEVT) or “dry rainforest” (Sprent and Sprent, 1970). Although once widespread, SEVT has been extensively cleared for agriculture, with only remnant patches persisting across limestone outcrops. Other native vegetation communities in the area include open forests, grasslands, and shrub woodlands (Martinez, 2010, Neldner, 2023). Fossil assemblages indicate that between 500,000 and 280,000 years ago (ka), the Mount Etna region supported a biodiverse rainforest ecosystem, representing the only known Quaternary rainforest vertebrate record in Australia (Hocknull et al., 2007, Hocknull, 2005b). This ecosystem is thought to have persisted until around 280–205 ka, when climatic shifts led to environmental changes that favoured xeric-adapted species, resulting in significant faunal turnover environment (Hocknull et al., 2007, Cramb et al., 2023, Louys et al., 2023, Cramb et al., 2020, Cramb et al., 2009).

In this study, we focus on several macropodid taxa of the following genera *Protemnodon, Notamacropus, Petrogale* and *Thylogale* residing at Mount Etna Caves in a stable rainforest environment between 280 – 330 ka. The primary aim is to reconstruct the foraging ranges and dietary preferences of these Middle Pleistocene macropodids using strontium (Sr) and carbon (C) isotopes. Historically, the co-occurrence of multiple species within a single fossil assemblage has been used to infer past sympatry or “disharmonious assemblages” (Lyman, 2016, Lundelius Jr, 1989). However, fossil accumulations often form diachronously, with deposition occurring over hundreds to tens of thousands of years. As such, spatial proximity in the fossil record may not accurately reflect ecological overlap at any given time. To address this, we test the hypothesis that macropodid diversity at Mount Etna Caves was driven by either geographic separation or niche partitioning across a mosaic landscape, rather than true sympatry. We predict that: (1) if macropodids were highly mobile, Sr isotope ratios will vary widely across individuals; and (2) if macropodid diversity was driven by dietary niche separation, then C isotope compositions will differ between taxa. These hypotheses are evaluated through direct isotopic characterisation of fossil tooth enamel, allowing us to infer individual-level habitat use and dietary behaviour.

Strontium isotopes (i.e., the ratio of ^87^Sr/^86^Sr) are a powerful tool for geo-provenance because they vary across the landscape with respect to the underlying geology. The Sr composition in herbivores reflects primarily that of the vegetation they consume (Flockhart et al., 2015) providing a powerful tool for examining foraging ranges (Crowley et al., 2017), with foraging range defined as the distance an individual travels to meet its dietary needs (Porolak et al., 2014). Strontium isotopes have been widely applied to examine foraging ranges in modern Caprine (Lazzerini et al., 2021), Caribou (Miller et al., 2021), and African Elephants (Koch et al., 1995) as well as extinct Lemuriformes (Crowley and Godfrey, 2019), Late Pleistocene Reindeer (Price et al., 2017b), and Mammoths and Mastodons in North America (Hoppe et al., 1999, Rowe et al., 2024), Central America (Pérez-Crespo et al., 2016) and Siberia (Arppe et al., 2009). In Australia, however, applications remain limited. Price et al. (2017a) suggested long-distance seasonal migrations in *Diprotodon optatum*, while Koutamanis et al. (2023) found restricted ranges in *Diprotodon*, *Procoptodon*, and *Protemnodon* from Bingara and Wellington Caves. These studies offer preliminary insights into Australian megafauna, but are constrained by a lack of baseline data predicting spatial Sr isotope variations. This can be addressed by large-scale sampling of Sr isotopes in vegetation (Bataille et al., 2020, Crowley et al., 2017), and/or mineralised tissue of extant, localised organisms (Funck et al., 2021, Adams et al., 2019), enabling the creation of a ‘strontium isoscape’, a map that encapsulates localised variation in Sr isotopes and provides more precise estimates of foraging across refined spatial scales (Funck et al., 2021, Bataille and Bowen, 2012, Bataille et al., 2020).

Carbon isotopes are widely used to reconstruct herbivore dietary preferences by distinguishing between plants that follow C_3_ and C_4_ photosynthetic pathways (Ehleringer and Cerling, 2002). C_4_ plants generally have higher δ¹³C values, averaging -12.1 ‰ (Ranging from -16 ‰ to -12 ‰), while C_3_ plants average -25 ‰ (Ranging from -37 ‰ to -20 ‰) (Boisserie et al., 2005, Munroe et al., 2022). Although isotopic fractionation occurs between vegetation and the herbivores (∼12 ‰) (DeNiro and Epstein, 1978, Montanari et al., 2013, Murphy et al., 2007), these isotopic differences are preserved allowing researchers to estimate the proportion of C_3_ versus C_4_ vegetation in an animal’s diet (Tieszen et al., 1983, DeNiro and Epstein, 1978).

In summary, direct isotopic characterisation of fossil specimens offers a way to capture individual distribution and habitat preferences. As more individuals are characterised, these cumulative results can be used to infer life history traits at the population level for a given spatiotemporal period. We hypothesise that across macropodid taxa, highly variable landscape use may explain the high level of faunal diversity observed in the Mount Etna Caves fossil record. If this hypothesis is valid, fossil tooth enamel should be characterised by highly variable Sr isotopes across individuals. Conversely, if most individuals were foraging locally, then the high diversity of taxa in the Mount Etna Caves faunal assemblages would reflect high levels of localised sympatry. In this case, we propose macropodid diversity would likely be driven by inter-specific niche diversification and predict individual taxa would be characterised by unique C isotope compositions.

## 2. Regional Setting

Fossil macropodids examined here were sourced from a series of cave deposits at Mount Etna Caves, central eastern Queensland (23°09′24″S 150°26′58″E) (Hocknull, 2005b). Large-scale stockpiling of key vertebrate deposits was undertaken while an active limestone mine operated, providing unique access to a large number of deposits. Fossils examined in this study were collected from four stratigraphic units (QML1311C/D, QML1311H, QML1384LU and QML1312) across two cave systems close to one another, on the western flank of Mount Etna: Speaking Tube Cave and Elephant Hole Cave (Supplementary Fig. S2). Fossil accumulation has been attributed to natural pitfall traps and predatory cave dwelling species, however, the macropodids by-in-large would have been derived from pit-fall accumulations due to their large body-size. Potentially some fossils may have been accumulated locally due to marsupial carnivore activities, however, bone modification by predators has not been observed and considered very rare (Hocknull et al., 2007, Martinez, 2010). It is also unlikely that the macropodids were sourced by the dominant cave-dwelling predators, the volant predators, owls, or ghost bats.

In the Speaking Tube Cave system, QML1311C/D and QML1311H occur at a similar level, separated by an older deposit, QML1311F (Hocknull, 2005b, Hocknull et al., 2007). QML1311C/D was bracket-dated with direct U-Th dating of fossil material producing an oldest minimum age of 280 +21/-18 ka (2σ), and thermally transferred optically stimulated luminescence (TT-OSL) dating of sediment providing a youngest maximum age of 291.5 ± 28.3 ka (1σ) (Laurikainen Gaete et al., 2025). Previous chronology established for QML1311H suggests this unit accumulated over a similar period of time with an oldest minimum age from U-series open system dating of macropodid teeth (285 ± 7 ka; 1σ), and a youngest maximum TT-OSL age of 304.2 ± 26.1 ka (1σ) (Hocknull et al., 2007, Laurikainen Gaete et al., 2025). This indicates that fossil deposition in both localities, QML1311H and QML1311C/D, occurred over a similar period in time, between 280 – 330 ka (Laurikainen Gaete et al., 2025). Faunal remains are diverse and include a wide range of vertebrates (Hocknull, 2017), including frogs (Hocknull, 2003); reptiles (Hocknull, 2005b, Hocknull et al., 2009); birds (Hocknull, 2005b); and marsupials including; kangaroos, possums, gliders, palorchestids, diprotodontids, (Hocknull, 2005b, Hocknull, 2005a), koalas (Price et al., 2009, Price and Hocknull, 2011), dasyurids, thylacinids and thylacoleonids (Cramb and Hocknull, 2010b, Cramb et al., 2023), and murids (Cramb and Hocknull, 2010a, Cramb et al., 2018).

Elephants Hole Cave contains a number of late Quaternary fossil deposits recovered throughout a vertically oriented cave system (Hocknull, 2005b). Stratigraphic positioning suggests the overlying deposit, QML1312 is significantly younger than underlying units QML1384UU and QML1384LU. Hocknull et al. (2007) bracket-dated QML1312 using U-series open system dating of macropodid teeth, and U-series closed-system dating from a reworked speleothem suggesting this deposit accumulated between 170 – 205 ka. For QML1384LU, Hocknull et al. (2007) suggested an oldest minimum age of 332 ± 14 ka (1σ) based on U-series open system dating of a calcite infill. Direct U-Th dating of macropodid tooth from Laurikainen Gaete et al. (2025) is in agreement with this oldest minimum age (291 +29/-30 ka; 2σ), however a lack of TT-OSL dating meant chronology could not be further constrained, and therefore deposition in QML1384LU is considered > 330 ka, predating other fossil localities considered here.

Bootstrap cluster analyses of faunal assemblages suggest that QML1384LU, QML1311H and QML1311C/D are associated with tropical rainforest environments in New Guinea and Australia’s Wet Tropics, whereas QML1312 is associated with modern, central Australian xeric fauna (Hocknull et al., 2007). This indicates that QML1384LU, QML1311H, and QML1311C/D represent fossil accumulation in a rainforest environment between 280 ka to >330 ka, while QML1312 reflects significant environmental transition towards xeric conditions accompanied by faunal turnover and an influx of arid adapted taxa reflects a dry-adapted fauna and the outcome of a significant environmental shift towards xeric conditions accompanied by faunal turnover, occurring sometime between 205–280 ka. In addition to faunal similarity, multiple independent lines of evidence support this interpretation of a stable rainforest habitat during this earlier time interval (280 – 330 ka). Deep-sea pollen records off the Queensland coast reveal significant rainforest elements during the Middle Pleistocene (Martin, 1993). This is consistent with phylogenetic affinities for several specialist arboreal mammalian lineages including *Dendrolagus, Pseudochirulus*, and *Phalanger* which show closer links to extant taxa residing in forested habitats in the New Guinea highlands (Hocknull et al., 2007, Hocknull, 2009). A rainforest habitat for several other taxa are also suggested, including signifiant body-size extremes in frogs (Hocknull, 2009, Hocknull, 2003); high diversity and specialisation in dasyurids (Cramb et al., 2009, Cramb and Hocknull, 2010b, Cramb et al., 2023); high diversity of bats including rainforest specialists (Martinez, 2010); forest and rainforest restricted rodents (Cramb et al., 2020, Hocknull et al., 2007, Hocknull, 2005b), and plesiomorphic rainforest koalas (Price and Hocknull, 2011).

Regional palaeoclimatic proxies provide further support for a prolonged period of climatic stability indicating that prior to a La Niña-dominated phase prior to 270 ka. Sea surface temperature gradients, n-alkane chain length indices, coccolith abundance, and planktonic foraminifera records all point towards a La Niña phases, with a transition to a drier El Niño phase around 270 ka (Li et al., 2015, López-Otálvaro et al., 2008, Gupta et al., 2010, Tian et al., 2022), within a period of low seasonality pre-280ka before the onset of significant seasonality beyond this (Windler et al., 2019, Taylor et al., 2022). These findings collectively indicate that the pre-280 ka environment at Mount Etna was characterised by a stable, forested environment with faunal assemblages suggesting similar conditions to modern rainforest systems in New Guinea or the Queensland Wet Tropics. By examining individuals of the only abundant macropodid taxa to occur in both rainforest and xeric deposits at Mount Etna Caves (i.e., *Petrogale*), we aim to explore whether diet and foraging ranges changed in *Petrogale* individuals in response to a drastic environmental transition between 205 – 280 ka.

## 3. Materials and Methods

### 3.1 Taxonomic assessment of fossil teeth

Due to the destructive nature of analysis, isolated molars and incisors were selected. Taxonomic allocation of these teeth by Scott Hocknull (SAH) was undertaken to the genus level through direct comparisons to modern comparative specimens and with better-preserved fossilised dental remains (e.g. maxillae and dentaries) identified from the same fossil localities (Hocknull, 2005b, Hocknull, 2009).

Identification of individuals was possible to the genus level but not equally possible across all groups at the species level. Therefore, we constrained our taxonomic precision of individuals to the genus level. Each generic level grouping also conformed to a sequence of generalised body sizes, with largest individuals represented by *Protemnodon*, then *Notamacropus*, *Petrogale,* and *Thylogale* being the smallest. Due to their rarity and particularities of preservation, other macropodine taxa known from the fossil deposits (Hocknull et al., 2020) were excluded from this study, however, will form part of future work.

### 3.2 Fossil sampling

This study examined 40 isolated fossil macropodid teeth representing at least 33 individuals (minimum number of individuals). These specimens represent four genera, including one that is entirely extinct, *Protemnodon* (n = 13); two extant taxa with species currently restricted to specialised habitats and at risk of extinction, *Petrogale* (n = 12) and *Thylogale* (n = 10), and a more cosmopolitan genus, *Notamacropus* (n = 5) (Supplementary material Table S1, Supplementary Fig. S3).

### 3.3 Vegetation sampling

To construct an isoscape of bioavailable strontium, 65 plant samples were collected representing 36 distinct geological units from an 10 000 km^2^ area surrounding Mount Etna Caves (Supplementary Material, Table S2). Plant samples were collected from the broader Rockhampton region between Byfield (22°54’42.86” S, 150°38’54.24” E), Gogango (23°39’58.46” S, 150°02’41.76” E), and Morinish (23°08’55.81” S, 149°58’16.84” E). To reduce anthropogenic influence on bioavailable strontium, areas with significant human activity, such as farmland and urban centres, were avoided to prevent contamination from pesticides, fertilisers, and other non-biological sources of strontium (Snoeck et al., 2020, Maurer et al., 2012).

### 3.4 Strontium isotopes

Both plant and enamel samples were prepared for Sr isotope analysis in a Class 10 Cleanroom, at the Wollongong Isotope Geochronology Laboratory (WIGL), University of Wollongong. Enamel powder from fossil specimens was collected using a Dremel^®^ rotary tool equipped with a 2-mm diamond-tipped conical drill bit (Roberts et al., 2019, Koutamanis et al., 2021). Trace-element (including REE) concentrations previously measured in individual *Protemnodon* from Mount Etna found little to diagenetic trace-element uptake in fossil enamel (Laurikainen Gaete et al., 2025). Therefore, it can be assumed, the Sr composition of enamel will represent bioaccumulated strontium. Contaminants on the outer surface of the enamel were removed with initial drill sweeps and 5 mg of powder was collected. For some specimens, two aliquots were taken from different parts of the tooth (from the top and base of the crown; Supplementary Fig. S3), to examine variations in ^87^Sr/^86^Sr isotope ratios within individuals during mineralisation (Supplementary Table S1). Enamel powder was digested in concentrated nitric acid (HNO_3_) (Balter et al., 2019, Hassler et al., 2021a, Hassler et al., 2021b, Martin et al., 2022, Koutamanis et al., 2021) before being redissolved in 2 M HNO_3_ prior to ion exchange chromatography.

Plant samples were allowed to dry naturally and then manually ground before being ashed at 550 °C using a CF1200 Muffle Furnace. Following this, 50 mg of ashed material was digested in a microwave system with 2 mL of 15 M HNO_3_. Samples were redissolved in 2 M HNO_3_ before ion exchange chromatography. Prior to chromatography a 1:100 dilution in 0.3 M HNO_3_ was prepared and for each sample to screen the Sr concentration using a Thermo iCAP Q quadrupole ICP-MS. Following this, 2 M HNO_3_ were re-diluted so all samples contained < 300 ng of Sr. Strontium was isolated from the sample matrix through automated low-pressure chromatography, using a prepFAST-MC™ equipped with a 1 mL Sr-Ca specific resin column (Balter et al., 2019, Martin et al., 2015, Martin et al., 2020, Martin et al., 2018, Tacail et al., 2014, Koutamanis et al., 2021). The Sr elution was evaporated to dryness before being redissolved in 0.3 M HNO_3_ prior to isotopic analysis.

Strontium isotope ratios were measured at WIGL using a Thermo-Fisher Neptune Plus MC-ICP-MS equipped with jet sample and X skimmer cones. Samples were introduced using cyclonic spray chamber equipped with an ESI Apex-St PFA MicroFlow nebuliser. Prior to analytical sessions, the instrument was tuned with a 20 ppb Sr solution of NIST SRM 987 standard, maximising sensitivity for ^88^Sr. Masses for ^85^Rb, ^86^Sr, ^87^Sr and ^88^Sr were collected on Faraday cups. Instrumental biases were corrected using the measured ^88^Sr/^86^Sr isotope ratio, and ^85^Rb was measured for isobaric interference corrections of ^87^Rb on ^87^Sr. Accuracy was assessed with repeated analyses of NIST SRM987 throughout analytical sessions. The mean measured ^87^Sr/^87^Sr isotope ratio for SRM987 (0.710253 ± 0.000004 (2SE), n = 32) was within error of published values in Weis et al. (2006) (0.710252 ± 0.000013 (2SE)). Procedural accuracy was assessed by processing an apple leaf NIST SRM1515 standard. The mean ^87^Sr/^86^Sr isotope ratio for SRM1515 (0.71389 ± 0.00007; 2SE, n = 2) was within error of the published value in Liu et al. (2016) (0.71398 ± 0.00004 (2SE)). Precision was assessed by preparing and processing multiple aliquots for several samples. For these replicates, repeat analyses were within 2SE of one another. Total procedure blanks ranged between 0.005 and 0.327 ng of Sr (n = 6), which is less than 0.00003% of the Sr processed in samples.

### 3.5 Carbon isotopes

Carbon isotopes were analysed in 21 fossil teeth of the following taxa: *Protemnodon* (n = 5), *Petrogale* (n = 8), *Thylogale* (n = 5), and *Notamacropus* (n = 3). For each sample, 2-3 mg of powdered enamel was collected using a Dremel rotary tool equipped with a 2-mm diamond tipped conical drill tip (Roberts et al., 2019, Koutamanis et al., 2021). Previous analyses of, trace-element (including REE) concentrations Mount Etna found little to diagenetic alteration of fossil enamel (Laurikainen Gaete et al., 2025). Therefore, it can be assumed, the C composition of enamel will represent bioaccumulated carbon.

Stable isotope analyses were carried out on a MAT253 dual inlet mass spectrometer (IRMS) linked to a Kiel IV carbonate device (Thermo-Fisher Scientific, Bremen, Germany) at the Stable Isotope Analytical Facility at the National Institute of Water and Atmospheric Research (NIWA), Wellington, New Zealand. Samples were reacted with 3 drops of H_3_PO_4_ at 75 °C in a Kiel automated individual carbonate reaction device coupled with the IRMS. Following analyses, using the International Atomic Energy Agency (IAEA) evaluation program ‘Stable Isotope calibration for routine δ-scale measurements’ (Gröning, 2011), isotope values from reference materials and laboratory standards δ^13^C values were drift-corrected on a daily (batch-by-batch) basis. Drift correction of isotope values was carried out before data normalisation. Post drift correction, sample δ^13^C values were three-point normalised using isotopic data from the daily analysis of reference materials; IAEA 603 Calcite (2.46 ± 0.01 ‰; 2SE), IAEA 611 Carbonate (-30.795 ± 0.04 ‰; 2SE), and NBS19 Limestone (-1.95 ‰) (Coplen et al., 2006). Values are reported relative to Vienna Pee Dee Belemnite (VPDB). Precision and accuracy were assessed through repeat analyses of IAEA NBS18 Calcite standard. Historical measurements show internal precision is 0.01–0.06 ‰ for δ^13^C and external reproducibility is 0.02‰ for δ^13^C relative to VPDB (all errors given as 2SE). For experimental sessions related to this study (Supplementary Material, Table S3), the mean δ^13^C value in the IAEA NBS18 Calcite standard (-5.01 ± 0.06 ‰, n = 16) was within error of the IAEA recommended value (-5.014 ± 0.035‰) (Coplen et al., 2006).

Due to the use of fossil fuels, there is a 1.2 permille decrease in δ^13^C values in modern samples compared to pre-industrial. Therefore, we apply a correction of -1.2% to all macropodids so comparisons can be made between fossils and modern datasets (Friedli et al., 1986, Montanari et al., 2013). In addition, as there is a ∼12 ‰ fractionation in δ^13^C values between marsupial diet and enamel (Montanari et al., 2013, Fraser et al., 2008, Murphy et al., 2007), a correction of -12‰ was applied to convert measured δ^13^C values to dietary δ^13^C values. The percentage of C_3_ and C_4_ plant intake was calculated following methodology outlined in Montanari et al. (2013).

### 3.6 Strontium isoscape of central eastern Queensland

A strontium isoscape for the region surrounding Mount Etna Caves was generated following methodology described in Bataille et al. (2020). ^87^Sr/^86^Sr isotope ratios measured in vegetation specimens were compiled into a pre-existing global database of bioavailable Sr isotope ratios (Bataille et al., 2020). Measured ^87^Sr/^86^Sr isotope ratios were then integrated into a spatially-explicit machine learning model with a range of 28 covariates compiled in Bataille et al. (2020) that have the potential to influence ^87^Sr/^86^Sr isotope ratios including geology, soil composition, vegetation types, and topographical features. Using the coordinates provided in the Sr isotope ratio database, environmental covariate values (e.g., geology, soil composition) were extracted for each location. These covariates were then analysed using the *VSURF* package (Variable Selection Using Random Forest) (Genuer et al., 2010) to identify the most important predictors of bioavailable ^87^Sr/^86^Sr (Supplementary Fig. 4, Bataille et al., 2020). Using the *caret* package (Kuhn, 2008), and framework presented in Bataille et al. (2020) the Sr dataset was used to train a random forest regression, incorporating these aforementioned environmental predictors. Briefly, random forest is a machine-learning algorithm that creates an ensemble of decision trees through bootstrap sampling and random feature selection to improve the accuracy of predictions (Bataille et al. 2020). The model’s predictions were validated using a subset of the data not included in the training phase ensuring the model accurately captured the spatial variability of ^87^Sr/^86^Sr ratios. Once validated, the model was used to generate continuous spatial predictions of ^87^Sr/^86^Sr isotope ratios across the Mount Etna region.

### 3.7 Geographic assignment of fossil specimens

Estimates of foraging range were determined by combining measured ^87^Sr/^86^Sr isotope ratios in fossilised enamel Sr isoscapes using the *assignR*, package described in (Ma et al., 2020). For each individual specimen, a ‘regional’ and ‘narrow’ probability map was produced using a ‘regional’ isoscape encompassing 10 000 km^2^ area around Mount Etna Caves, and a smaller 1 000 km^2^ defined as ‘narrow’. This approach enables the prediction of geographic origin and range of individual specimens during enamel mineralisation.

Using the *assignR* package, joint probability maps were created grouping individuals from the same taxon with similar individual probability maps. Considering site chronology, stratigraphic units examined here represent three distinct periods of time; >330 ka (QML1384LU), 280 – 330 ka (QML1311H and QML1311C/D), and 180 – 205 ka (QML1312) (Hocknull, 2005b, Hocknull et al., 2007, Laurikainen Gaete et al., 2025). Using these three distinct periods in time, a series of joint probability maps were created, grouping individuals from the same taxon with a similar depositional age and geographic origin. This was done to examine changes in foraging range, across a period of relatively stable rainforest environment and then in response to broader scale environmental shifts.

## 4. Results

### 4.1 Strontium isotope variation in macropodids fossil enamel

The smallest macropodid individuals, represented by *Thylogale*, return a mean ^87^Sr/^86^Sr ratio of 0.70888 ± 0.00021 (n = 12) with values ranging from 0.70774 ± 0.00003 to 0.70914 ± 0.00013. No statistical differences were found between stratigraphic units (Kruskal-Wallis, p = 0.24) with a mean value of 0.70878 ± 0.00035 (n = 3) in QML1311C/D; 0.07844 ± 0.00041 (n = 4) in QML1311H; and 0.70874 ± 0.00033 (n = 5) in QML1384LU (Fig. 1; Table 1).

**Figure 1.**
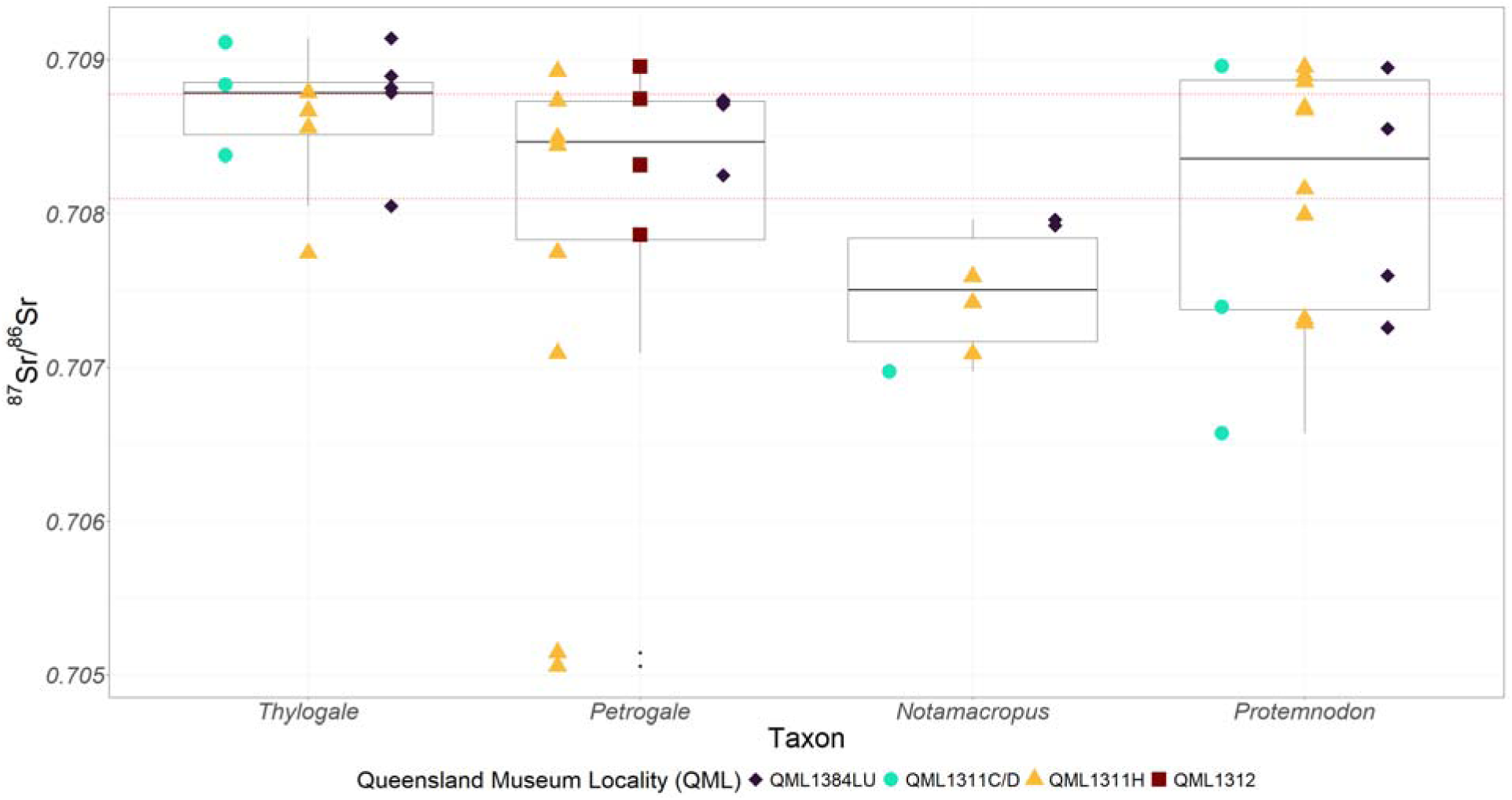
^87^Sr/^86^Sr ratios measured in fossil enamel. Isotope ratios are compared to the range of values for the local rock formation Mount Etna Beds Limestone reported in Deer (2011) (red dashed lines).

**Table 1.** ^87^Sr/^86^Sr ratios measured in fossil enamel.

| Sample Name | Genus | Sample Type | Tooth | $^{87}\text{Sr}/^{86}\text{Sr}$ | 2SE |
| --- | --- | --- | --- | --- | --- |
| QML1311C/D_Mac1 | *Notamacropus* | Molar | M3 | 0.706974 | 0.000062 |
| QML1311H_Mac2a | *Notamacropus* | Molar | M3 (Base) | 0.707418 | 0.000098 |
| QML1311H_Mac2b | *Notamacropus* | Molar | M3 (Top) | 0.707089 | 0.000042 |
| QML1311H_Mac3 | *Notamacropus* | Molar | M2 | 0.707590 | 0.000030 |
| QML1384LU_Mac4 | *Notamacropus* | Molar | M2 | 0.707962 | 0.000036 |
| QML1384LU_Mac5 | *Notamacropus* |  |  | 0.707923 | 0.000079 |
| QML1311H_Pet1a | *Petrogale* | Incisor | I1 (Base) | 0.708488 | 0.000055 |
| QML1311H_Pet1b | *Petrogale* | Incisor | I1 (Middle) | 0.708442 | 0.000055 |
| QML1311H_Pet2 | *Petrogale* | Molar | M3 | 0.708924 | 0.000058 |
| QML1311H_Pet3a | *Petrogale* | Molar | M3 (Base) | 0.707745 | 0.000087 |
| QML1311H_Pet3b | *Petrogale* | Molar | M3 (Top) | 0.707093 | 0.000050 |
| QML1311H_Pet4 | *Petrogale* | Incisor | I1 | 0.708730 | 0.000065 |
| QML1311H_Pet5a | *Petrogale* | Molar | M4 (Base) | 0.705145 | 0.000028 |
| QML1311H_Pet5b | *Petrogale* | Molar | M4 (Top) | 0.705058 | 0.000026 |
| QML1384LU_Pet6 | *Petrogale* | Incisor | I1 | 0.708250 | 0.000088 |
| QML1384LU_Pet7 | *Petrogale* | Incisor | I1 | 0.708739 | 0.000047 |
| QML1384LU_Pet8a | *Petrogale* | Incisor | I1 | 0.708708 | 0.000075 |
| QML1384LU_Pet8b | *Petrogale* | Incisor | I1 | 0.708724 | 0.000071 |
| QML1312_Pet9 | *Petrogale* | Molar | M2-M4 | 0.707862 | 0.000049 |
| QML1312_Pet10 | *Petrogale* | Molar | M2-3 | 0.708956 | 0.000054 |
| QML1312_Pet11 | *Petrogale* | Molar | M3-4 | 0.708746 | 0.000049 |
| QML1312_Pet12 | *Petrogale* | Molar | M1-M3 | 0.708314 | 0.000060 |
| QML1311H_Pro1 | *Protemnodon* | Molar | M4 | 0.708160 | 0.000020 |
| QML1311H_Pro2 | *Protemnodon* | Molar | m4 | 0.707992 | 0.000041 |
| QML1311H_Pro3 | *Protemnodon* | Molar | M/4 | 0.707313 | 0.000045 |
| QML1311H_Pro4 | *Protemnodon* | Molar | M2/ | 0.707288 | 0.000051 |
| QML1384LU_Pro5 | *Protemnodon* | Molar | M4/ | 0.707597 | 0.000039 |
| QML1384LU_Pro6 | *Protemnodon* | Molar | M2/ | 0.707258 | 0.000019 |
| QML1311C/D_Pro7 | *Protemnodon* | Incisor | i1 | 0.707393 | 0.000023 |
| QML1311C/D_Pro8 | *Protemnodon* | Molar | m4 | 0.706570 | 0.000018 |
| QML1311C/D_Pro9 | *Protemnodon* | Molar | M4/ | 0.708958 | 0.000095 |
| QML1311H_Pro10a | *Protemnodon* | Molar | m2 (Base) | 0.708858 | 0.000068 |
| QML1311H_Pro10b | *Protemnodon* | Molar | m2 (Top) | 0.708671 | 0.000054 |
| QML1311H_Pro11 | *Protemnodon* | Molar | m2 or m3 | 0.708952 | 0.000046 |
| QML1311H_Pro12a | *Protemnodon* | Molar | M4 (Base) | 0.708686 | 0.000047 |
| QML1311H_Pro12b | *Protemnodon* | Molar | M4 (Top) | 0.708894 | 0.000060 |
| QML1384LU_Pro13a | *Protemnodon* | Molar | M3/ (Base) | 0.708550 | 0.000041 |
| QML1384LU_Pro13b | *Protemnodon* | Molar | M3/ (Top) | 0.708948 | 0.000086 |
| QML1311C/D_Thy1 | *Thylogale* | Molar | m/3 (Base) | 0.708378 | 0.000102 |
| QML1311C/D_Thy2a | *Thylogale* | Molar | m/2 | 0.708838 | 0.000077 |
| QML1311C/D_Thy2b | *Thylogale* | Molar | m/3 | 0.709112 | 0.000072 |
| QML1311H_Thy3 | *Thylogale* | Molar | m/3 | 0.708558 | 0.000067 |
| QML1311H_Thy4 | *Thylogale* | Molar | m/3 | 0.707741 | 0.000033 |
| QML1311H_Thy5 | *Thylogale* | Molar | M4/ | 0.708667 | 0.000122 |
| QML1311H_Thy6 | *Thylogale* | Molar | M2/ | 0.708785 | 0.000031 |
| QML1384LU_Thy7 | *Thylogale* | Molar | M2/ | 0.708895 | 0.000077 |
| QML1384LU_Thy8a | *Thylogale* | Incisor | i/1 (Base) | 0.708786 | 0.000034 |
| QML1384LU_Thy8b | *Thylogale* | Incisor | i/1 (Top) | 0.708817 | 0.000057 |
| QML1384LU_Thy9 | *Thylogale* | Molar | M4/ | 0.708049 | 0.000088 |
| QML1384LU_Thy10 | *Thylogale* | Incisor | i/1 | 0.709139 | 0.000128 |

*Petrogale,* returned a mean ^87^Sr/^86^Sr isotope ratio of 0.70800 ± 0.00060 (n = 16), with no significant differences compared to *Thylogale* (Fig. 1; Table 1; Kruskal-Wallis, p = 0.17). Isotope ratios range from 0.7056 ± 0.00003 to 0.70896 ± 0.00005 with two samples measured in one individual from QML1311H returning the lowest Sr isotope ratios of any sample. This individual lowered the mean ^87^Sr/^86^Sr ratio to 0.70745 ± 0.00103 (n = 8) in QML1311H contrasting the remaining stratigraphic units: 0.70861 ± 0.00021 (n = 4) in QML1384LU; 0.70847 ± 0.00042 (n = 4) in QML1312. However, no significant differences were found between stratigraphic units (Kruskal-Wallis, p = 0.34).

*Notamacropus* exhibited the lowest mean ^87^Sr/^86^Sr ratio across all taxa, at 0.70749 ± 0.00031 (n = 6), significantly lower than both *Petrogale* and *Thylogale* (Kruskal-Wallis, p = 0.01). Isotope ratios ranged from 0.70697 ± 0.00006 to 0.70792 ± 0.00008 with no significant differences between stratigraphic units (Kruskal-Wallis, p = 0.12).

The largest macropodid individuals, *Protemnodon*, showed no significant differences in Sr isotope composition (mean = 0.70813 ± 0.00038, n = 16) compared to *Thylogale, Petrogale*, or *Notamacropus* (Kruskal-Wallis, p = 0.27, p = 0.16, and p = 0.08, respectively). Except for one *Petrogale* outlier, *Protemnodon* exhibited the greatest variation in ^87^Sr/^86^Sr ratios, ranging from 0.70657 ± 0.00002 to 0.70896 ± 0.00010 (Fig. 1). Isotope ratios varied slightly across stratigraphic units: 0.70764 ± 0.00114 (n = 3) in QML1311C/D, 0.70831 ± 0.0041 (n = 9) in QML1311H, and 0.70809 ± 0.00069 (n = 4) in QML1348LU, however differences are not statistically significant (Kruskal-Wallis, p = 0.78).

Overall, ^87^Sr/^86^Sr ratios showed no significant differences between *Thylogale, Petrogale*, and *Protemnodon* (Kruskal-Wallis, p = 0.23), but were significantly lower in *Notamacropus* (Kruskal-Wallis, p = 0.03). Variation across macropodids suggests individuals occurred across distinct geological substrates (Fig. 1; Table 1), highlighting the need for an Sr isoscape to predict past geographic ranges accurately.

### 4.3 Carbon isotope variation within macropodids

Measured δ^13^C values across all macropodid samples ranged from -13.54 ‰ to -4.14 ‰ indicating diet ranging from C_3_ specialist feeders, C_3_/C_4_ mixed feeders, and one C_4_ specialist (Fig. 2; Table 2). Due to the smaller sample size, detailed comparisons between genera and stratigraphic units were not possible, and most results are interpreted qualitatively.

**Figure 2.**
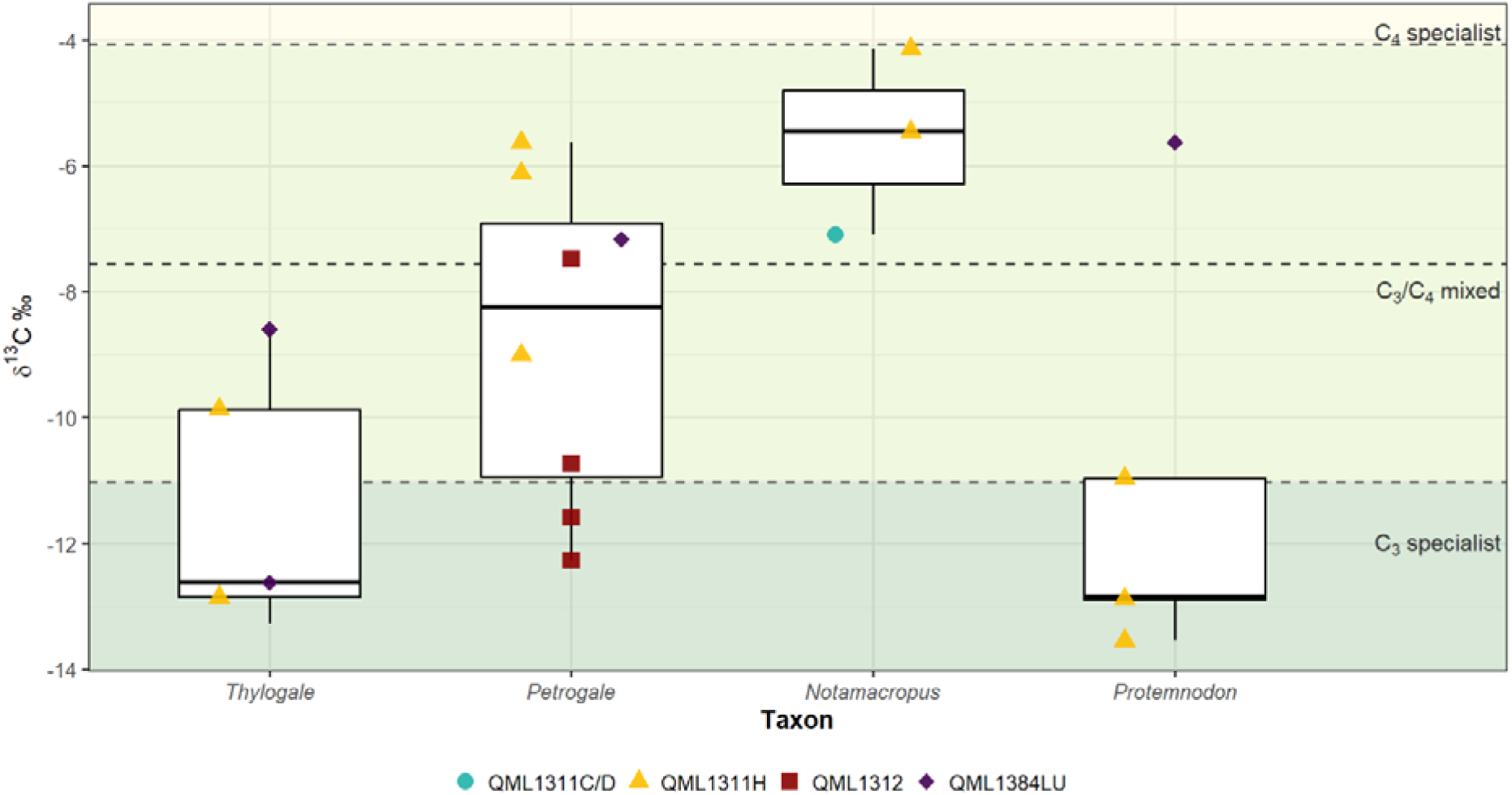
Carbon isotope (δ^13^C) values (Corrected for the Suess Effect) measured in fossil enamel. Dashed grey lines indicate dietary thresholds with diets comprising >75% C_4_ vegetation are classified as ‘C_4_ specialists’, while those with >75% C_3_ vegetation are classified as ‘C_3_ specialists’ black dashed line represents a true ‘mixed feeder’ with C_3_ and C_4_ vegetation both comprising 50% of dietary intake. Proportional intake of C_3_ and C_4_ vegetation (%) used in dietary thresholds is calculated based on Johnson et al. (1997).

**Table 2.** Dietary δ^13^C values, and proportional intake of C_3_ and C_4_ vegetation in macropodid specimens.

| Sample Name | Genera | Sample Type | $\delta^{13}\text{C}$ Measured (Suess Effect) | $\delta^{13}\text{C}$ Diet | Proportion $\text{C}_3$ (%) | Proportion $\text{C}_4$ (%) | Dietary Preference Classification |
| --- | --- | --- | --- | --- | --- | --- | --- |
| QML1311C/D_Mac1 | *Notamacropus* | Molar | -7.10 | -19.10 | 47% | 53% | $\text{C}_3/\text{C}_4$ mixed |
| QML1311H_Mac2 | *Notamacropus* | Molar | -4.14 | -16.14 | 25% | 75% | $\text{C}_4$ specialist |
| QML1311H_Mac4 | *Notamacropus* | Molar | -5.46 | -17.46 | 35% | 65% | $\text{C}_4$ preference |
| QML1311H_Pet2 | *Petrogale* | Molar | -9.01 | -21.01 | 61% | 39% | $\text{C}_3/\text{C}_4$ mixed |
| QML1311H_Pet3a | *Petrogale* | Molar | -6.12 | -18.12 | 40% | 60% | $\text{C}_3/\text{C}_4$ mixed |
| QML1311H_Pet5 | *Petrogale* | Molar | -5.63 | -17.63 | 36% | 64% | $\text{C}_3/\text{C}_4$ mixed |
| QML1384LU_Pet7 | *Petrogale* | Incisor | -7.17 | -19.17 | 47% | 53% | $\text{C}_3/\text{C}_4$ mixed |
| QML1312_Pet9 | *Petrogale* | Molar | -10.73 | -22.73 | 73% | 27% | $\text{C}_3$ preference |
| QML1312_Pet10 | *Petrogale* | Molar | -7.48 | -19.48 | 49% | 51% | $\text{C}_3/\text{C}_4$ mixed |
| QML1312_Pet11 | *Petrogale* | Molar | -11.58 | -23.58 | 79% | 21% | $\text{C}_3$ specialist |
| QML1312_Pet12 | *Petrogale* | Molar | -12.27 | -24.27 | 84% | 16% | $\text{C}_3$ specialist |
| QML1311H_Pro1 | *Protemnodon* | Molar | -10.97 | -22.97 | 75% | 25% | $\text{C}_3$ specialist |
| QML1311H_Pro2 | *Protemnodon* | Molar | -12.88 | -24.88 | 88% | 12% | $\text{C}_3$ specialist |
| QML1384LU_Pro6 | *Protemnodon* | Molar | -5.63 | -17.63 | 36% | 64% | $\text{C}_3/\text{C}_4$ mixed |
| QMI1311H_Pro10 | *Protemnodon* | Molar | -12.85 | -24.85 | 88% | 12% | $\text{C}_3$ specialist |
| QML1311H_Pro11 | *Protemnodon* | Molar | -13.54 | -25.54 | 93% | 7% | $\text{C}_3$ specialist |
| QMI1311C/D_Thy1 | *Thylogale* | Molar | -13.27 | -25.27 | 91% | 9% | $\text{C}_3$ specialist |
| QML1311H_Thy5 | *Thylogale* | Molar | -9.87 | -21.87 | 67% | 33% | $\text{C}_3$ preference |
| QML1311H_Thy6 | *Thylogale* | Molar | -12.85 | -24.85 | 88% | 12% | $\text{C}_3$ specialist |
| QML1384LU_Thy7 | *Thylogale* | Molar | -8.60 | -20.60 | 58% | 42% | $\text{C}_3/\text{C}_4$ mixed |
| QML1384LU_Thy9 | *Thylogale* | Molar | -12.62 | -24.62 | 86% | 14% | $\text{C}_3$ specialist |

$\delta^{13}\text{C}$ diet values are obtained by taking measured $\delta^{13}\text{C}$ values in enamel (Corrected for the Suess Effect) and subtracting the diet-enamel enrichment factor of 12 ‰ (Montanari et al., 2013, Fraser et al., 2008, Murphy et al., 2007). Proportional intake of $\text{C}_3$ and $\text{C}_4$ vegetation (%) is calculated based on Johnson et al. (1997).

*Thylogale* returned a mean δ^13^C value of -11.44 ± 1.86 (n = 5) indicating a strong preference for C_3_ vegetation (78 ± 13% of dietary intake). Values ranged from -13.27 ‰ to - 8.60 ‰ with three individuals considered C_3_ specialists (-12.92 ± 0.38 ‰, 89 ± 2% of dietary intake), and a mixed C_3_/C_4_ diet observed in two individuals (-9.23 ± 1.27 ‰, 62 ± 9% of dietary intake).

*Petrogale* individuals returned a mean δ^13^C value of -9.68 ± 1.80 ‰ (n = 8) constituting a mixed C_3_/C_4_ intake with a minor C_3_ preference (65 ± 13 % of dietary intake). Values range from -12.27 ‰ to -5.63 ‰. A higher mean δ^13^C value (-6.98 ± 1.50 ‰; n = 4) is observed in individuals from the older deposits – QML1311H and QML1384LU – compared to those from the younger, xeric-adapted fauna of QML1312 ( -10.51 ± 2.12 ‰; n = 4) (Welch two sample t-test, p =0.04). This suggests the older individuals may have consumed a higher proportion of C_4_ vegetation (54 ± 3% of dietary intake) compared to the younger individuals (C_3_ forming 71 ± 16% of dietary intake), however, this difference warrants future fine-scale study.

*Notamacropus* individuals have the highest mean δ^13^C value (-5.57 ± 1.71 ‰, n = 3), indicating a diet comprised predominately of C_4_ plants (64 ± 12 % of dietary intake). Values range from -7.10 ‰ to -4.14 ‰ with C_4_ specialisation observed in one individual, but more generalist trends in another individual with C_4_ vegetation only forming 53% of total dietary intake (Table 2).

*Protemnodon* had a mean δ^13^C value of -11.18 ± 2.90 ‰ (n = 5), indicating a strong C_3_ preference (76 ± 21 % of total dietary intake). Inter-sample variation was high, with values ranging from -13.54 ‰ to -4.43 ‰. While four specimens were C_3_ specialists (87 ± 8% C_3_), one individual showed a preference for C_4_ vegetation (64%).

### 4.4 Strontium isoscape of central-eastern Queensland

Using the random forest regression model, continuous spatial predictions of ^87^Sr/^86^Sr isotope ratios were undertaken for the broader Mount Etna region (Fig. 3). Spatial uncertainty for the strontium isoscape was assessed based on the relationship between absolute residual values and predicted bio-available ^87^Sr/^86^Sr (Fig. 3, (Bataille et al., 2020)). The spatial distribution of ^87^Sr/^86^Sr isotope ratios is strongly correlated with lithology with the highest ^87^Sr/^86^Sr isotope ratios observed in the north of the isoscape, in an area underlain with Princhester with values exceeding 0.7016, but a greater uncertainty due to the lack of bioavailable Sr data (Fig. 3). High Sr isotope ratios are also observed in localised Neoproterozoic to Palaeozoic serpentine units to the north-west (> 0.7111) and east (> 0.710) of Mount Etna Caves, although these are lower than those predicted for Princhester Serpentinite.

**Figure 3.**
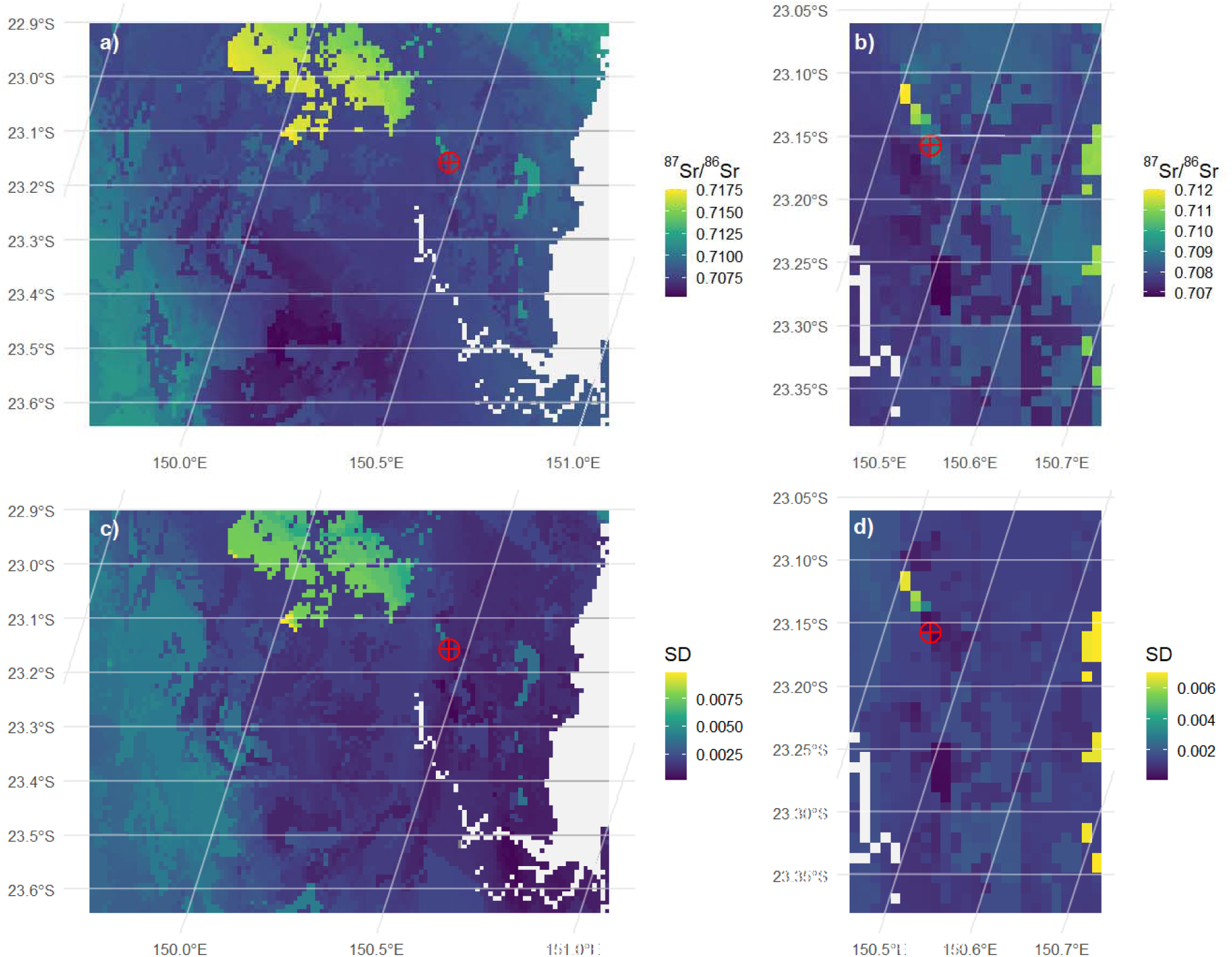
(a) ‘Regional’ and (b) ‘Narrow’ 87Sr/86Sr isoscape. (c) associated error for the ‘regional’ and (d) 472 ‘narrow’ isoscape. In this an following figures, Mount Etna Caves is denoted by a red circle, and x and y axes show longitude and latitude in a WGS84 Web Merca projection. field of view is 115 x 90 km in the ‘regional’ and 25 x 40km in the ‘narrow’ Sr isoscape. White pixels represent the Fitzroy the Pacific Ocean.

Lower ^87^Sr/^86^Sr isotope ratios (< 0.708) are observed ∼40 km west of the Fitzroy River (Fig. 3a), underlain by Native Cat Andesite, and Dalma Basalt, as well as in a small patch (<10 km) south of Mount Etna Caves, underlain by a Late Permian to Early Triassic Gabbroid (Fig. 3).

To explore potential foraging ranges, we generated isoscapes at two spatial scales: a larger 8,000 km² region covering the total distribution of plant samples (‘regional’; Fig. 3a) and a smaller 900 km² region around the caves (‘narrow’; Fig. 3c). Both the strontium isoscape and the geo-provenance package have been made publicly available (Supplementary Material) providing a useful tool for future provenance studies of both modern and fossil material from the wider Rockhampton region.

### 4.5 Geographic assignment of fossil specimens

#### 4.5.1 Thylogale

Individual probability maps for 10 *Thylogale* specimens both the ‘regional’ and ‘narrow’ isoscape show the highest probabilities within a 9 km radius of Mount Etna Caves (Supplementary Figs. 5 and 6). Joint probability maps grouped by stratigraphic unit display similar foraging ranges across time, with minor differences attributed to sample size (Supplementary Fig. 7). For two individuals with multiple aliquots analysed show no intra-individual variation indicating that foraging ranges remained consistent during enamel mineralisation. These small discrete home ranges have been characterised as ‘local-north’ (n = 4) with high probabilities adjacent to (< 1 km), and <9 km northwest of Mount Etna Caves (Fig. 4).

**Figure 4.**
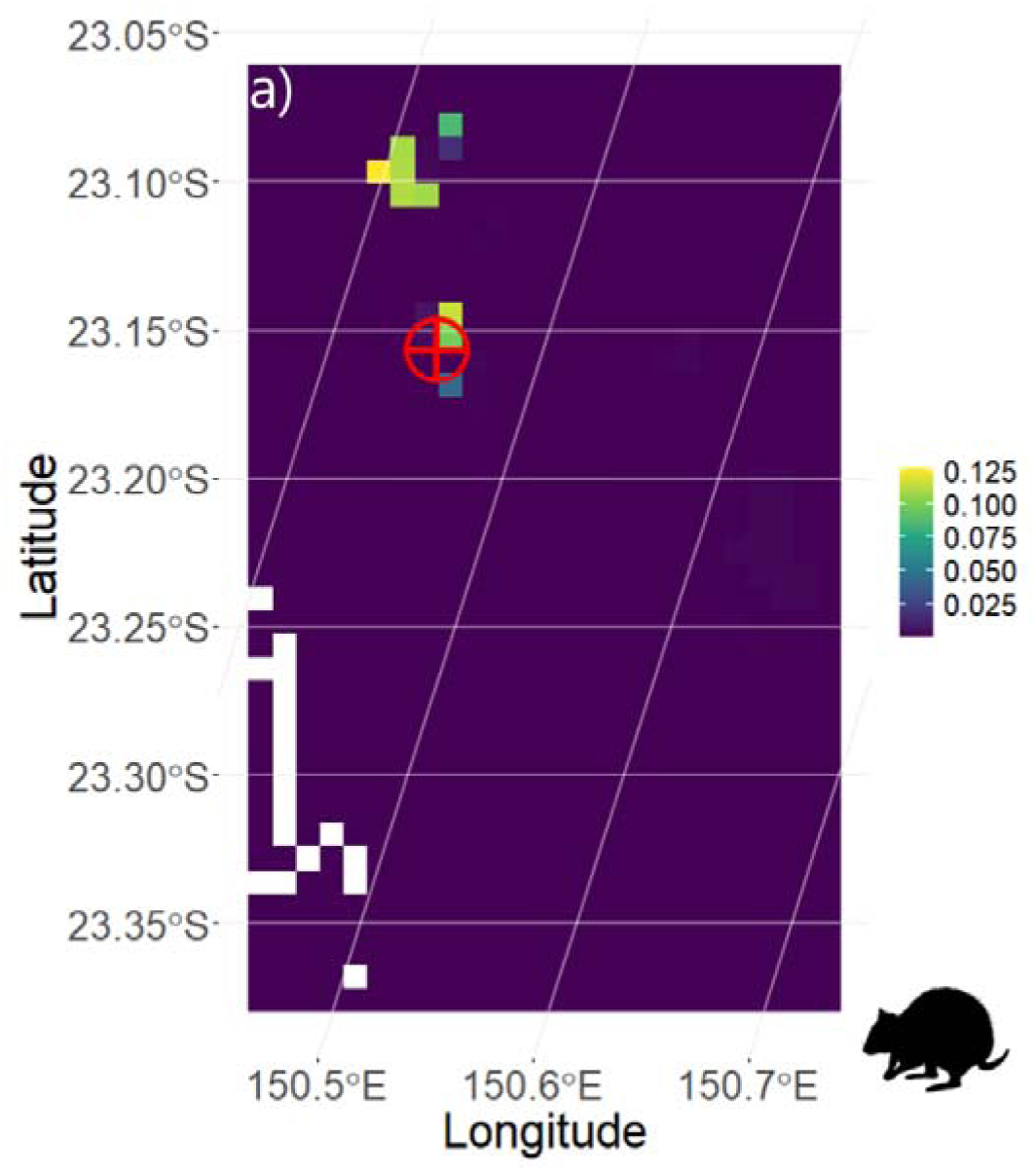
Joint probability map for ten *Thylogale* specimens with showing a high probability adjacent to, and <9 km north-west of the fossil bearing caves. In this and following figures, brighter colours denote higher probability values (as shown on the right-hand scale). Red circle is Mount Etna Caves. White pixels represent Fitzroy River. probability within a 6 km radius of Mount Etna Caves, using the ‘narrow’ Sr isoscape, and termed ‘local (north)’ (see text for details).

#### 4.5.2 Petrogale

For *Petrogale*, individual probability maps for most individuals using the ‘regional’ isoscape show the highest probabilities directly adjacent to Mount Etna Caves and toward the mouth of the Fitzroy River (Supplementary Fig. 8). Whilst similar patterns are seen in most individuals, QML1311H_Pet3b differs, foraging 15 km south (Supplementary Fig. 8). Two aliquots from QML1311H_Pet5(a & b) show a distinct foraging range >60 km southwest of Mount Etna Caves, west of the Fitzroy River (Supplementary Fig. 8). Using the ‘narrow’ isoscape reveals four main foraging regions: (1) nine aliquots showing a hyper-localised signal within a 1 km radius of Mount Etna Caves, (2) four individuals occurring <9 km northwest of Mount Etna Caves, (3) four *Petrogale* specimens with a local to southern origin still within 5 km, and (4) one individual ∼15 km south (Supplementary Fig. 9). Due to these distinct ranges, a joint probability map for all *Petrogale* individuals doesn’t yield a clear point of origin. Therefore, we treat the groups independently using both the ‘regional’ and ‘narrow’ isoscape (Fig. 5a–e).

**Figure 5.**
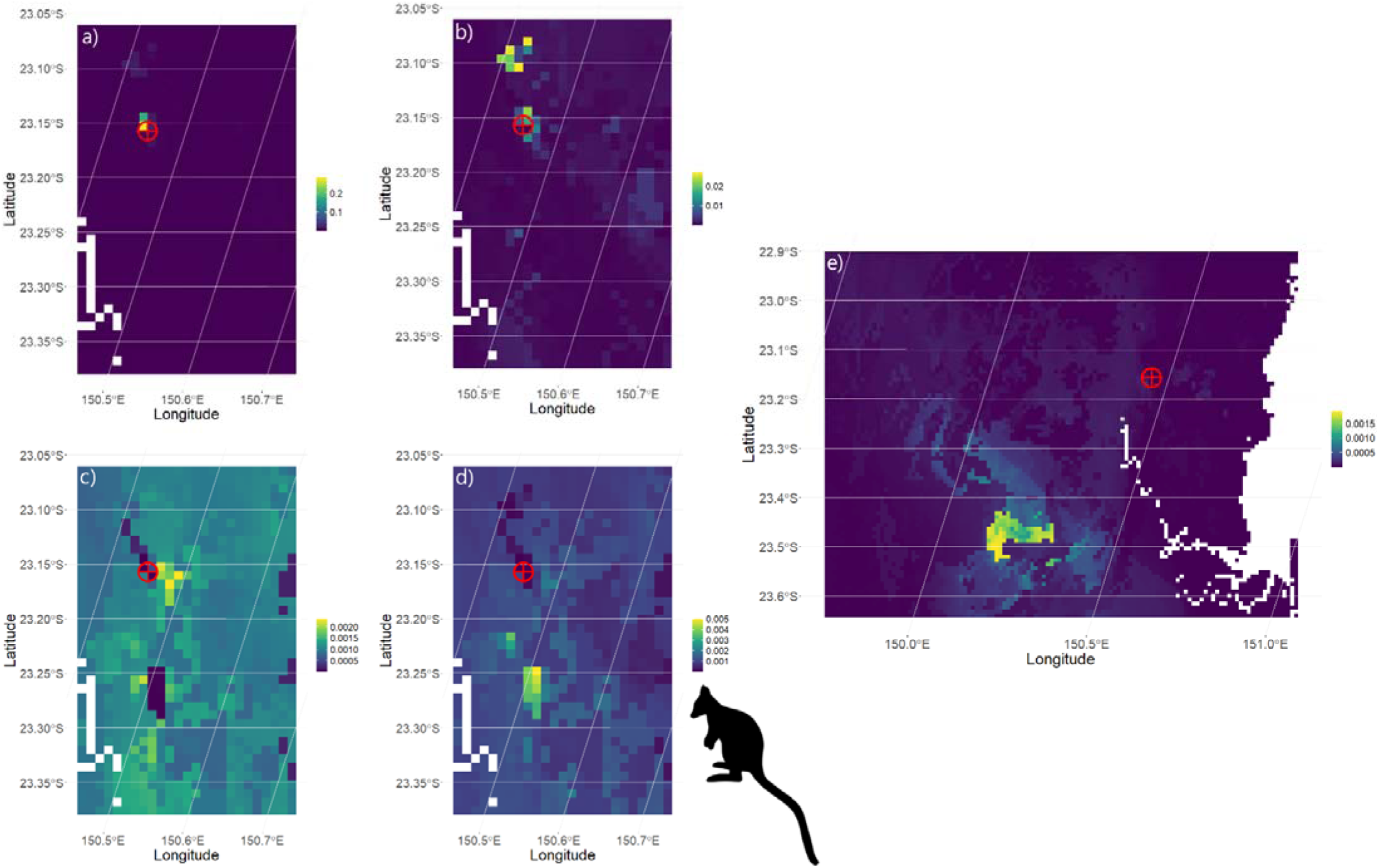
(a) Joint probability map for nine *Petrogale* specimens with showing a higher probability within a 1 km radius of Mount Etna Caves, using the ‘narrow’ Sr isoscape, and termed ‘local’. (b) Joint probability map for four *Petrogale* specimens with showing a higher probability <9 km to the north of Mount Etna Caves, using the ‘narrow’ Sr isoscape, and termed ‘local (north)’. (c) Joint probability map for four *Petrogale* specimens with showing a higher probability to the south (<5 km) of Mount Etna Caves, using the ‘narrow’ Sr isoscape, and termed ‘local (south)’. (d) Probability map for one of two aliquots of a *Petrogale* specimen (QML1311H_Pet3b), showing higher probability 15 km south of the fossil deposits, using the ‘narrow’ Sr isoscape, and termed thereafter ‘southern’. (e) joint probability map for two aliquots from an individual specimen (QML1311H_Pet5a & b), showing a higher probability >60 km south-west of the fossil deposits, using the ‘regional’ Sr isoscape, and termed thereafter ‘foreign’. Red circle is Mount Etna Caves. White pixels represent Fitzroy River.

For hyper-localised individuals (henceforth ‘local’; n = 9), the joint probability shows high probabilities <1 km from the fossil site (Fig. 5a). For individuals with a local to northern signature (n = 4), the joint probability map mirrors that of *Thylogale* (Fig. 4), and as such, these individuals are also classified as ‘local (north)’ (Fig. 5b). A joint probability map four individuals residing directly south of Mount Etna Caves (Fig. 5c) shows they occupied a distinctly different area when compared to ‘local (north)’ *Petrogale* and *Thylogale* individuals and are henceforth termed ‘local (south)’. Aliquot ‘b’ of QML1311H_Pet3, from the molar base, suggests a high probability ∼15 km south of Mount Etna (Fig. 4b), while aliquot ‘a’, from the tooth crown, indicates a ‘local (north)’ origin (Fig. 4a). This suggests a shift in foraging range during enamel mineralisation, with the individual originating at Mount Etna before travelling south, thus indicating connectivity of viable habitat between these regions.

For QML1311H_Pet5, two repeat aliquots show the highest probability > 60 km south-southwest of Mount Etna Caves, near Mount Morgan (Fig. 4c). This suggests the individual foraged near Mount Morgan before dispersing to Mount Etna Caves. Given the vast distance, post-mortem transport by a predator seems unlikely, with no know predators exhibiting collecting behaviour over such an immense distance, even for very large raptorial predators. Therefore, until viable taphonomic agents are available, results indicate this ‘foreign’ individual dispersed >60 km following enamel mineralisation of the M4 molar.

Joint probability maps by fossil age indicate that the local foraging ranges remained consistent across all three time periods (Supplementary Fig. 10). However, the distinct ‘southern’ and ‘foreign’ foraging ranges were only observed in fossils from QML1311H, dating between 280–330 ka (Laurikainen Gaete et al., 2025). We propose that these unique ranges pre-280ka are indicative of suitable habitat connectivity for *Petrogale* between the local Mount Etna population with habitats 15 km to the south and potentially as far as 60 km southwest. A broader sample set is needed to confirm this hypothesis and examine whether such connectivity persisted beyond 280 ka.

#### 4.5.3 Notamacropus

For *Notamacropus*, using the ‘regional’ Sr isoscape, four individuals show diffuse probability maps, with the highest probabilities near Mount Etna Caves (Supplementary Fig. S11). The remaining two individuals show a distinct origin south of Mount Etna Caves (Supplementary Fig. S11). The ‘narrow’ Sr isoscape further supports this, with four individuals foraging within a 5 km radius of Mount Etna (Supplementary Fig. S12). A joint probability map for these four individuals indicates that they resided near the caves (Fig. 6a), in a similar habitat occupied by ‘local (south)’ Petrogale (Fig. 5a). The two remaining individuals indicate a ‘southern’ foraging range (Fig. 6b), overlapping with the region occupied by the ‘southern’ *Petrogale* individual (Fig. 5d).

**Figure 6.**
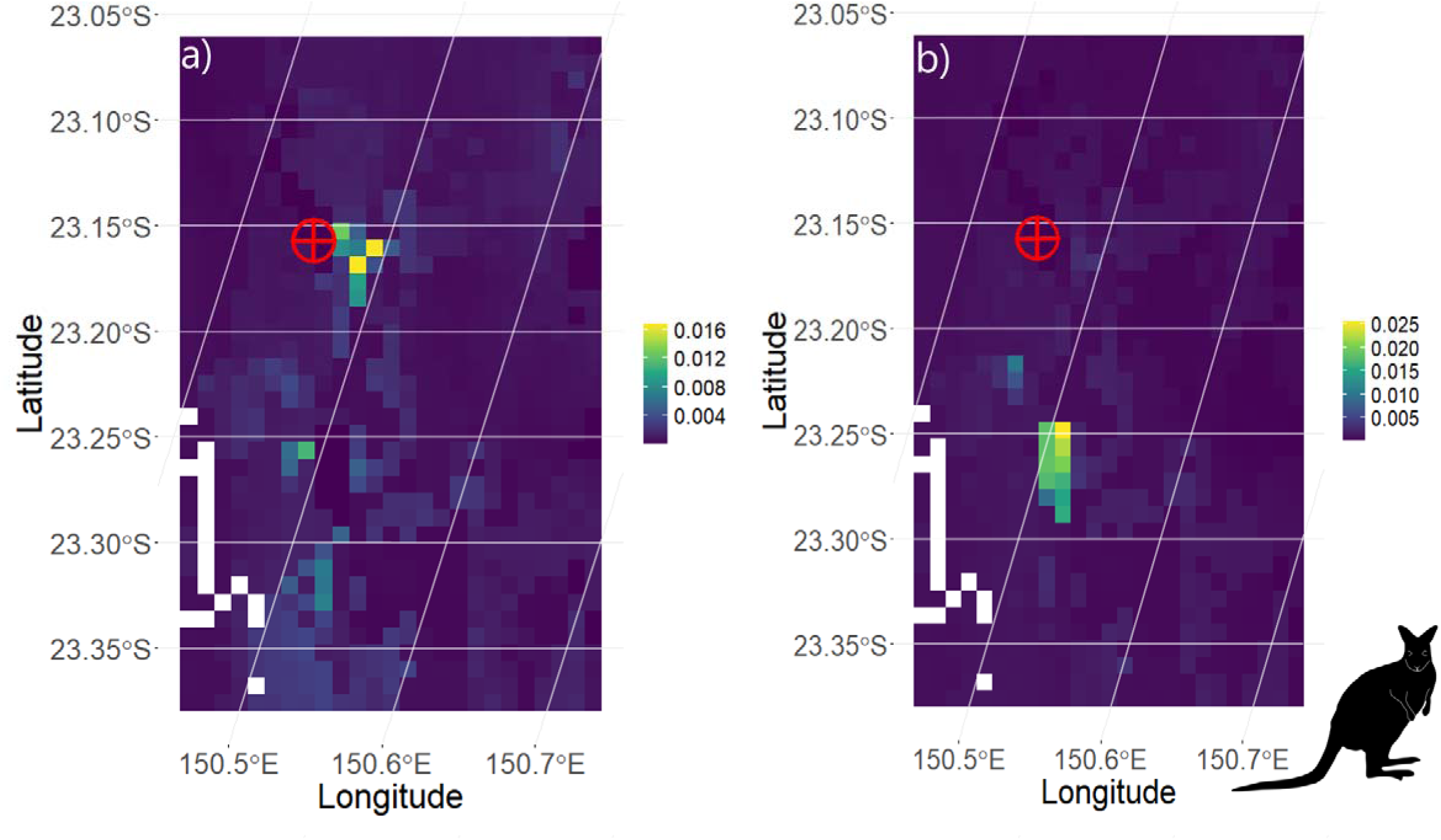
(a) Joint probability map for four *Notamacropus* specimens with individual probability maps showing a foraging area <5 km south of the caves termed thereafter ‘local (south)’ (b) Joint probability map for two *Notamacropus* specimens with individual probability maps showing a foraging area ∼15 km south of the caves termed thereafter ‘southern’. Both probability maps are derived using the ‘narrow’ Sr isoscape. Red circle is Mount Etna Caves. White pixels represent Fitzroy River.

For QML1311H_Mac2, an aliquot from the base of the tooth (QML1311H_Mac2a) indicates a ‘southern’ foraging range, whereas the crown (QML1311H_Mac2b) reveals a ‘local (south)’ signature (Supplementary Fig. 12). This suggests early-life dispersal from the ‘local (south)’ habitat to a ‘southern’ foraging region, a similar, but counter-directional movement to that described *Petrogale*, QML1311H_Pet3. Regarding fossil age, the two *Notamacropus* individuals from the oldest deposit, QML1384LU (> 330ka), nearly identical probability maps, both classified as ‘local (south)’ (Supplementary Fig. 13). While this ‘local (south)’ foraging range is still present in younger deposits (QML1311C/D and QML1311H; 280 – 330 ka), two individuals also show a distinct ‘southern’ foraging range (Supplementary Fig. S13).

#### 4.5.4 Protemnodon

For *Protemnodon*, individual probability maps of eight specimens using the ‘regional’ isoscape show relatively diffuse probabilities, with the highest probabilities occurring near Mount Etna Caves (Supplementary Fig. 14). Using the ‘narrow’ Sr isoscape individual probability maps indicate a ‘local’ to ‘local (north)’origin (Supplementary Fig. 15). A joint probability map for these eight specimens (Fig. 7a), supports this ‘local (north)’ mirroring ‘local (north)’ *Thylogale* and *Petrogale* individuals (Figs. 4 and 5b). Four specimens indicate a foraging area around 9 km south-west of the caves, along the Fitzroy River (Supplementary Fig. 15), slightly further afield when compared to ‘local (south)’ members of *Petrogale* and *Notamacropus* (Figs. 5c and 6a). One individual, QML1311C/D_Pro8 shows a unique foraging range, 15 km south of Mount Etna Caves (Fig. 6c), directly overlapping the ‘southern’ range seen in both *Petrogale* and *Notamacropus* (Figs. 5d and 6b).

**Figure 7.**
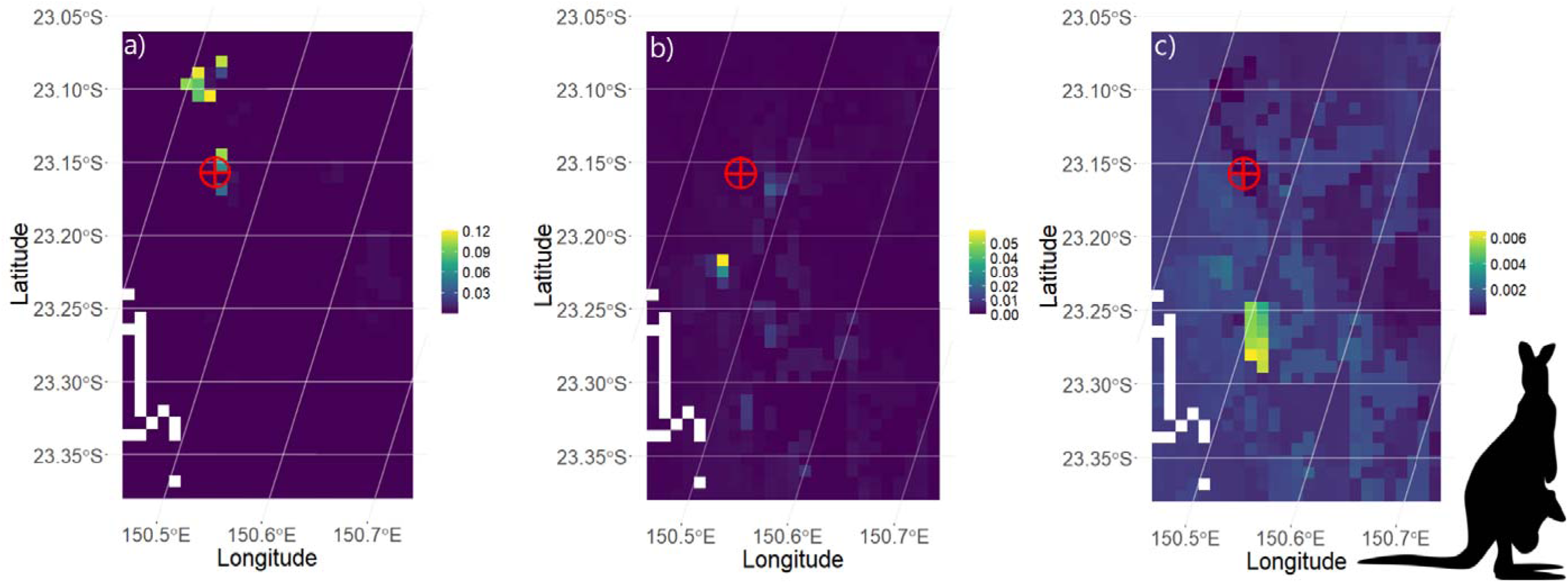
(a) Joint probability map for 8 *Protemnodon* individuals with individual probability maps showing a foraging area <9 km from of the caves termed thereafter ‘local (north)’. (b) joint probability map for 4 *Protemnodon* individuals with individual probability maps showing a foraging area <9 km south-west now included in the ‘local (south)’ classification. (c) Probability map for one *Protemnodon* specimen (QML1311H_Pro8), showing high probability 15 km south of the fossil deposits termed thereafter ‘southern’. All probability maps are derived using the ‘narrow’ Sr isoscape. Red circle is Mount Etna Caves. White pixels represent Fitzroy River.

Chrono-stratigraphically, ‘local (north)’ and ‘local (south)’ foraging ranges are observed in the oldest stratigraphic unit QML1384LU (> 330 ka, Supplementary Fig. 16). These ranges persist in younger specimens (QML1311H and QML1311C/D; 280–330 ka), with one additional individual exhibiting a distinct ‘southern’ foraging range (Supplementary Fig. 16).

#### 4.5.5 Geographic assignment summary

Overall, similar trends in foraging ranges were observed across individual macropodids, regardless of taxon or body mass (Figs. 4–7). All individuals of *Thylogale*, and most individuals of *Petrogale* (n = 11), *Notamacropus* (n = 4), and *Protemnodon* (n = 8) were classified as ‘local’, foraging within a 6 km radius of the caves (Fig. 8). A distinct northwest signal, termed ‘local (north)’, was observed for individuals of *Thylogale, Petrogale*, and *Protemnodon*, excluding *Notamacropus*, while a ‘local (south)’ to ‘southern’ signal was found for individuals of *Protemnodon, Petrogale*, and *Notamacropus*, excluding *Thylogale* (Fig. 8). We acknowledge potential limitations due to sample size and note that future studies are needed to further investigate this north-to-south division at Mount Etna Caves during the Middle Pleistocene, modulated by the presence or absence of *Thylogale* and *Notamacropus*.

**Figure 8.**
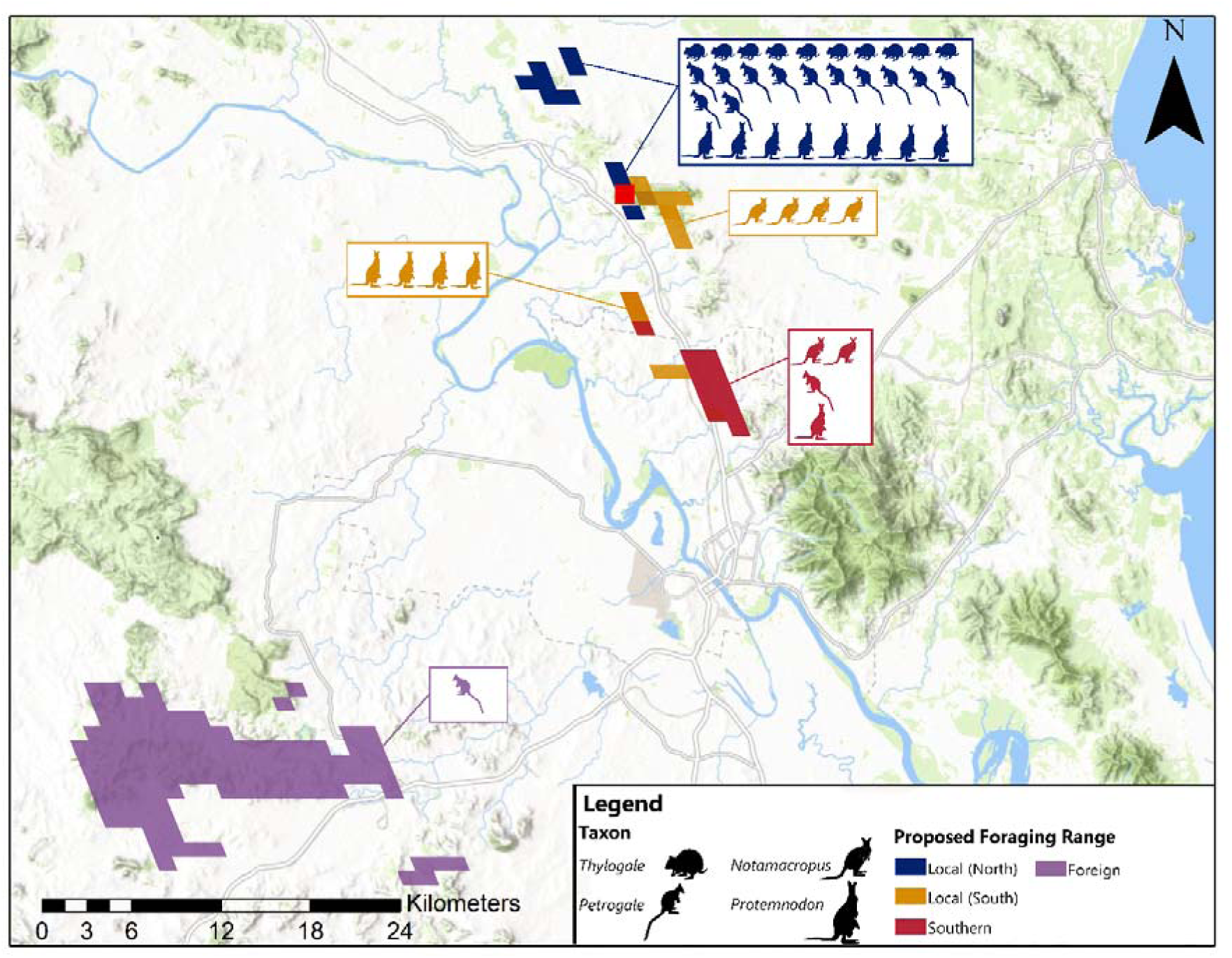
Simplified regional map showing the key categorical foraging ranges proposed for individual macropodids from Mount Etna Caves. ‘Local’ macropodids have been separated into distinct ‘local (North)’, including ‘local’ *Petrogale* and other ‘local (north)’ individuals residing <9 km northwest of fossil deposits (n = 30) and ‘local (south)’ residing <9 km south to south-west of fossil deposits (n = 8). ‘Southern’ individuals are characterised by a unique range 15 km south (n = 4). The ‘foreign’ individual is characterised by a distinct range 60 south-west of Mount Etna (n = 1). Field of view is 80 x 60 km.

### 4.6 Combining isotopic proxies

Considering that foraging range and diet are interrelated, dietary preferences were examined for each taxon across the distinct foraging ranges described above (Fig. 8).While all *Thylogale* were characterised by a ‘local (north)’ foraging range, a divide in δ¹³C values was observed between C_3_ specialists (over 75% of dietary intake, n = 3) and mixed C_3_/C_4_ individuals with a C_3_ preference (C_3_ > 50%, n = 2) (Fig. 2).

In *Petrogale*, there is a clear distinction in both dietary preference and foraging behaviour in younger individuals from QML1312, and older members from QML1311H and QML1384LU. In QML1312*, Petrogale* exhibit ‘local (north)’ foraging range and C_3_/C_4_ dietary plasticity, ranging from C_3_ specialisation to C_3_/C_4_ mixed feeding, but with a tendency for higher C_3_ inclusion (Table 2; Fig. 2). Older individuals are characterised by broader foraging ranges (Fig. 8), but a similar degree of dietary plasticity – with marginally higher C_4_ inclusions in some individuals (Fig. 9).

**Figure 9.**
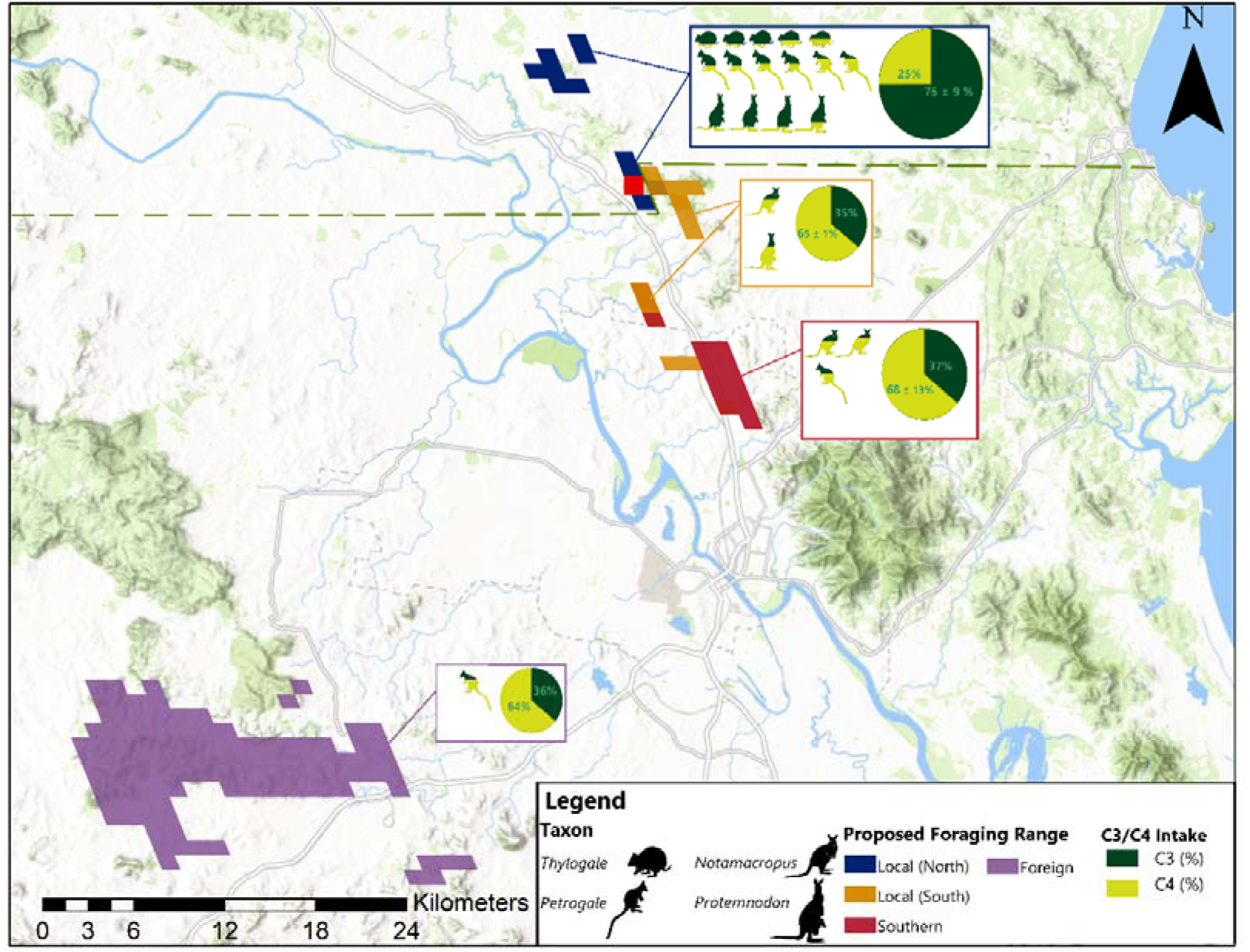
Simplified regional map showing the key categorical foraging ranges (defined in Fig. 8) and C_3_/C_4_ preferences for individual macropodids (Table 2). For each distinct foraging range, the mean proportional C_3_/C_4_ intake is depicted with a pie chart.

*Notamacropus* shows distinct foraging range and δ¹³C values compared to most other macropodids with a clear ‘local (south)’ to ‘southern’ foraging range (Fig. 8) and a stronger preference for C_4_ vegetation relative to other macropodids. This provides further evidence of a distinct transition in vegetation communities to the south of Mount Etna (Fig. 9).

Most *Protemnodon* are ‘local (north)’ C_3_ specialists; however, one individual with a ‘local (south)’ foraging range showed a stronger preference for C_4_ vegetation, resembling other ‘local (south)’ and ‘southern’ members of *Petrogale* and *Notamacropus* (Fig. 9).

Overall, a division in dietary preferences is evident between ‘local (north)’ and ‘local (south)’ to ‘southern’ individuals, indicating that Mount Etna was situated at the junction of two ecotones with varying C_3_/C_4_ vegetation compositions (Fig. 9). A north-western population, consisting of specialist to preferential C_3_ consumers and a south and southwest community composed of mixed feeders with greater C_4_ inclusions (Fig. 9).

## 5. Discussion

### 5.1 Thylogale

Our results showed a relatively restricted total foraging range for *Thylogale* individuals, confined to a 1 km radius with a potential maximum of 9 km from Mount Etna (Fig. 8). Such restricted foraging ranges are consistent with modern *Thylogale* species. Although *T. billardierii, T. stigmatica,* and *T. thetis* exhibit temporal range shifts between closed forests and pastoral lands (Le Mar and Mcarthur, 2005, Vernes et al., 1995, Johnson, 1980), overall landscape use is minimal (Vernes et al., 1995, Johnson, 1980).

When considering C isotope data at the genus level, an overall mixed C_3_/C_4_ intake is consistent with the observed diet of modern *Thylogale* species (Rose and Rose, 2018, Johnson and Vernes, 1994, Vernes et al., 1995, Smith et al., 2022) suggesting that the total dietary breadth has remained unchanged since the Middle Pleistocene. Observed dietary δ¹³C values (Table 2) are not as negative as those expected for closed-canopy understory vegetation in the modern Wet Tropics (-35.5 ± 1.7‰; Cheesman et al., 2020). Given the faunal composition indicative of closed rainforest (Hocknull, 2009, Hocknull, 2003, Cramb et al., 2023, Cramb et al., 2020, Cramb and Hocknull, 2010a, Cramb et al., 2009), these isotope values suggest that *Thylogale* were foraging along the margins of these closed-canopy rainforest patches, in habitats comparable to SEVT still observed in the region today (Neldner, 2023). This interpretation aligns with the edge-foraging behaviour seen in modern *Thylogale* species (Le Mar and Mcarthur, 2005, Vernes et al., 1995, Johnson, 1980) indicating a consistent foraging strategy since the Middle Pleistocene.

Variable C_3_/C_4_ dietary intake appears to be linked to body mass, with larger-bodied individuals – comparable in size to *T. thetis* – operating as mixed C_3_/C_4_ feeders, while smaller-bodied individuals – comparable to *T. stigmatica or T. billardierii* – showing stronger C_3_ preferences (Tables 2, Supplementary Table S1). In modern sympatric populations, *T*.

*stigmatica* and *T. thetis* exhibit temporal, spatial and dietary separation focusing on rainforest browse (i.e. C_3_ preference) and graze (C_3_/C_4_ consumer) respectively (Smith et al., 2022). A recent study *Thylogale* biomechanics supports this species-specific niche differentiation suggesting clear morphological differences in the crania of *T. stigmatica stigmatica*, *T. stigmatica wilcoxi* and, *T. thetis* were directly related to the proportional intake of graze and browse (Mitchell et al., 2020). Hocknull (2005b), (2009) identified two species of *Thylogale* from the Middle Pleistocene deposits at Mount Etna Caves, a species comparable in size to *T. stigmatica* and an exceptionally small, undescribed new species. Only individuals of the larger *Thylogale* were selected for this study, and while they were assumed to have been conspecific with *T. stigmatica*, measurements fall within the size range of *T. thetis*, indicating the possibility of a third species of *Thylogale* associated with the caves. While additional studies of cranial and post-cranial remains from Mount Etna Caves are now required to identify species-level delimitations in *Thylogale*, dietary divergence could indicate the presence of three, sympatric species of *Thylogale* at Mount Etna Caves during the Middle Pleistocene.

Therefore, for taxa with very similar molar morphology, such as *Thylogale,* combined Sr and C isotopes may provide an independent line of evidence for species-level delimitation. These independent and objective isotopic signals may present valuable tools for taxonomic allocation of fossil taxa with highly fragmentary but morphologically similar dentition.

### 5.2 Petrogale

In modern *Petrogale*, reliance on fixed diurnal shelters significantly limits the extent of foraging ranges (Laws and Goldizen, 2003, Telfer and Griffiths, 2006). Studies of *P. assimilis* (Horsup, 1994), *P. brachyotis* (Telfer and Griffiths, 2006), and *P. penicillata* (Laws and Goldizen, 2003) suggest foraging ranges of 0.02 – 0.19 km^2^. The hyper-restricted (< 1 km) ranges observed in most *Petrogale* individuals from Mount Etna (Fig. 5a) likely reflect similar constraints, with individuals centred around rocky refugia associated the caves.

While most individuals were either hyper-localised (< 1 km), ‘local (north)’, or ‘local (south)’ foraging range, one specimen exhibited variable Sr composition, indicating a 15 km journey south during the mineralisation of an M3 molar. Although modern species of *Petrogale* are understood to be highly localised, moderate-scale male-biased dispersal (∼8 km) has been observed between colonies of *P. lateralis* (Eldridge et al., 2001), and *P. penicillata* (Hazlitt et al., 2006). Both local and ‘southern’ habitats (Fig. 8) are occupied by modern *P. inornata* (Supplementary Fig. 17), residing in remnant SEVT (Supplementary Fig. 18). This suggests connectivity between these two habitats at least until 280 ka. A larger sample from QML1312 and genetic analyses of modern *P. inornata* would be valuable in reconstructing the past connection-disconnection of metapopulations determining at what point in time dispersal between *Petrogale* colonies ceased, and whether this had an impact on genetic diversity and survivorship of these two groups.

One *Petrogale* specimen displayed a large-scale dispersal of over 60 km, originating near Mount Morgan (Fig. 8). While such movements have not been observed in extant species, Eldridge et al. (2001) noted low genetic differentiation between populations of *P. brachyotis* separated by >67 km, may reflect male-biased dispersal across undisturbed habitat. Interestingly, this large-scale dispersal indicates a degree of inter-specific interconnectivity between *P. inornata* at Mount Etna and *P. herberti* to the west of the Fitzroy River (Supplementary Figure S17). Although *P. inornata* and *P. herberti* populations abut at the Fitzroy River (Mitchell et al., 2024), phylogenetic analysis shows these species diverged around 2 million years ago, long before the Middle Pleistocene deposits at Mount Etna (Eldridge et al., 1991, Eldridge and Close, 1992, Barker and Close, 1990). This ‘foreign’ isotope signature may indicate multiple *Petrogale* lineages at Mount Etna during the Middle Pleistocene. Although presently inconclusive, the foreign individual *Petrogale* is larger in size, compared to the other *Petrogale* specimens, suggesting either a male-biased dispersal, or a different species may be present (Supplementary Table 1). Future work examining more dental and post-cranial remains of *Petrogale* is now crucial to (1) determine whether multiple species of *Petrogale* can be identified within Mount Etna’s fossil deposits, and (2) the relative abundance of these species through time and their ranges.

Considering modern barriers to dispersal, Smith et al. (2023) suggested that the construction of an exclusion fence in The Grey Ranges, Queensland provided a significant barrier for dispersal, effectively isolating modern colonies of *P. xanthopus celeris* separated by ∼5 km. At Mount Etna, current results indicate the presence of highly vagile individuals during the Middle Pleistocene, however, a small sample set beyond 280 ka means we are unable to conclusively determine at what point in time this dispersal ceased. This lack of knowledge highlights a need for future studies utilising a broader set of *Petrogale* individuals from the late Pleistocene through to the present day to conclusively determine when this larger-scale individual dispersal ceased, and whether this can be attributed to a changing climate, or human impacts. From a conservation management point of view, understanding these historic corridors for dispersal and determining what external factors contributed to their loss is crucial for (1) maintaining interconnectivity between modern meta-populations where dispersal is still reported (Smith et al., 2023, Eldridge et al., 2001, Hazlitt et al., 2006), and (2) re-establishing connections between isolated *Petrogale* colonies to avoid help combat genetic erosion (Smith et al., 2023) and facilitate greater survivorship of threatened populations.

At Mount Etna, *Petrogale* diet broadly reflects a generalist consumption of C_3_/C_4_ vegetation, mirroring dietary plasticity observed in multiple modern lineages (Horsup, 1994, Roth, 2015, Tuft et al., 2011). In modern taxa, significant dietary plasticity ensures *Petrogale* can thrive in geographically constrained areas, directly adjacent to their diurnal shelter (Laws and Goldizen, 2003, Telfer and Griffiths, 2006). Greater C_4_ inclusions in local individuals – relative to *Thylogale* and *Protemnodon* – indicates an ability to access locally available C_4_ resources not available to specialist browsing taxa. Overall, a mixed diet consisting of both C_3_ and C_4_ vegetation with spatio-temporal variability indicate that broader foraging envelopes and an ability to respond to variable resource availability were already well-established behavioural traits by the Middle Pleistocene.

### 5.3 Notamacropus

*Notamacropus* from Mount Etna are characterised by small, discrete foraging ranges an affinity for C_4_ vegetation (Fig. 9). Considering known grazing specialisations in *Notamacropus* (Arman et al., 2025), C_4_ preferences likely reflect an abundance of graze resources in these distinct ‘local (south)’ and ‘southern’ ranges (Fig. 9). Overall results for *Notamacropus* mirror extant behaviours in members of *N. agilis* in the Northern Territory (Stirrat, 2002, Stirrat, 2003, Stirrat, 2004) suggesting, behaviour has remained relatively unchanged since the Middle Pleistocene.

### 5.4 Protemnodon

Most *Protemnodon* were characterised by small foraging ranges (<9 km) comparable to those observed in individuals of *Thylogale* and *Petrogale* (Fig. 8). Small foraging ranges mirror those reported in marsupial megafauna taxa, *Protemnodon*, *Procoptodon,* and *Diprotodon* from Wellington Caves (Koutamanis et al., 2023) but contrast two-way latitudinal migrations up to 200 km observed in *Diprotodon* from the Darling Downs (Price et al., 2017a). Koutamanis et al. (2023) proposed that small ranges reflect a highly productive environment where marsupials could meet all dietary needs within a limited area. Preliminary results from Mount Etna *Protemnodon* in Laurikainen Gaete et al. (2025) further support this argument attributing limited ranges to (1) an abundance of local resources (Bengsen et al., 2016, Herfindal et al., 2005, Relyea et al., 2000, Willems et al., 2009, Fisher and Owens, 2000) and/or (2) unique, lower-geared, quadrupedal movements in some species of *Protemnodon* (Jones et al., 2022, Den Boer, 2018, Kerr, 2023, Janis et al., 2020, Kerr et al., 2024). An aggregation of macropodids with strong C_3_ preferences directly adjacent to and north-west of Mount Etna Caves indicates a likely abundance of C_3_ vegetation (i.e., browse) within a relatively small geographic area (Fig. 9). This may indicate that, unlike placental mammals, macropodid foraging ranges in the Middle Pleistocene may have been more closely linked to environmental productivity. However, further studies in less productive environments are needed to determine whether these small ranges were solely a result of environmental conditions or a broader dispersal limit across the Macropodidae.

A strong C_3_ preference in ‘local (north)’ *Protemnodon* aligns with previous carbon isotope findings from *Protemnodon* at Chinchilla Sands (Montanari et al., 2013), and Cuddie Springs (DeSantis et al., 2017), with these browsing inferences further supported by skeletal morphology, dental structure and microwear analyses (DeSantis et al., 2017, Montanari et al., 2013, Hocknull, 2009). In contrast, the ‘local (south)’ population of *Protemnodon* showed dietary divergence, with a greater reliance on C_4_ grasses (Table 2, Fig. 9). Grazing is not unexpected, as Kerr et al. (2024) suggested a range of dietary niches across Pliocene and Pleistocene species of *Protemnodon*, with obligate grazing observed in *P. mamkurra* (Arman et al., 2025). We propose that this dietary diversification may either reflect significant dietary plasticity across *Protemnodon* individuals from Mount Etna Caves, or more likely, dietary niche diversification across distinct species of *Protemnodon*.

Multiple Australian fossil deposits show evidence of inter-specific *Protemnodon* co-existence (Kerr et al., 2024). Although species-level identification from Mount Etna are not definitive, three size groups are evident, smaller individuals (n = 5) comparable in size to *P. otibandus* and *P. tumbuna* (Kerr et al., 2024); an intermediate size morph (n = 5) comparable to *P. anak* (Kerr et al., 2024), and the largest morph (n = 3) comparable in size to *P. viator* (Kerr et al. (2024)). While the ‘medium’ and ‘large’ morphs share similar foraging ranges and C_3_ preferences, the smaller *Protemnodon* morph displayed a distinct grazing niche and a separate foraging range to the south-west of Mount Etna Caves (Fig. 9, Supplementary Fig. 19). Although further analysis of dental, cranial, and post-cranial fossils is necessary for species-level delimitations, these geographic, dietary, and morphological differences suggest at least two distinct *Protemnodon* populations associated with Mount Etna Caves during the Middle Pleistocene: a larger, browsing species to the north-west and a smaller, grazing species to the south. As with *Thylogale*, Sr and C isotopes provide a powerful tool for identifying palaeoecological niches, indicating greater levels of species-level diversity than what might be drawn on the basis of morphology alone.

### 5.5 Broader Community Assemblages

Considering the broader community assemblages, trends across taxa indicate a clear geographic divide centred on Mount Etna with at least two distinct ecotones occurring within a relatively small area (Fig. 9). The north-western region comprised of at least two *Thylogale* species, one *Petrogale* species, and a large, browsing *Protemnodon* species. In contrast, the southern ecotone hosts a *Notamacropus* species, one *Petrogale* species, and a smaller, grazing *Protemnodon*.

These ecotones appear linked to underlying geological features, with north-western, C_3_ habitat associated with the Mount Alma Formation and Mount Alma Formation limestone. Limestone karst formations often support high species diversity, including endemic plant, vertebrate, and invertebrate communities (Struebig et al., 2009, Schilthuizen et al., 2005, Clements et al., 2008, Clements et al., 2006). Similar influences are seen in Brazilian neotropical regions, where high fertility from limestone outcrops formed ‘litho-refugia,’ supporting seasonally dry tropical forests (SDTF) (de Aguiar-Campos et al., 2020). In the present day, the Mount Etna Caves limestone formations are characterised by remnant patches of semi-evergreen vine thicket (SEVT) or “dry rainforest” habitat (Martinez, 2010, Sprent and Sprent, 1970, Neldner, 2023). This strong association between SEVT and underlying lithology may indicate that during the Middle Pleistocene, the Mount Etna region was characterised by mosaic habitats with larger areas of continuous “dry rainforest” associated with the Mount Alma Formation limestone, and more open C_4_ dominated habitats associated with the Rockhampton Group.

However, the Mount Etna Caves deposits are characterised by a significant diversity of vertebrate fauna indicative of closed-forest conditions, including several arboreal mammalian lineages, *Dendrolagus, Pseudochirulus*, and *Phalanger* (Hocknull et al., 2007, Hocknull, 2009), high diversity of dasyurids (Cramb et al., 2009, Cramb and Hocknull, 2010b, Cramb et al., 2023), rainforest-restricted rodents (Cramb et al., 2020, Hocknull et al., 2007, Hocknull, 2005b), and plesiomorphic rainforest koalas (Price and Hocknull, 2011). SEVT or “dry rainforest” alone would be unable to support this species-level diversity, indicating the presence of a closed-canopy rainforest comparable to the Queensland Wet Tropics or New Guinea highlands (Hocknull et al., 2007).

In global context, understory vegetation in Amazon and Congo closed-canopy rainforests is characterised by exceptionally low δ¹³C values with a threshold of < –31.5‰ proposed in Kohn (2010). If this threshold were applicable in the Australian context, we would expect closed-canopy rainforest macropodids to exhibit dietary δ¹³C values approaching –31.5‰. However, this is not reflected in our fossil dataset with a mean dietary δ¹³C values of –24.54 ± 1.33‰ our ‘local (north)’ C_3_ specialists (Table 2), significantly higher than the closed-canopy threshold of –31.5‰ (Kohn, 2010). This discrepancy highlights the need to consider regional variations in δ¹³C values, as global thresholds may not directly apply to Australopapuan rainforest systems. This is supported by a national carbon isoscape which indicates a mean vegetation δ¹³C value of –26.97 ± 1.82‰ in the Wet Tropics (Munroe et al., 2022). This aligns closely with our fossil C_3_ specialists and suggests they may have occupied habitats comparable to modern closed Australopapuan rainforests. However, a localised study in the Daintree rainforest found a pronounced closed-canopy effect with δ¹³C values as low as –35.5 ± 1.7‰ in understory vegetation compared to –29.9 ± 1.0‰ in canopy vegetation (Cheesman et al., 2020).

These discrepancies in δ¹³C values across modern, Australian closed-forest habitats, complicates direct comparisons with fossil data, particularly when interpreting the habitat preferences of C_3_-specialist macropodids. Considering observed behavioural trends in modern *Thylogale*, discussed in more detail in section 5.1, individuals often forage at rainforest margins, transitional zones between closed-forest and more open pastural lands (Le Mar and Mcarthur, 2005, Vernes et al., 1995, Johnson, 1980). Similar geographic assignment in ‘local’ *Protemnodon* and *Petrogale* (Fig. 9) suggests that these macropodid with a C_3_ preference may have also resided in this transitional zone, potentially comparable to remnant SEVT observed in the landscape today (Neldner, 2023). This interpretation is supported by modern observations of *Petrogale penicillata*, residing in rocky escarpments bordering naturally fragmented vegetation assemblages or transition zones with a mixture of vegetation types including eucalypt woodland, dry rainforest, open cattle pastures, and sclerophyll forest (Murray et al., 2008, Laws and Goldizen, 2003). These findings support the interpretation of a structurally complex mosaic landscape, with at least two, but potentially three distinct habitat zones: (1) open C_4_ grasslands to the south, (2) transitional SEVT-like margins supporting C_3_ macropodids, and (3) closed-canopy rainforest interiors likely occupied by arboreal taxa not assessed in this study.

Rainforest-specific arboreal taxa such, *Dendrolagus, Pseudochirulus, Phalanger*, and rainforest-restricted rodents are well represented in the Mount Etna Caves fossil assemblage (Hocknull et al., 2007, Cramb et al., 2023, Cramb et al., 2020, Cramb and Hocknull, 2010b, Cramb et al., 2009), yet their isotopic profiles remain unknown. Future Sr and C isotope analyses of these closed-forest taxa are essential to refine interpretations of habitat partitioning within this mosaic rainforest system. If they show significantly lower δ¹³C values, then it is likely our macropodids described here are residing on the fringes of a closed-forest habitat, as observed in their modern counterparts. If however, δ¹³C values are comparable to our C_3_ specialist macropodids, this may indicate a less pronounced canopy effect in Australian rainforests (Munroe et al., 2022), and that our macropodids were in fact foraging in a closed-forest environment.

Irrespective of specific habitat allocations, a local mosaic of habitats, linked to underlying geology presents major implications for interpreting past environments based on fossil assemblages alone. On the basis of stratigraphic association, the co-occurrence of multiple species may represent sympatric distributions (Lyman, 2016). However, Sr and C isotope data challenge this notion, suggesting not all macropodids at Mount Etna Caves were truly sympatric. Rather, a mosaic environment with distinct, C_3_ dominant and open C_4_ ecotones providing distinct habitat patches which allowed multiple, similar mammalian lineages to co-exist with minimal interspecific interactions in a limited overall geographic area.

### 5.6 Extinction and Survivorship Patterns

Limited mobility may have been beneficial in climatically stable environments; however, we hypothesise that intensifying aridity and shifts to xeric conditions post-280 ka caused the disappearance of intervening habitats for low-mobility taxa. By the Middle Pleistocene, macropodids had evolved specialised feeding, biomechanics, and biogeographic distributions, but the capacity to rapidly adapt to increased aridity may have been insufficient, leading to the extinction of low-mobility taxa with niche dietary preferences. This is evident at Mount Etna with the localised disappearance of non-vagile C_3_ specialists (*Protemnodon* and *Thylogale*) indicating they lacked the dietary flexibility and biomechanical capacity for large-scale movement needed in a more arid environment. This mirrors size-biased extinction patterns observed in megafaunal lemurs (Crowley and Godfrey, 2019), supporting the idea that both the smallest and largest individuals are more vulnerable to habitat fragmentation and localised extinction(Haskell et al., 2002).

In contrast, the surviving taxa – *Petrogale* and *Notamacropus* – exhibit broad dietary envelopes and potential for moderate to large-scale dispersal, suggesting generalist behaviours already established by the Middle Pleistocene meant these species were better suited to respond to environmental shifts. This mobility likely allowed them to persist in increasingly arid conditions by moving between refugial patches as environmental conditions deteriorated through the Pleistocene (Hocknull et al., 2020).

Overall, these extinction and survivorship patterns suggest that for macropodids, dietary plasticity and dispersal ability were key to survival during periods of major environmental change. Understanding the baseline life history traits and the intimate connection between patterns of past to present distribution, along with dietary preference and capacity to adapt to environmental changes is crucial for effective management and conservation of modern species. In particular, our research demonstrates the importance of understanding the long-term distributions of habitat corridors so that dispersal between key core ranges can be maintained.

## 6. Conclusions

In this study, Sr isotopes in macropodid enamel and a Sr isoscape were used to reconstruct past distribution and foraging ranges of several Middle Pleistocene macropodine taxa from Mount Etna Caves. While a linear regression model predicted larger ranges with increasing body mass, the actual foraging ranges of most macropodids highly localised. Small ranges in *Thylogale, Petrogale*, and *Notamacropus* align with modern species, suggesting ranges have remained unchanged since the Middle Pleistocene. For *Protemnodon*, small foraging ranges corroborate findings from Bingara and Wellington Caves (Koutamanis et al., 2023) indicating that macropodid range extent may be more strongly linked to habitat quality than body mass (Fisher and Owens, 2000). While most macropodids were non-vagile, some *Petrogale* individuals exhibited high mobility, suggesting that the isolation in modern *Petrogale* populations is likely due to habitat fragmentation, and the loss of lowland dispersal corridors. Reintroducing these corridors could allow highly mobile individuals to move between metapopulations and promote gene flow. Dietary preferences in *Thylogale, Petrogale*, and *Notamacropus* align with modern species, indicating that macropodid dietary envelopes were already well established by the Middle Pleistocene. For *Protemnodon*, we propose that dietary, geographic, and morphological separation may reflect the presence of at least two different species at Mount Etna. Similarly, in *Thylogale*, distinct dietary preferences correlated with body mass suggest the presence of two species, emphasising the potential of Sr and C isotopes to distinguish species-level differences, particularly when taxonomic classification is restricted to isolated teeth. Combining Sr and C isotopes reveal two distinct ecotones, a C_3_-dominated region to the northwest, and a C_4_-dominated grassland to the south. This suggests that throughout the middle Pleistocene, a highly productive mosaic of habitats driven by underlying geology and climatic stability supported two distinctly different macropodid communities within a small geographic area. However, increasing aridity post 280 ka disrupted this equilibrium with the localised extinction of non-vagile, dietary specialists and the cessation of moderate to large scale dispersal of highly vagile individuals driven by the loss of lowland habitats.

## 7. Acknowledgements

We acknowledge the Traditional Custodians of the Mount Etna region, the Darumbul people, and pay our respect to past, present, and emerging leaders. We also wish to thank Noel Sands, Emily Bradstock, and staff from Capricorn Caves, for their field and collecting assistance. We thank Cement Australia, the Central Queensland Speleological Society, Queensland Parks and Wildlife and Capricorn Caves for their assistance during fossil collection at Mount Etna.

Fieldwork was undertaken at Mount Etna with permission from and in collaboration with Cement Australia, and in collaboration with Queensland Parks and Wildlife Service (Scientific Permit WITK17469216, Permit to Take, Use, Keep or Interfere P-PTUKI-100563635, Permit to Collect P-PTC-100563640). SAH thanks Queensland Museum staff involved with Project DIG along with the support provided for this Project from BHP and BMA. SAH thanks Caitlin Syme for collection management assistance.

## Data Availability

All strontium and carbon isotope data are available in the electronic supplementary material.

All R codes are available in the ‘Rockhampton Geoprovenance Package’ Zip file, and an OSF repository.

