## Supplementary Material (Text) for "Niche partitioning and limited mobility characterise Middle Pleistocene kangaroos from eastern Australia"

Supporting Information

**Taphonomy**

The four fossil deposits studied here have been described by Hocknull (2005a), Hocknull (2005b), Hocknull et al. (2007) and the dominant accumulating mechanisms are provided therein. All four deposits accumulated through a combination of pitfall and volant predators, specifically owl and ghost bat. Unlike open alluvial sites, these cavernous deposits are derived by fauna entering the cave entrances through animal behaviour (e.g., pitfall or predator) and not as alluvial sedimentary clasts transported from distance into a deposit.

Owls and ghost bats are clearly involved, with small-sized faunal dominating all accumulations, however, we cannot entirely rule out the influence of other predators known to have occurred in the environment at the time. Predator modification of bones, however, is inexplicably rare, with no clear indicators of large mammalian carnivore modification (i.e., modification by *Dasyurus*, *Sarcophilus*, *Thylacinus* or *Thylacoleo*). Volant predators known from the caves include owls and ghost bats, and although we cannot rule out large owl species, such as the Powerful Owl *Ninox strenua*, bringing in the smallest macropodid individuals (e.g. *Thylogale* and *Petrogale*), the foraging ranges of these predators would only produce a local signal with foraging ranges of owls and ghost bats from 100ms around the entrance to ~12 km^2^(Soderquist and Gibbons, 2007, Augusteyn et al., 2017).

Recently, the discovery of undescribed large raptorial remains from QML1384LU indicate the presence of a large volant predator, the size of a Wedge-tail eagle (*Aquila* spp.) or possibly an extinct giant form (Hocknull & Lawrence pers obs., 2023). Therefore, we cannot rule out these large raptorial predators as accumulator agents, although the rarity of their remains in the fossil deposits suggests they were a minor component of the local volant predator mix. The foraging ranges of *Aquila* species is highly variable, however, present day observations of Wedge-tail eagles suggest foraging ranges between 3 and ~30km^2^ (Cherriman, 2024). Regardless, to account for this possible yet rare bias, we subsampled a subset of individuals of *Thylogale*, *Petrogale* and *Notamacropus* to test whether geographic signal derived from Sr isotopes changed during mineralisation. A change from distal to local signal over time would justify a foraging range size that included both local and distal areas, and indicate the individuals dispersed to Mount Etna Caves from these distal areas and were not brought in volant predators.

### Strontium isotopes in vegetation

Overall, ^87^Sr/^86^Sr ratios in plants exhibit a broad range of values, from 0.704561 ± 0.000006 to 0.713402 ± 0.000006, with variations associated to underlying geology (Fig. S1 and S2). The lowest mean ^87^Sr/^86^Sr isotope ratios are observed in plants overlying basalts, 0.705716 ± 0.000024 (n = 2), and arenite-rudite, 0.705906 ± 0.000011. In contrast, higher mean ^87^Sr/^86^Sr isotope ratios are prevalent in plants growing over sand, 0.709178 ± 0.000005 (n = 2), and colluvium, 0.711016 ± 0.000011 (n = 2). When considering individual rock formations, the lowest ^87^Sr/^86^Sr isotope ratios are found in plants overlying Native Cate Andesite, 0.704561 ± 0.000006, and Dalma Basalt, 0.704704 ± 0.000012. The highest isotope ratios are found in the Back Creek Group Formation, 0.713402 ± 0.000006, and Qr-QLD, a Quaternary aged colluvium substrate, 0.711016 ± 0.000011 (n = 2).

^87^Sr/^86^Sr ratios show no significant variation within individual plants, with similar isotope ratios observed in the leaf (0.709178 ± 0.00005) and the fruiting body (0.709393 ± 0.000014) of a Tropical Almond tree (*Terminalia catappa*). This is in agreement with Kochergina et al. (2021) who reported no differences in ^87^Sr/^86^Sr isotope ratios in the xylem profile, pine needles, bark, and fine roots of European Spruce (*Picea abies*).

In most instances, measured ^87^Sr/^86^Sr isotope ratios do not differ between plants growing on the same rock formation (Supplementary Table S3). Consistent ^87^Sr/^86^Sr isotope ratios are measured in a native grass (0.708476 ± 0.000024), celerywood (*Polyscias elegans*) (0.708889 ± 0.000017), and *Ficus* sp. (0.708633 ± 0.000016), all growing on Mount Alma Formation limestone. Similarly, plants growing over the Rockhampton Group formation: a native grass (0.706367 ± 0.000016) and *Ficus* sp. (0.707998 ± 0.000007) showed similar isotope ratios, indicating ^87^Sr/^86^Sr ratios did not vary significantly across plant species from the same locality or substrate. However, two instances revealed significant differences in isotope ratios in plants from the same locality and substrate. In one case, a *Grass Tree* (*Xanthorrhoea* sp.) growing on a Trachyte plug (Mount Hedlow Trachyte) has a lower isotope ratio (0.707321 ± 0.000007) compared to isotope ratios measured in a deeper-rooted *Ficus* sp. (0.709084 ± 0.000006), and *Araucaria* sp. (0.709395 ± 0.000010) also underlain with Mount Hedlow Trachyte. In addition, a similar foreign influence was noted with a *Grass* specimen (0.713937 ± 0.000022) overlying the Mount Alma Formation limestone exhibiting a significantly higher ^87^Sr/^86^Sr isotope ratio than other plants from the same locality, including two different *Ficus* sp. (0.708476 ± 0.000024, 0.707912 ± 0.000039), a *Vine* sp. (0.708060 ± 0.000044), and a celerywood (0.708597 ± 0.000016). In both instances, the difference in isotope ratio between shallower rooted grasses and deeper-rooted vegetation could reflect the influence of an external source of strontium (potentially anthropogenic) on shallow rooted plants (Maurer et al., 2012). Given that both localities are managed by the Queensland Parks and Wildlife Service and have active weed-management programs, dissimilar ^87^Sr/^86^Sr in ratios in shallow-rooted plants compared to deeper-rooted plants at the same location, could be the result of anthropogenic inputs, such as herbicide application (Vitòria et al., 2004). Thus, these two grass specimens with anomalous isotope ratios were excluded from further analysis.

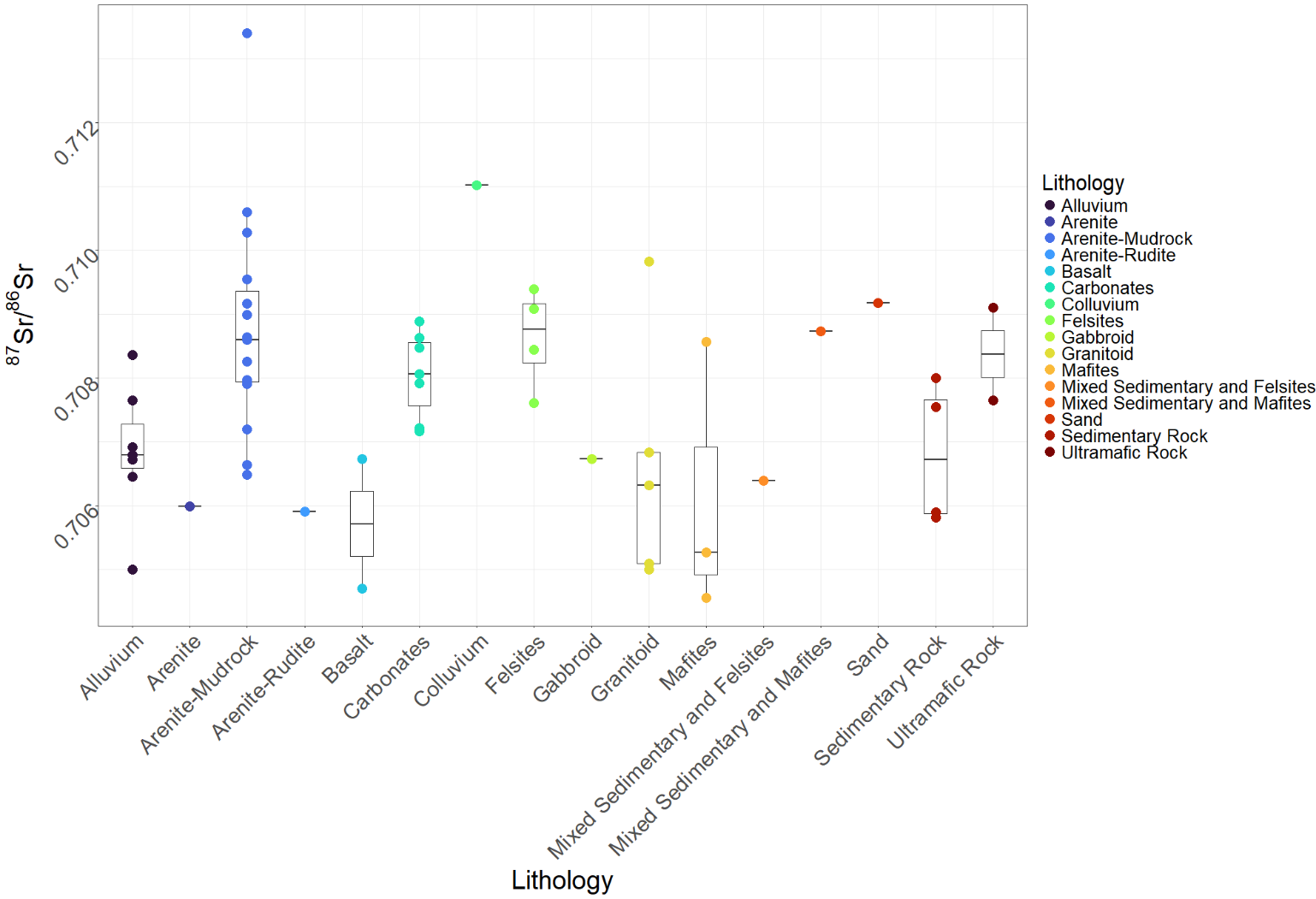

Figure S1. ^87^Sr/^86^Sr ratios in vegetation (n = 54) as a function of underlying geology. Lithology data are derived from the regional geological map: Queensland Government, [GeoResGlobe](https://georesglobe.information.qld.gov.au/) detailed 1:100K surface geology dataset © State of Queensland (Department of Resources) 2023. Figure created in RStudio (Team, 2019) using the ggplot2 package (Hadley, 2016).

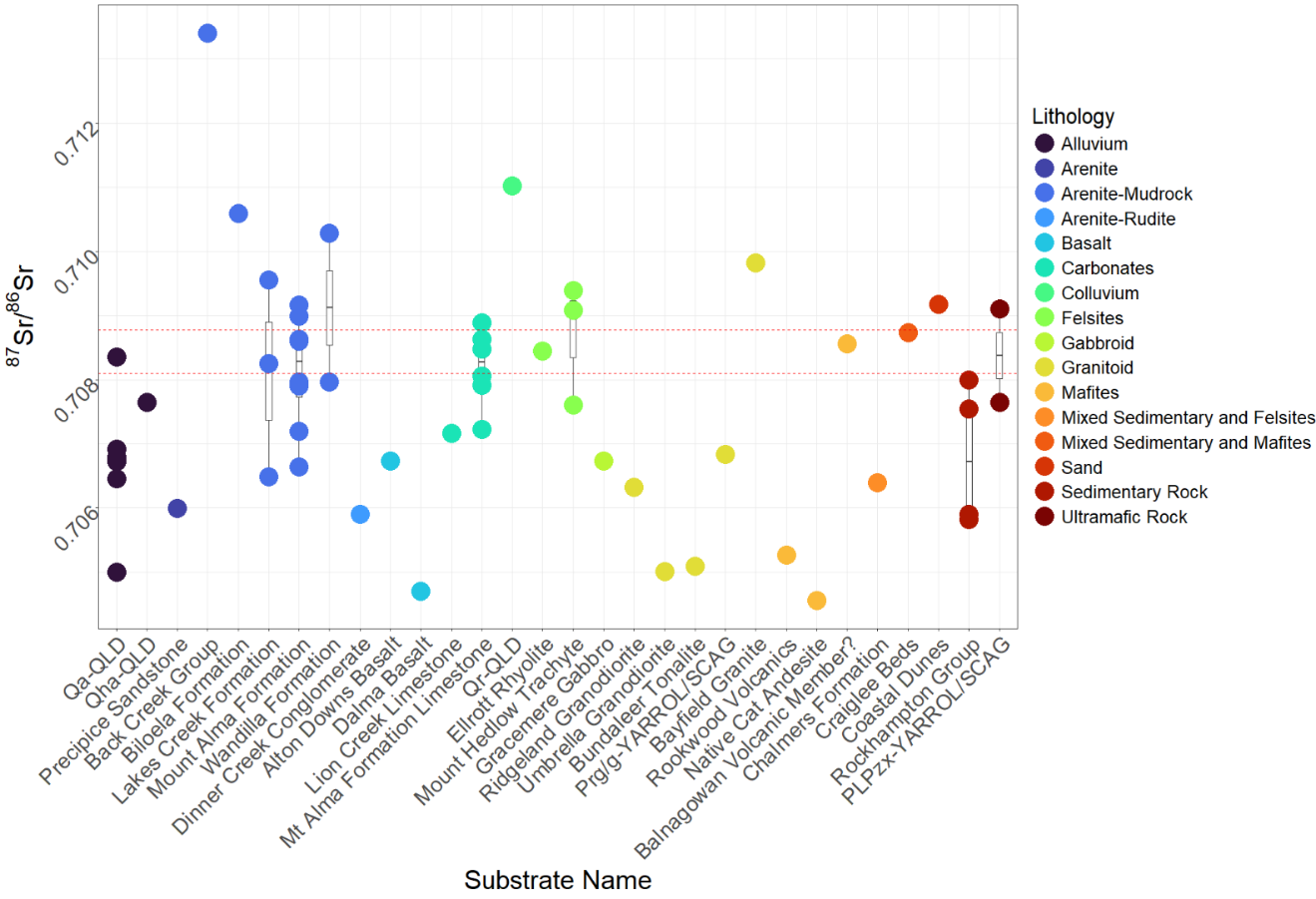

Figure S2. ^87^Sr/^86^Sr ratios measured in vegetation (n = 54) as a function of underlying geology. Lithology is derived from the regional geological map: Queensland Government, [GeoResGlobe](https://georesglobe.information.qld.gov.au/) detailed 1:100K surface geology dataset © State of Queensland (Department of Resources) 2023.

### Assessing the validity of the geographic assignment of fossil specimens

When using the *assignR* package (Ma et al., 2020), probability distribution is modulated by the spatial uncertainty (error associated with the isoscape) meaning there is potential trends in geographic assignment may be influenced by sampling bias. As such, we validated our geographic assignment model by running a second iteration keeping uncertainty constant across the isoscape. Comparing this constant uncertainty to our standard predictive model allows us to determine whether predictive ranges are a product of intensive local sampling biases, or that our predicted spatial distribution remains constant. When considering measured ^87^Sr/^86^Sr isotope ratios in vegetation, mean internal analytical uncertainty was 0.000012 and as such our constant uncertainty model utilised this value.

This comparative approach was undertaken for *Thylogale* as individual probability maps revealed distinct discrete foraging ranges across all individuals (See Fig. 3). Predicted foraging ranges for individuals under the constant uncertainty model somewhat differ when compared to individual probability maps generated using measured uncertainty (Fig. S3). Proposed foraging ranges appear more disjunct in the constant uncertainty model, restricted to a few isolated pixels. Despite the sporadic geographic assignment, high probabilities are observed to the north-west of fossil bearing caves are observed in 11 of the 12 specimens analysed, residing in a similar ‘local (north)’ area defined using our measured uncertainty model (Fig. S3). Contrastingly, despite localised foraging observed in 11/12 specimens, 5 individuals show higher probabilities to the east of Mount Etna Caves, albeit a handful of isolated pixels. Ultimately, while some variability can be observed between the constant uncertainty and measured uncertainty models, similar core ranges to the north-west suggest reported home ranges reflect geographic origin and are not a product of vegetation sampling bias.

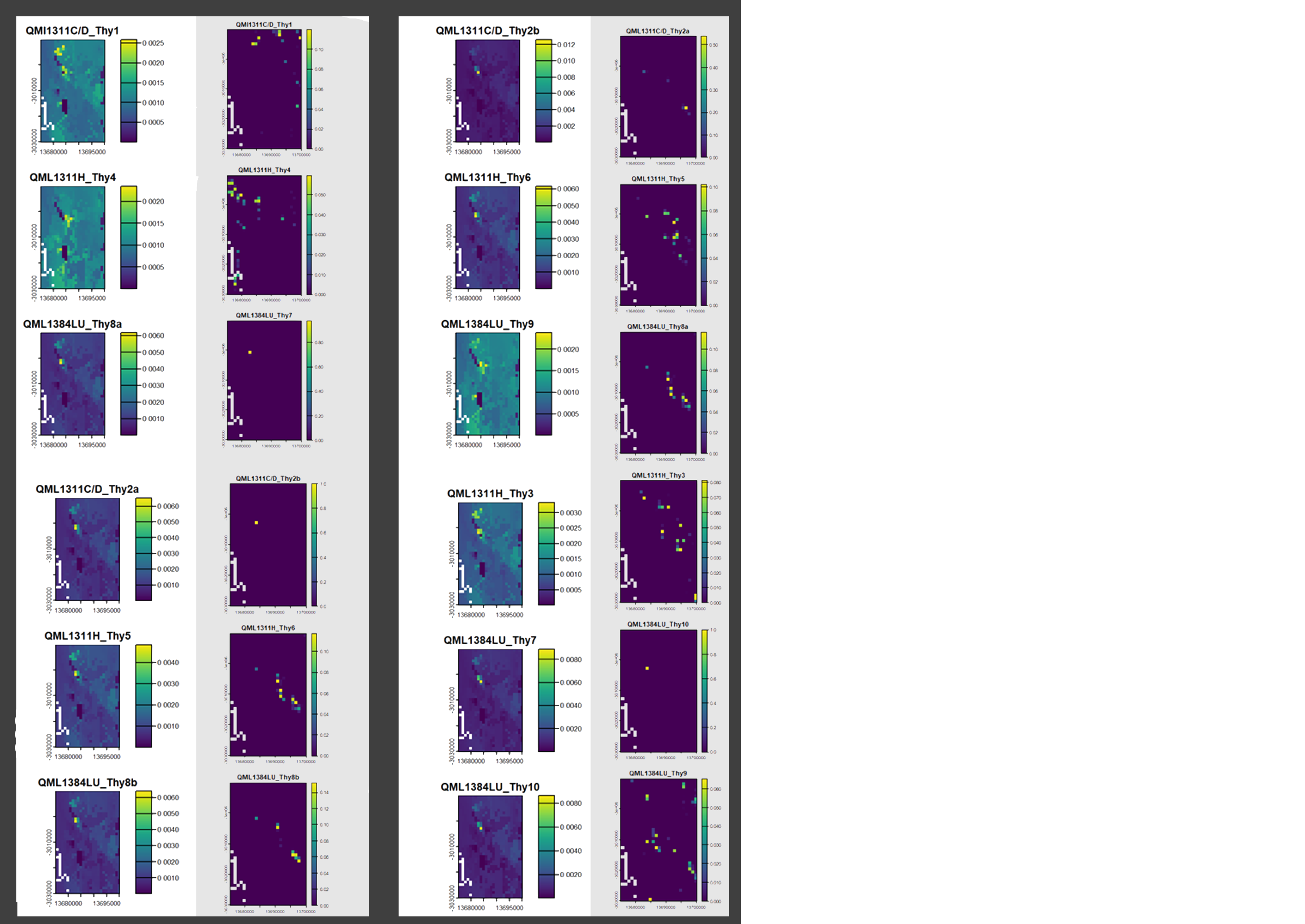

**Figure S3.** Individual probability maps for the *Thylogale* specimens, produced using measured uncertainty and constant uncertainty value of 0.000012 (Grey shading).

List of Figures

**Supplementary Figure 1.** Stratigraphy of Mount Etna Caves. Exposed fossil deposits at Mount Etna Limestone Mine, western benches. 1–5 QML1311; 1. A/B, 2. C/D, 3. F, 4. H, 5. J. 6–7 QML1384; 6. LU, 7. UU. 8. QML1310 Unit 2, 9. QML1383 A, 10. Open chamber to Speaking Tub Cave System, 11. QML1313, 12. Bench 0 (A/B), 13. QML1385. Reprinted with permission from Hocknull et al. (2007) © 2007 Elsevier B.V. All rights reserved.

**Supplementary Figure 2.** Images of macropodid dental samples included in this study (Table S1). Teeth have been grouped by taxa and can be identified by a unique Sample Name based on taxa and fossil locality.

**Supplementary Figure 3.** Images of a typical sampling regime (a) denoting where enamel powders were collected for a singular aliquot reflecting mean Sr composition and (b) more refined analyses with powder collected from the top and base of the crown.

**Supplementary Figure 4.** Random forest regression model for the bioavailable ^87^Sr/^86^ Sr dataset used in this study (a) Variable importance plot after selection of predictors by *VSURF*. (b) N-fold cross-validation results with best fit linear model (red line).

**Supplementary Figure 5.** Individual probability maps for *Thylogale* specimens, using the ‘regional’ Sr isoscape (see text for details). In this and following figures using the ‘regional’ Sr isoscape, x and y axes show longitude and latitude in an Eckert IV equal-area pseudocylindrical map projection, and north is oriented towards the top of the figure. field of view is 115 x 90 km.

**Supplementary Figure 6.** Individual probability maps for *Thylogale* specimens, using the 'narrow’ Sr isoscape (see text for details). In this and following figures using the ‘local’ Sr isoscape x and y axes show longitude and latitude in an Eckert IV equal-area pseudocylindrical map projection, and north is oriented towards the top of the figure. field of view is 25 x 40km.

**Supplementary Figure 7.** Joint probability maps for *Thylogale* specimens grouped by stratigraphy unit, using the ‘narrow’ Sr isoscape (see text for details). In this, and following age figures, the location of Mount Etna Caves is denoted with a solid red square.

**Supplementary Figure 8.** Individual probability maps for *Petrogale* specimens, using the ‘regional’ Sr isoscape (see text for details).

**Supplementary Figure 9.** Individual probability maps for *Petrogale* specimens, using the ‘narrow’ Sr isoscape (see text for details).

**Supplementary Figure 10.** Joint probability maps for *Petrogale* specimens grouped by fossil age, using both the ‘narrow’ and ‘regional’ Sr isoscapes (see text for details).

**Supplementary Figure 11.** Individual probability maps for *Notamacropus* specimens, using the ‘regional’ Sr isoscape (see text for details).

**Supplementary Figure 12.** Individual probability maps for *Notamacropus* specimens, using the ‘narrow’ Sr isoscape (see text for details).

**Supplementary Figure 13.** Joint probability maps for *Notamacropus* specimens grouped by fossil age, using the ‘narrow’ Sr isoscape (see text for details).

**Supplementary Figure 14.** Individual probability maps for *Protemnodon* specimens, using the ‘regional’ Sr isoscape (see text for details).

**Supplementary Figure 15.** Individual probability maps for *Protemnodon* specimens, using the ‘narrow’ Sr isoscape (see text for details).

**Supplementary Figure 16.** Joint probability maps for *Protemnodon* specimens grouped by fossil age, using the ‘narrow’ Sr isoscape (see text for details).

**Supplementary Figure 17.** (A) Reported occurrence of modern individuals belonging to *Petrogale herberti* (Orange) and *Petrogale inornata* (white) in the Mount Etna region. Points are compared to the ‘local’, ‘southern’ and ‘foreign’ joint probability maps defined in Figure 5.

**Supplementary Figure 18.** (a) Current extent of rainforest and vine-thicket vegetation types compared to historic foraging ranges defined in Figure 8. Vegetation extent reflects ‘Current major vegetation group (class)’ from the [Atlas of Living Australia Spatial portal](https://spatial.ala.org.au/).

**Supplementary Figure 19.** Joint probability maps for *Protemnodon* – Grouped by predicted size morphs defined in Table 4.

List of Tables

**Table S1.** Summary of measured ^87^Sr/^86^Sr isotope ratios in all marsupial enamel specimens examined during this study from Mount Etna, Queensland, Australia.

**Table S2.** Measured ^87^Sr/^86^Sr isotope ratios in plant samples collected from an 8,000 km^2^ region surround Mt Etna Caves (Denoted in Supplementary Figure 3).

**Table S3.** Mean drift corrected δ^13^C isotope values (± StdDev) measured in a series of IAEA Standards.

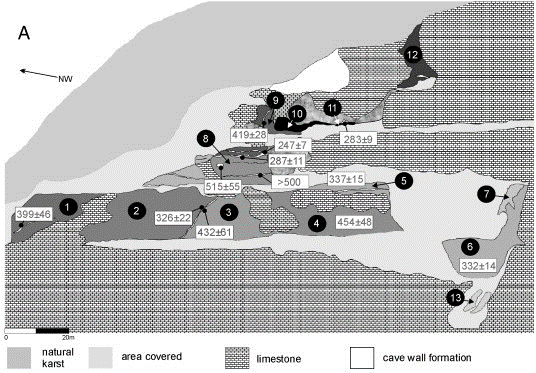

**Supplementary Figure 1.** Stratigraphy of Mount Etna Caves. Exposed fossil deposits at Mount Etna Limestone Mine, western benches. 1–5 QML1311; 1. A/B, 2. C/D, 3. F, 4. H, 5. J. 6–7 QML1384; 6. LU, 7. UU. 8. QML1310 Unit 2, 9. QML1383 A, 10. Open chamber to Speaking Tub Cave System, 11. QML1313, 12. Bench 0 (A/B), 13. QML1385. Reprinted with permission from Hocknull et al. (2007) © 2007 Elsevier B.V. All rights reserved.

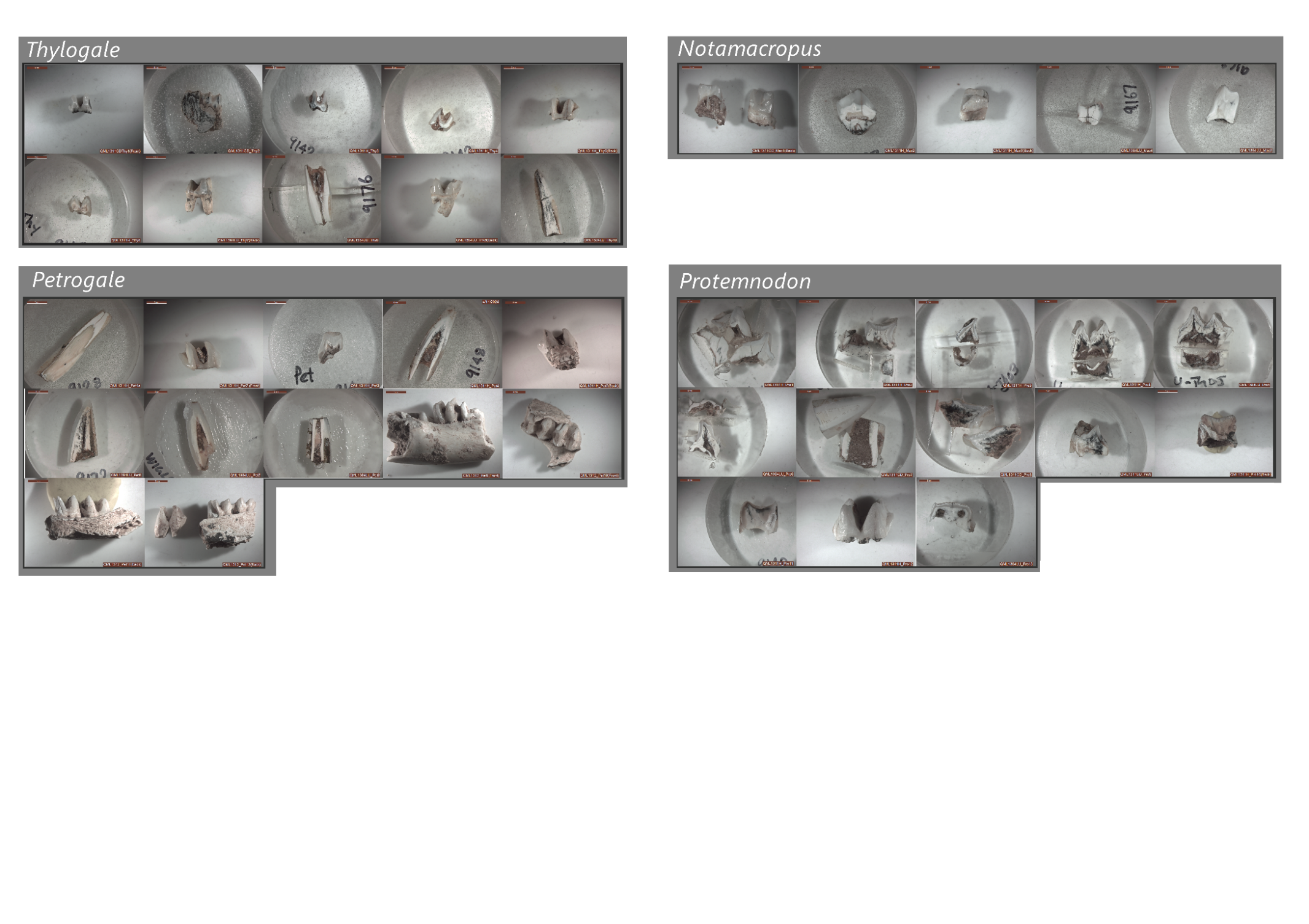

**Supplementary Figure 2.** Images of macropodid dental samples included in this study (Table S1). Teeth have been grouped by taxa and can be identified by a unique Sample Name based on taxa and fossil locality.

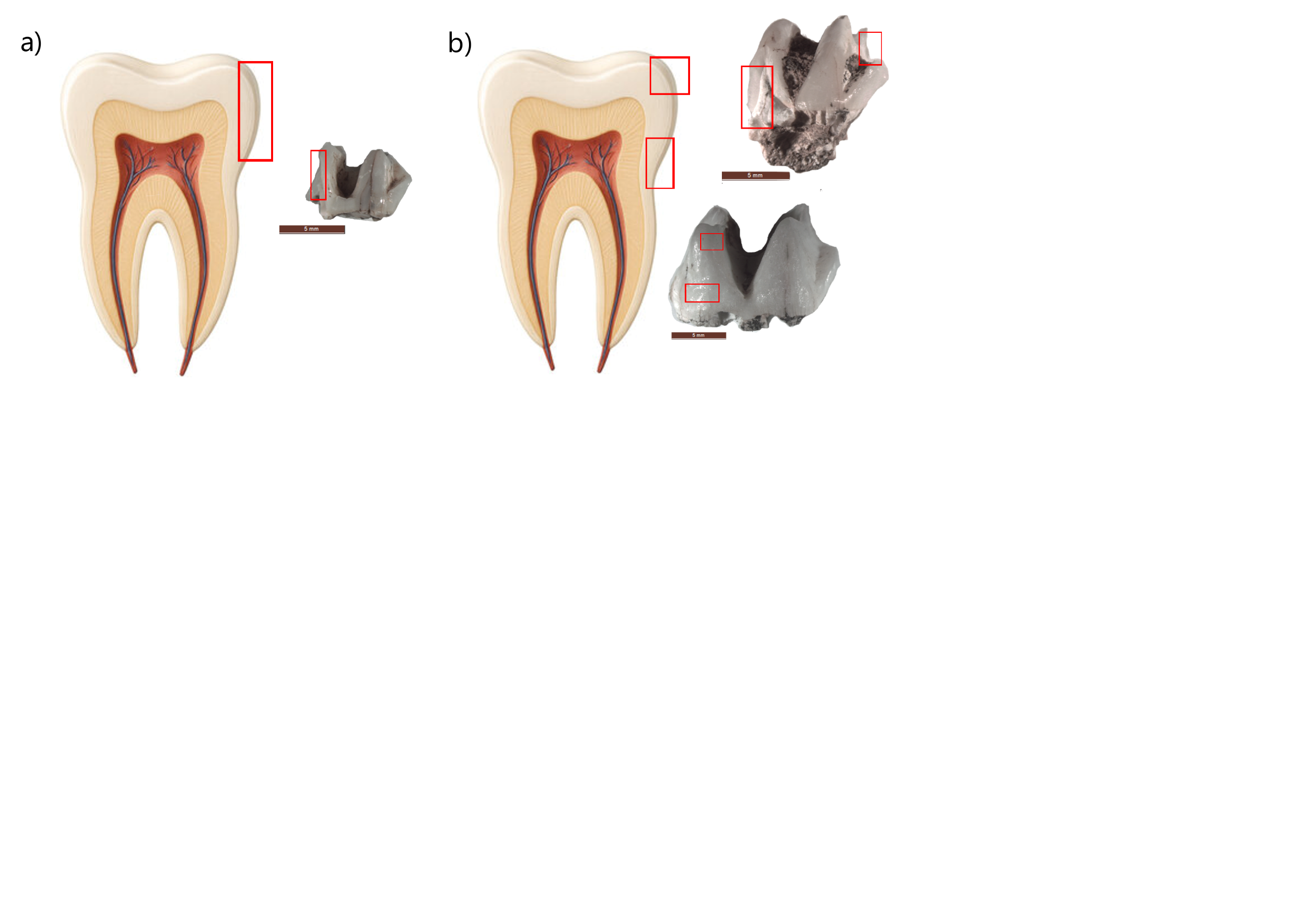

**Supplementary Figure 3.** Images of a typical sampling regime (a) denoting where enamel powders were collected for a singular aliquot reflecting mean Sr composition and (b) more refined analyses with powder collected from the top and base of the crown.

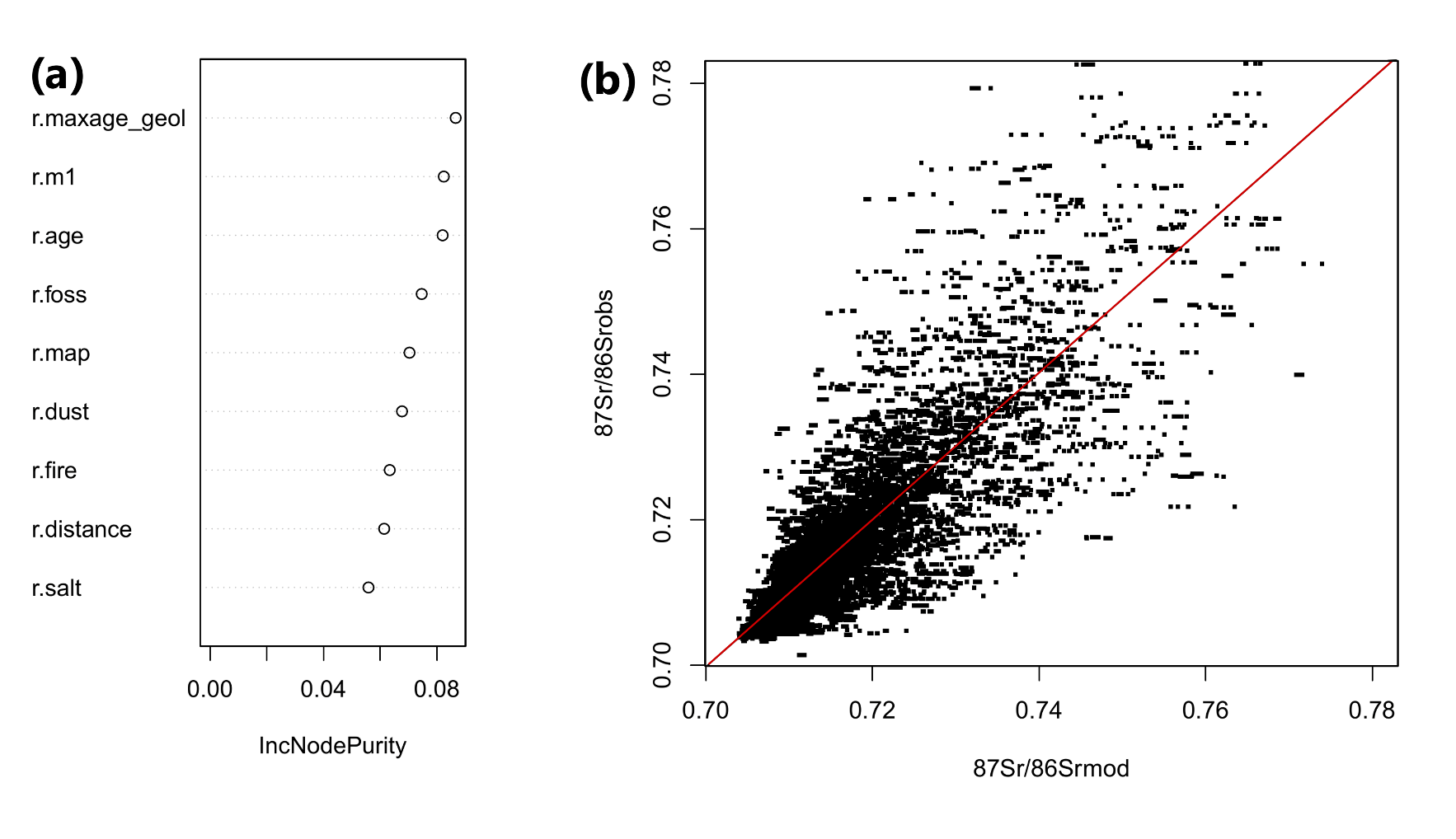

**Supplementary Figure 4.** Random forest regression model for the bioavailable ^87^Sr/^86^ Sr dataset used in this study (a) Variable importance plot after selection of predictors by *VSURF*. (b) N-fold cross-validation results with best fit linear model (red line).

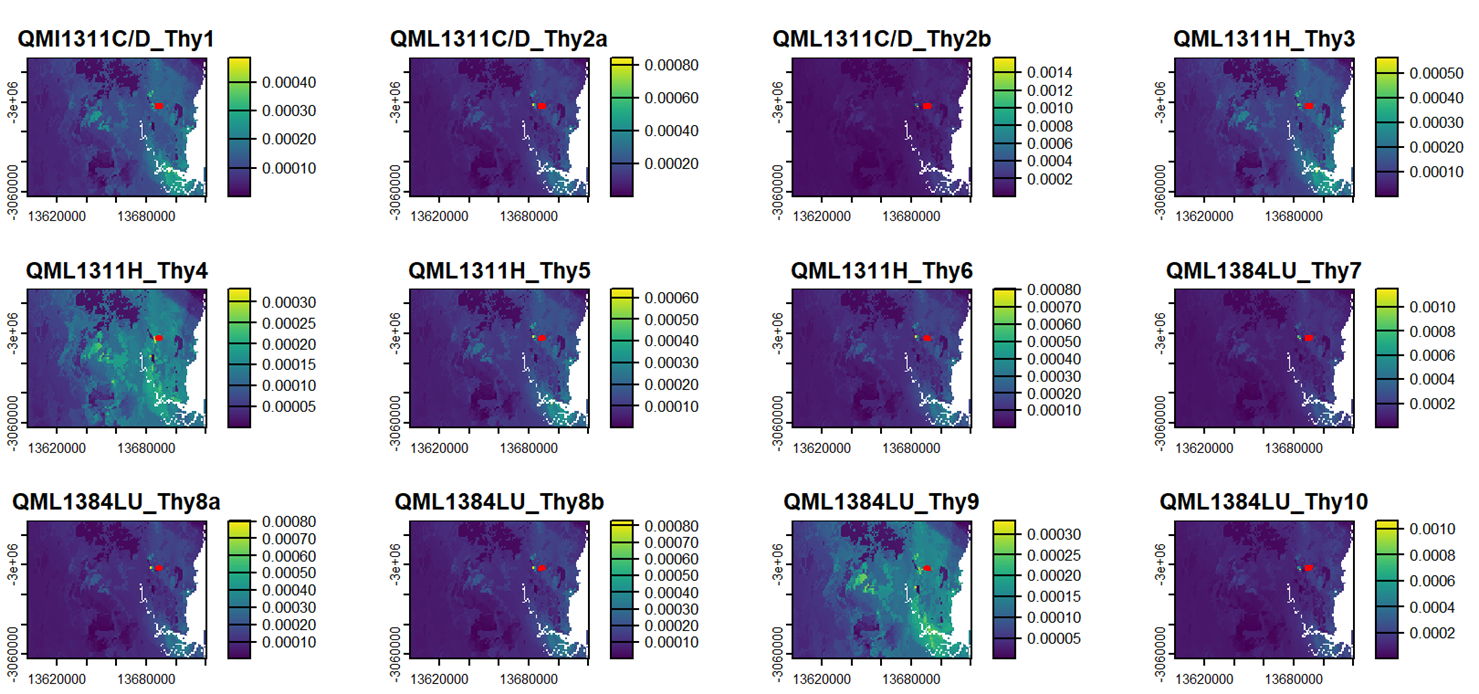

**Supplementary Figure 5.** Individual probability maps for *Thylogale* specimens, using the ‘regional’ Sr isoscape (see text for details). In this and following figures using the ‘regional’ Sr isoscape, x and y axes show longitude and latitude in an Eckert IV equal-area pseudocylindrical map projection, and north is oriented towards the top of the figure. field of view is 115 x 90 km.

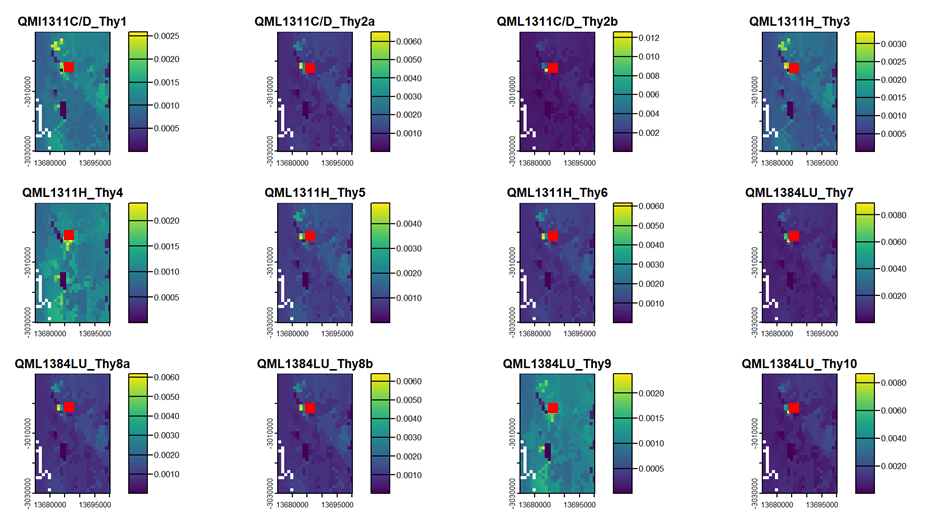

**Supplementary Figure 6.** Individual probability maps for *Thylogale* specimens, using the 'narrow’ Sr isoscape (see text for details). In this and following figures using the ‘local’ Sr isoscape x and y axes show longitude and latitude in an Eckert IV equal-area pseudocylindrical map projection, and north is oriented towards the top of the figure. field of view is 25 x 40km.

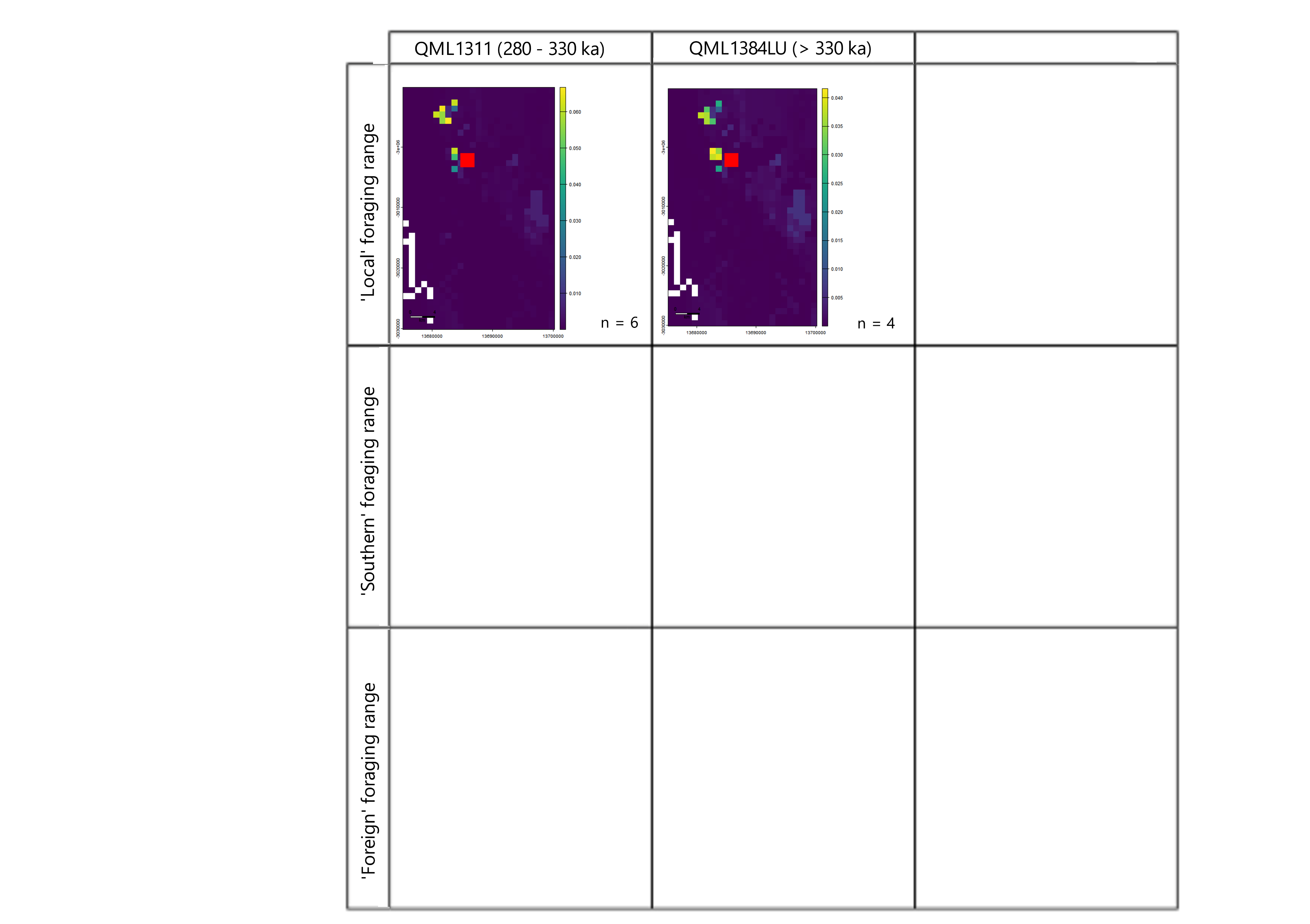
**Supplementary Figure 7.** ’Local’ foraging range over time for *Thylogale* showing limited change. Joint probability maps for *Thylogale* specimens grouped by stratigraphy unit, using the ‘narrow’ Sr isoscape (see text for details). Mount Etna Caves (red square).

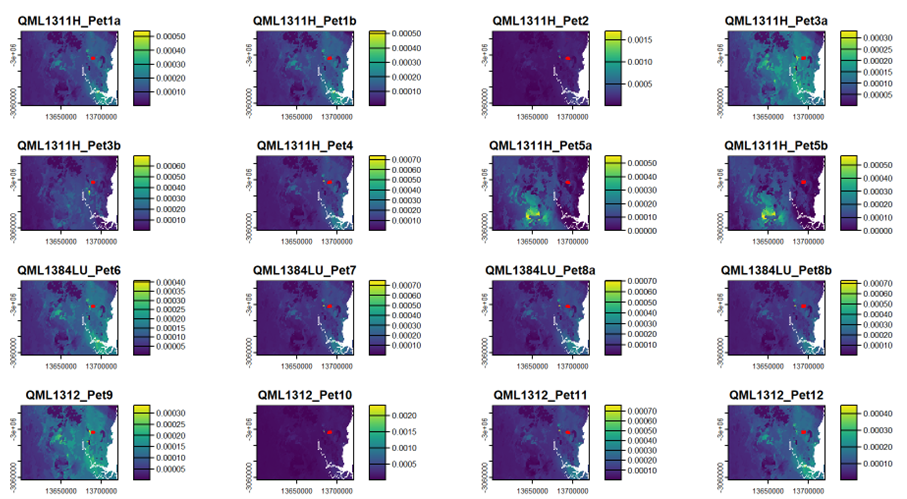

**Supplementary Figure 8.** Individual probability maps for *Petrogale* specimens, using the ‘regional’ Sr isoscape (see text for details).

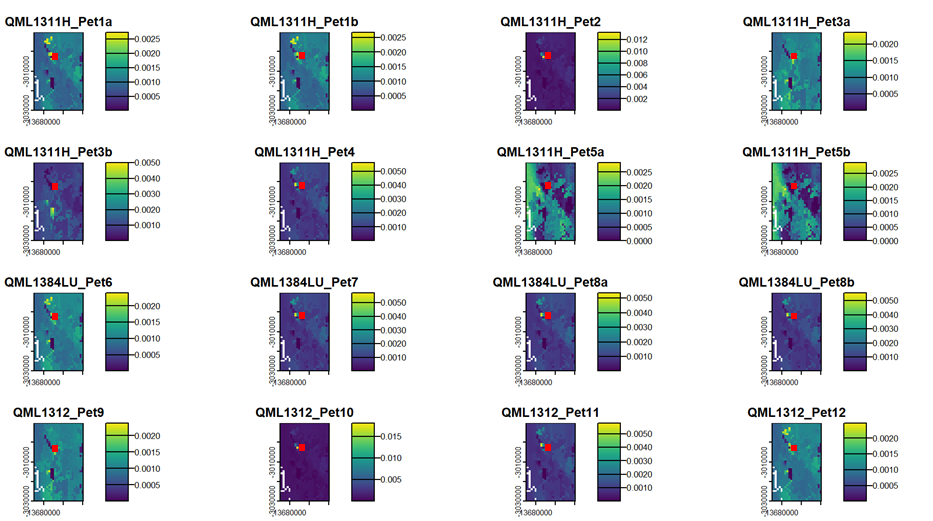

**Supplementary Figure 9.** Individual probability maps for *Petrogale* specimens, using the ‘narrow’ Sr isoscape (see text for details).

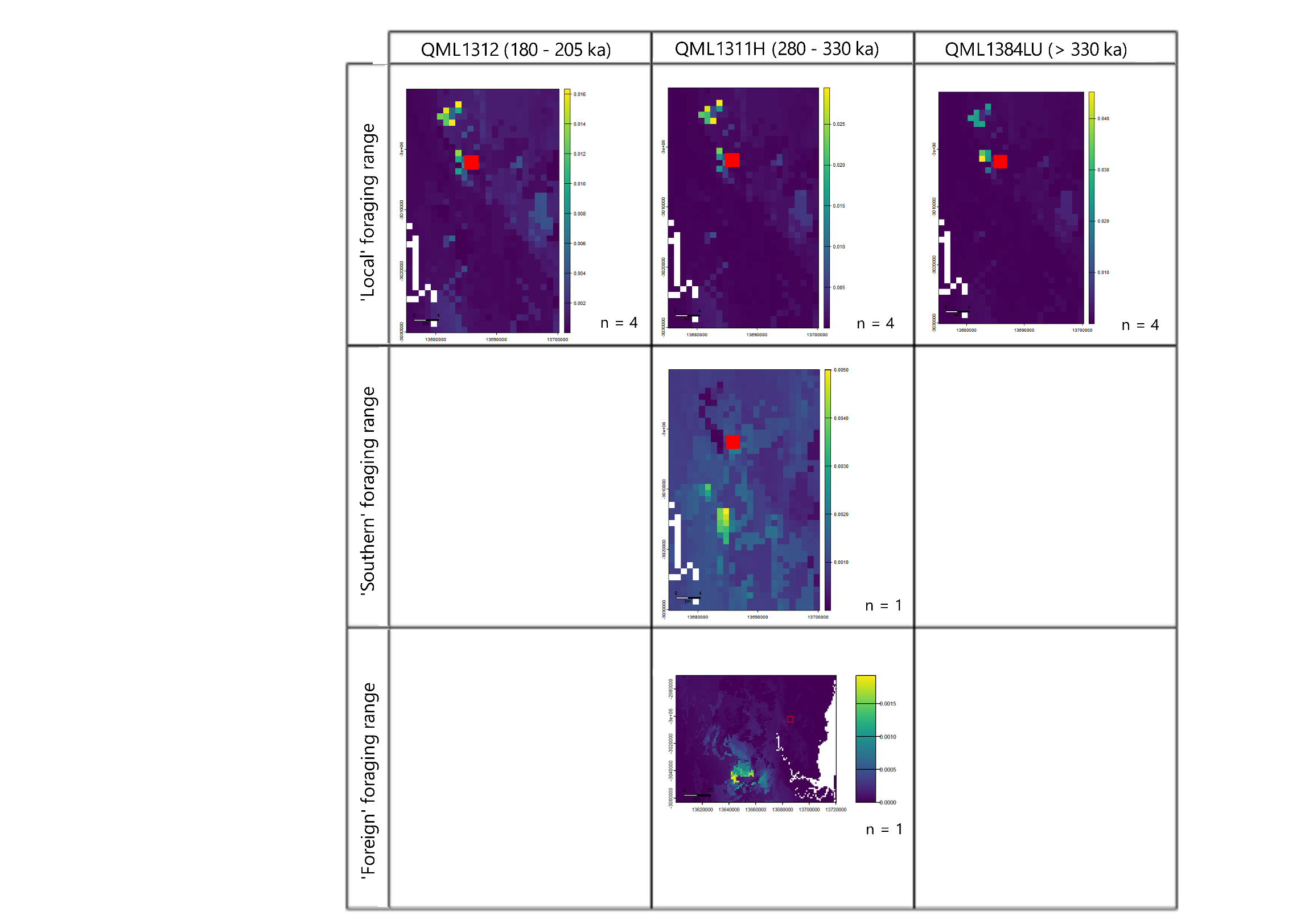

**Supplementary Figure 10.** Foraging range over time in *Petrogale* showing a possible expansion and then retraction in range over time. Joint probability maps for *Petrogale* specimens grouped by fossil age, using both the ‘narrow’ and ‘regional’ Sr isoscapes (see text for details). Mount Etna Caves (Red Square)

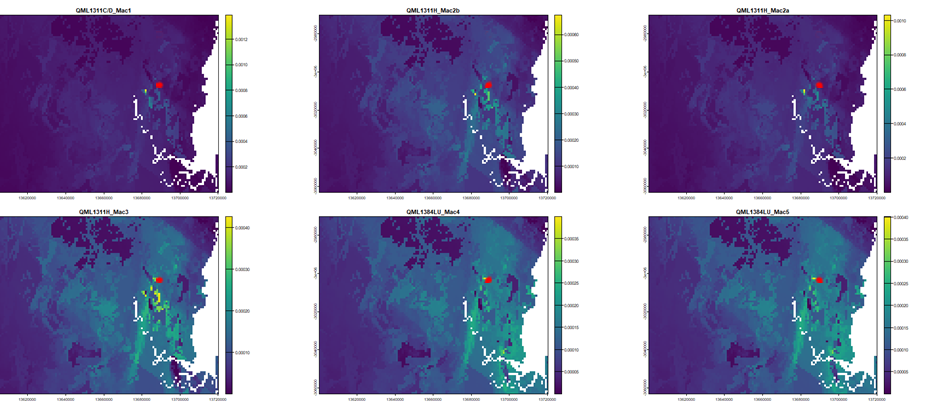

**Supplementary Figure 11.** Individual probability maps for *Notamacropus* specimens, using the ‘regional’ Sr isoscape (see text for details)

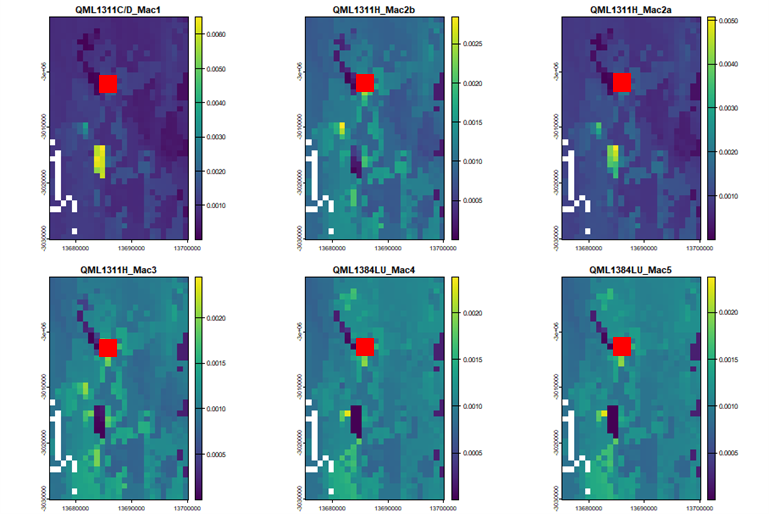

**Supplementary Figure 12.** Individual probability maps for *Notamacropus* specimens, using the ‘narrow’ Sr isoscape (see text for details).

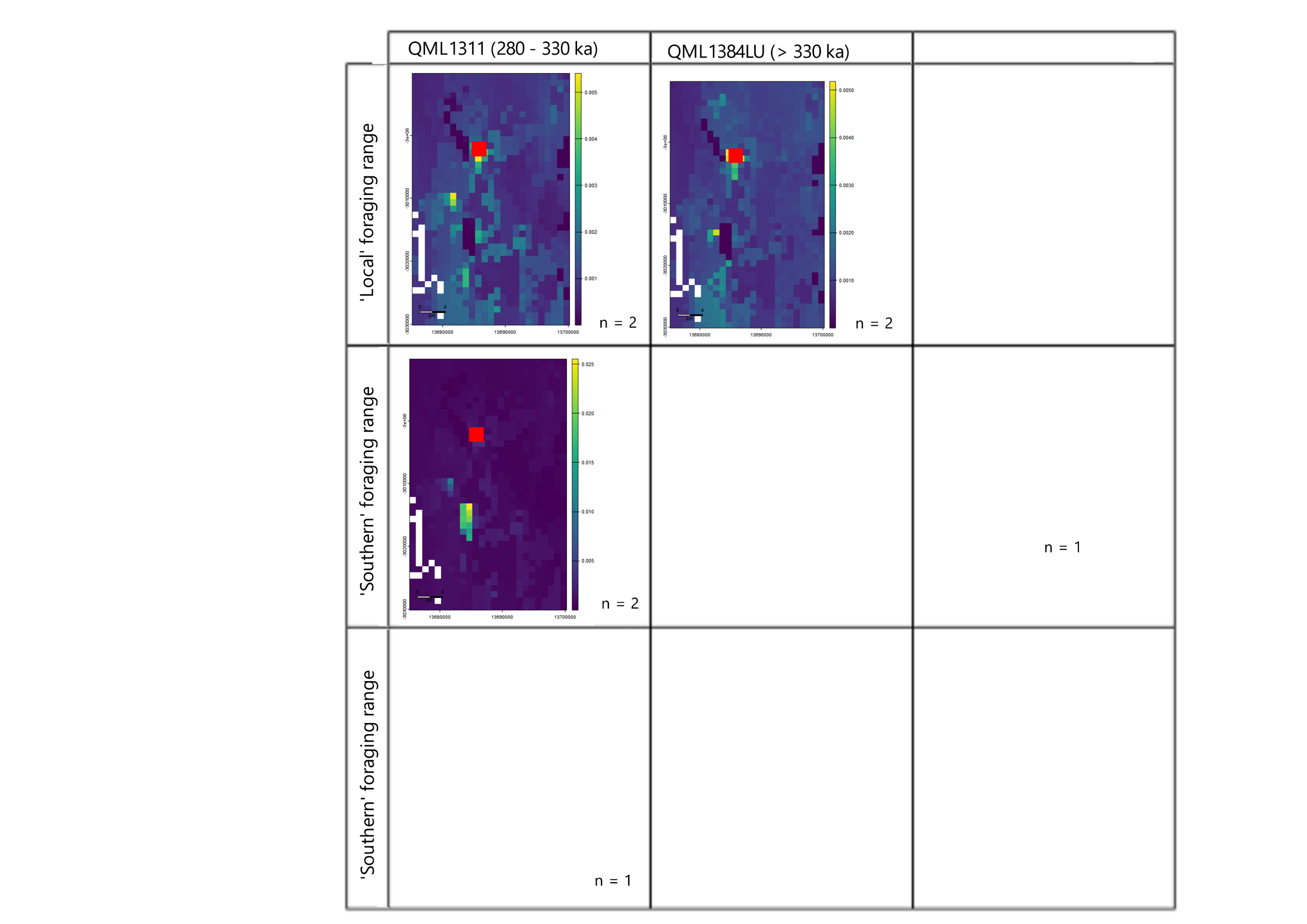

**Supplementary Figure 13.** Foraging range change of time in *Notamacropus* showing limited change over time. Joint probability maps for *Notamacropus* specimens grouped by fossil age, using the ‘narrow’ Sr isoscape (see text for details). Mount Etna Caves (red square).

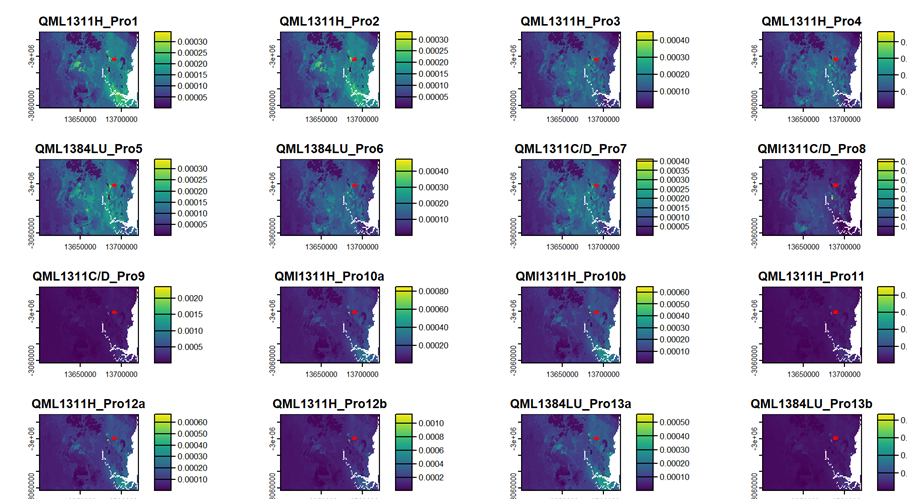

**Supplementary Figure 14.** Individual probability maps for *Protemnodon* specimens, using the ‘regional’ Sr isoscape (see text for details).

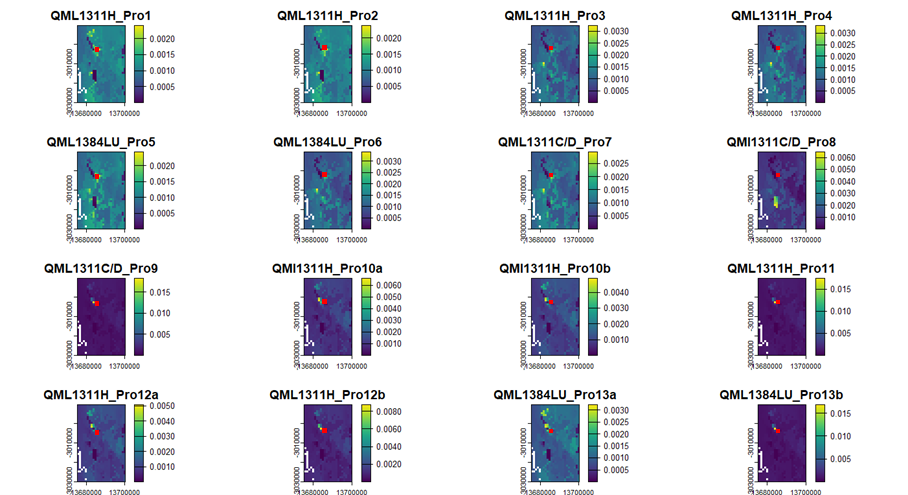

**Supplementary Figure 15.** Individual probability maps for *Protemnodon* specimens, using the ‘narrow’ Sr isoscape (see text for details).

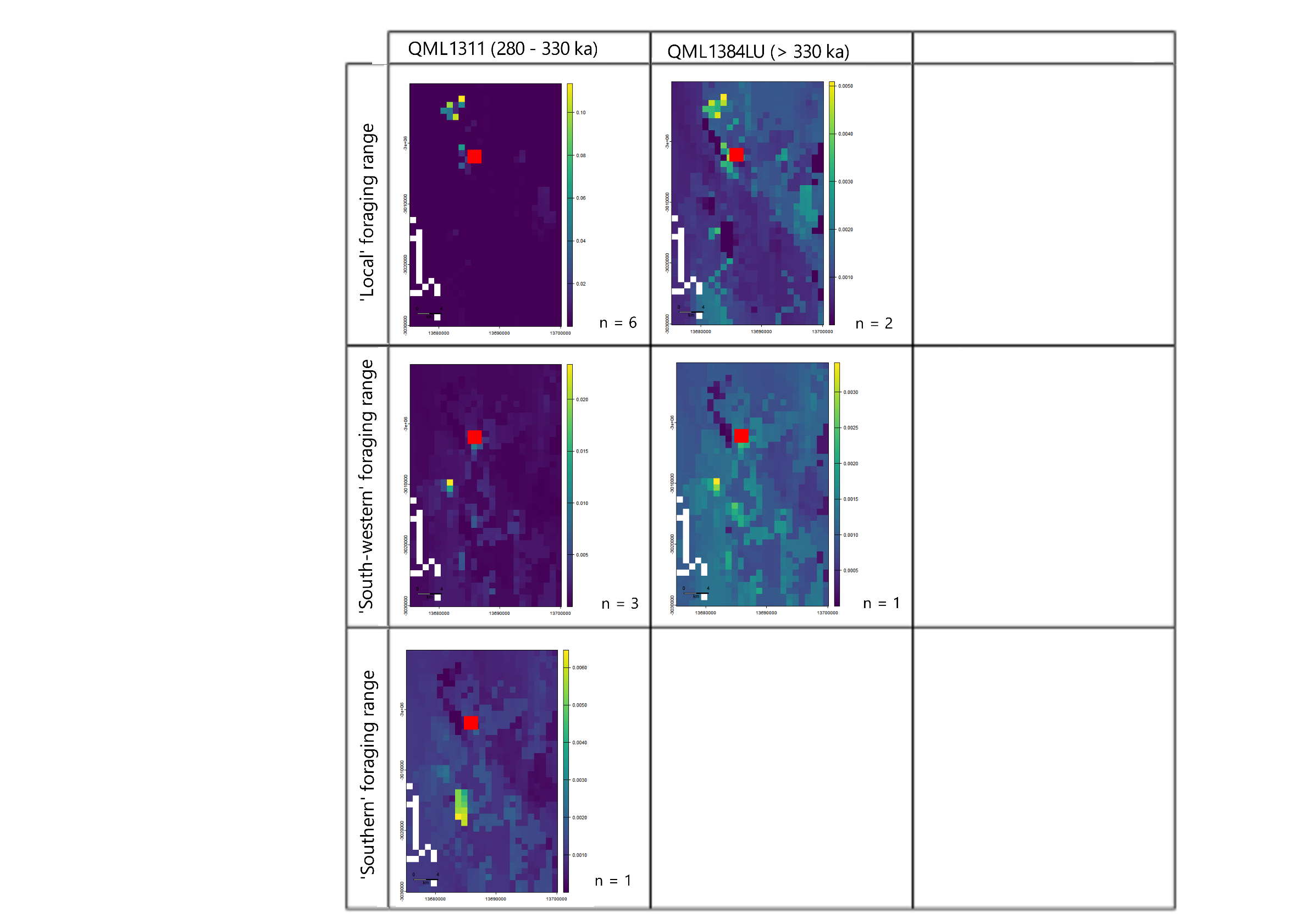

**Supplementary Figure 16.** Foraging range in *Protemnodon* showing limited change over time. Joint probability maps for *Protemnodon* specimens grouped by fossil age, using both the ‘narrow’ and ‘regional’ Sr isoscape (see text for details). Mount Etna Caves (red square).

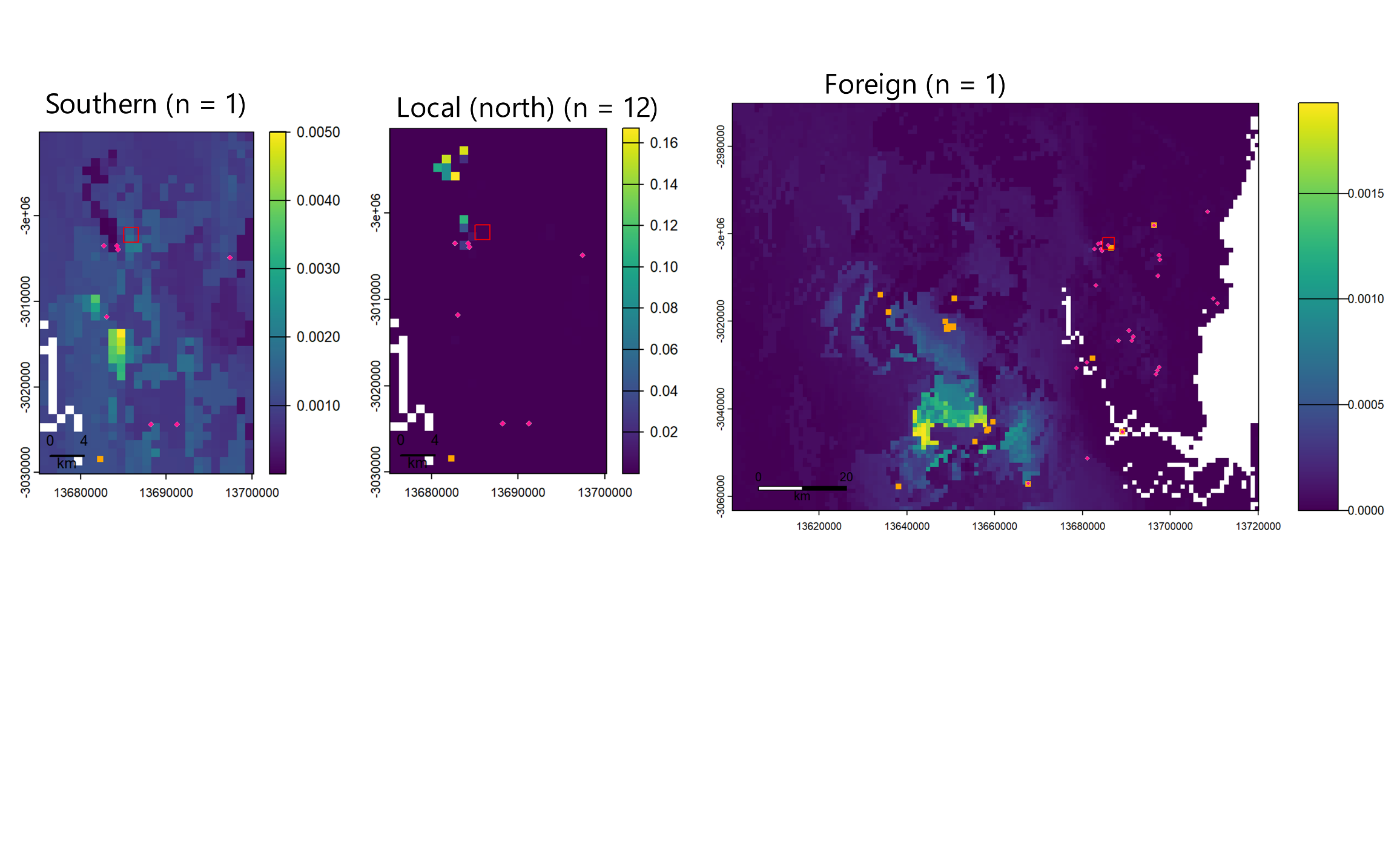

**Supplementary Figure 17.** Comparison of Pleistocene *Petrogale* with modern records. Reported occurrence of modern individuals belonging to *Petrogale herberti* (Orange) and *Petrogale inornata* (pink) in the Mount Etna region. Points are compared to the ‘southern’, ‘local (north)’, and ‘foreign’ joint probability maps defined in Figure 5. Modern reported Petrogale records from the [Atlas of Living Australia Spatial portal](https://spatial.ala.org.au/).

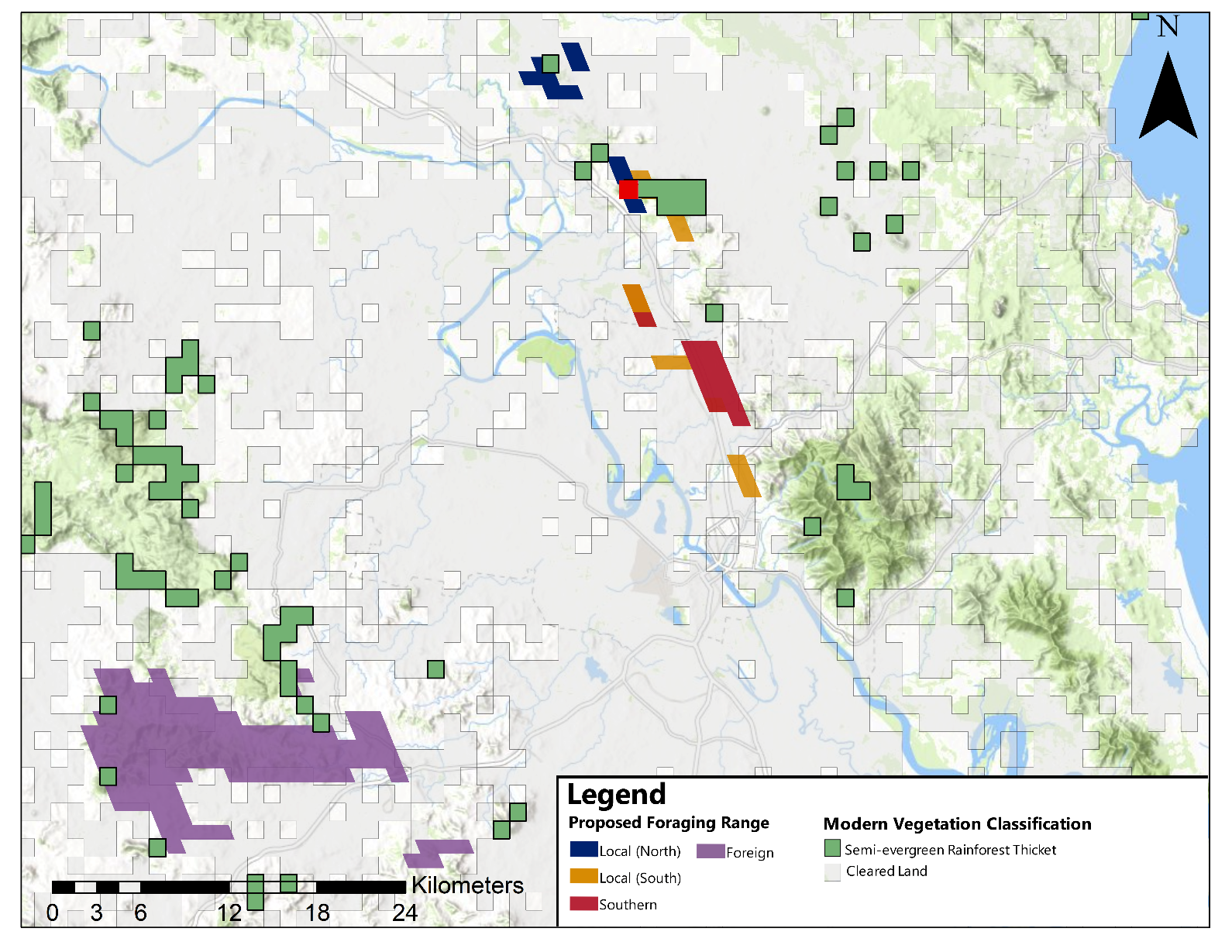

**Supplementary Figure 18.** (a) Current extent of rainforest and vine-thicket vegetation types compared to historic foraging ranges defined in Figure 8. Vegetation extent reflects ‘Current major vegetation group (class)’ from the [Atlas of Living Australia Spatial portal](https://spatial.ala.org.au/).

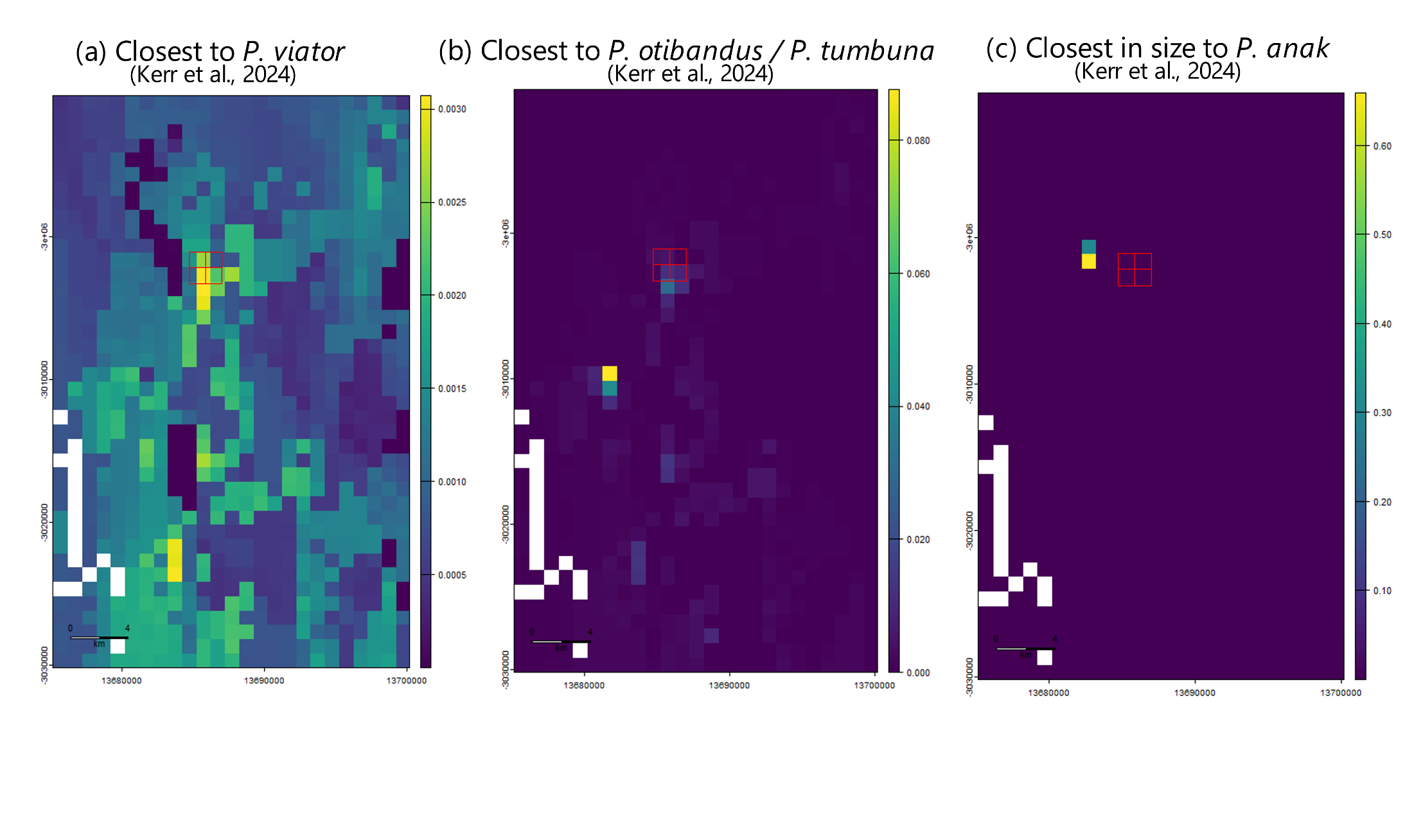

**Supplementary Figure 19.** Joint probability maps for *Protemnodon* – Grouped by predicted size morphs defined in Table 4. Mount Etna Caves (red square)

**Table S1.** Summary of measured ^87^Sr/^86^Sr isotope ratios in all marsupial enamel specimens examined during this study from Mount Etna, Queensland, Australia.

| Sample Name | Queensland Museum Locality (QML) | Deposit Age (ka) | Genus | Sample Type | Tooth | Sampling Location | ^87^Sr/^86^Sr | 2SE | Notes on size and closest taxon in (morphologically and size) |
| --- | --- | --- | --- | --- | --- | --- | --- | --- | --- |
| QML1311C/D_Mac1 | QML1311C/D | 280 - 330 | *Notamacropus* | Molar | M3 |  | 0.706974 | 0.000062 | ~20-25kg, *Notamacropus agilis* |
| QML1311H_Mac2a | QML1311H | 280 - 330 | *Notamacropus* | Molar | M3 | Base | 0.707418 | 0.000098 | ~20-25kg, *Notamacropus agilis* |
| QML1311H_Mac2b | QML1311H | 280 - 330 | *Notamacropus* | Molar | M3 | Top | 0.707089 | 0.000042 | ~20-25kg, *Notamacropus agilis* |
| QML1311H_Mac3 | QML1311H | 280 - 330 | *Notamacropus* | Molar | M2 |  | 0.707590 | 0.000030 | ~20-25kg, *Notamacropus agilis* |
| QML1384LU_Mac4 | QML1384LU | > 330 | *Notamacropus* | Molar | M2 |  | 0.707962 | 0.000036 | ~20-25kg, *Notamacropus agilis* |
| QML1384LU_Mac5 | QML1384LU | > 330 | *Notamacropus* |  |  |  | 0.707923 | 0.000079 | ~20-25kg, *Notamacropus agilis* |
| QML1311H_Pet1a | QML1311H | 280 - 330 | *Petrogale* | Incisor | I1 | Base | 0.708488 | 0.000055 | 6-11kg, *Petrogale* spp. |
| QML1311H_Pet1b | QML1311H | 280 - 330 | *Petrogale* | Incisor | I1 | Middle | 0.708442 | 0.000055 | 6-11kg, *Petrogale* spp. |
| QML1311H_Pet2 | QML1311H | 280 - 330 | *Petrogale* | Molar | M3 |  | 0.708924 | 0.000058 | 6-11kg, *Petrogale* (large, *penicillata* size) |
| QML1311H_Pet3a | QML1311H | 280 - 330 | *Petrogale* | Molar | M3 | Base | 0.707745 | 0.000087 | 6-11kg, *Petrogale* (large, *penicillata* size) |
| QML1311H_Pet3b | QML1311H | 280 - 330 | *Petrogale* | Molar | M3 | Top | 0.707093 | 0.000050 | 6-11kg, *Petrogale* (large, *herberti*-*penicillata* size) |
| QML1311H_Pet4 | QML1311H | 280 - 330 | *Petrogale* | Incisor | I1 |  | 0.708730 | 0.000065 | 6-11kg, *Petrogale* spp. |
| QML1311H_Pet5a | QML1311H | 280 - 330 | *Petrogale* | Molar | M4 | Base | 0.705145 | 0.000028 | 6-11kg, *Petrogale* (large, *penicillata* size). |
| QML1311H_Pet5b | QML1311H | 280 - 330 | *Petrogale* | Molar | M4 | Top | 0.705058 | 0.000026 | 6-11kg, *Petrogale* (large, *penicillata* size) |
| QML1384LU_Pet6 | QML1384LU | > 330 | *Petrogale* | Incisor | I1 |  | 0.708250 | 0.000088 | 6-11kg, *Petrogale* spp. |
| QML1384LU_Pet7 | QML1384LU | > 330 | *Petrogale* | Incisor | I1 |  | 0.708739 | 0.000047 | 6-11kg, *Petrogale* spp. |
| QML1384LU_Pet8a | QML1384LU | > 330 | *Petrogale* | Incisor | I1 |  | 0.708708 | 0.000075 | 6-11kg, *Petrogale* spp. |
| QML1384LU_Pet8b | QML1384LU | > 330 | *Petrogale* | Incisor | I1 |  | 0.708724 | 0.000071 | 6-11kg, *Petrogale* spp. |
| QML1312_Pet9 | QML1312 | 170 - 205 | *Petrogale* | Molar | M2-M4 |  | 0.707862 | 0.000049 | 6-11kg, *Petrogale* (small, closest to *inornata*) |
| QML1312_Pet10 | QML1312 | 170 - 205 | *Petrogale* | Molar | M2-3 |  | 0.708956 | 0.000054 | 6-11kg, *Petrogale* (small, closest to *inornata*) |
| QML1312_Pet11 | QML1312 | 170 - 205 | *Petrogale* | Molar | M3-4 |  | 0.708746 | 0.000049 | 6-11kg, *Petrogale* (small, closest to *inornata*) |
| QML1312_Pet12 | QML1312 | 170 - 205 | *Petrogale* | Molar | M1-M3 |  | 0.708314 | 0.000060 | 6-11kg, *Petrogale* (small, closest to *inornata*). |
| QML1311H_Pro1 | QML1311H | 280 - 330 | *Protemnodon* | Molar | M4 |  | 0.708160 | 0.000020 | Closest to *P. viator* (Kerr et al., 2024) |
| QML1311H_Pro2 | QML1311H | 280 - 330 | *Protemnodon* | Molar | m4 |  | 0.707992 | 0.000041 | Closest to *P. viator* (Kerr et al., 2024) |
| QML1311H_Pro3 | QML1311H | 280 - 330 | *Protemnodon* | Molar | M/4 |  | 0.707313 | 0.000045 | Closest to *P. otibandus / P. tumbuna* (Kerr et al., 2024) |
| QML1311H_Pro4 | QML1311H | 280 - 330 | *Protemnodon* | Molar | M2/ |  | 0.707288 | 0.000051 | Closest to *P. otibandus / P. tumbuna* (Kerr et al., 2024) |
| QML1384LU_Pro5 | QML1384LU | > 330 | *Protemnodon* | Molar | M4/ |  | 0.707597 | 0.000039 | Closest to *P.otibandus/P.tumbuna* (Kerr et al., 2024) |
| QML1384LU_Pro6 | QML1384LU | > 330 | *Protemnodon* | Molar | M2/ |  | 0.707258 | 0.000019 | Closest to *P. otibandus / P. tumbuna* (Kerr et al., 2024) |
| QML1311C/D_Pro7 | QML1311C/D | 280 - 330 | *Protemnodon* | Incisor | i1 |  | 0.707393 | 0.000023 | ?*P. otibandus* |
| QMl1311C/D_Pro8 | QML1311C/D | 280 - 330 | *Protemnodon* | Molar | m4 |  | 0.706570 | 0.000018 | Closest to P. viator (Kerr et al, 2024) |
| QML1311C/D_Pro9 | QML1311C/D | 280 - 330 | *Protemnodon* | Molar | M4/ |  | 0.708958 | 0.000095 | Closest to *P. anak/P.tumbuna/P.otibandus* (Kerr et al., 2024) |
| QMl1311H_Pro10a | QML1311H | 280 - 330 | *Protemnodon* | Molar | m2 | Base | 0.708858 | 0.000068 | Closest in*P. otibandus//P.anak* (Kerr et al., 2024) |
| QMl1311H_Pro10b | QML1311H | 280 - 330 | *Protemnodon* | Molar | m2 | Top | 0.708671 | 0.000054 | Closest to *P. otibandus/P.anak* (Kerr et al., 2024) |
| QML1311H_Pro11 | QML1311H | 280 - 330 | *Protemnodon* | Molar | m2 or m3 |  | 0.708952 | 0.000046 | Closest to *P. anak/P.tumbuna/P.otibandus* (Kerr et al., 2024) |
| QML1311H_Pro12a | QML1311H | 280 - 330 | *Protemnodon* | Molar | M4 | Base | 0.708686 | 0.000047 | Closest to. *P.anak/P. mamkurra* (Kerr et al, 2024) |
| QML1311H_Pro12b | QML1311H | 280 - 330 | *Protemnodon* | Molar | M4 | Top | 0.708894 | 0.000060 | Closest to. *P.anak/P. mamkurra* (Kerr et al, 2024) |
| QML1384LU_Pro13a | QML1384LU | > 330 | *Protemnodon* | Molar | M3/ | Base | 0.708550 | 0.000041 | Closest to *P. anak/P.tumbuna/P.otibandus* (Kerr et al., 2024) |
| QML1384LU_Pro13b | QML1384LU | > 330 | *Protemnodon* | Molar | M3/ | Top | 0.708948 | 0.000086 | Closest to *P. anak/P.tumbuna/P.otibandus* (Kerr et al., 2024) |
| QMl1311C/D_Thy1 | QML1311C/D | 280 - 330 | *Thylogale* | Molar | m/3 | Base | 0.708378 | 0.000102 | <7kg, *Thylogale, (stigmatica/billardierii)* |
| QML1311C/D_Thy2a | QML1311C/D | 280 - 330 | *Thylogale* | Molar | m/2 | m/2 | 0.708838 | 0.000077 | <7kg, *Thylogale (stigmatica/billardierii)* |
| QML1311C/D_Thy2b | QML1311C/D | 280 - 330 | *Thylogale* | Molar | m/3 | m/3 | 0.709112 | 0.000072 | <7kg, *Thylogale (stigmatica/billardierii* |
| QML1311H_Thy3 | QML1311H | 280 - 330 | *Thylogale* | Molar | m/3 |  | 0.708558 | 0.000067 | <7kg, *Thylogale(stigmatica/billardierii)* |
| QML1311H_Thy4 | QML1311H | 280 - 330 | *Thylogale* | Molar | m/3 |  | 0.707741 | 0.000033 | <7kg, *Thylogale (stigmatica/billardierii)* |
| QML1311H_Thy5 | QML1311H | 280 - 330 | *Thylogale* | Molar | M4/ |  | 0.708667 | 0.000122 | <7kg, *Thylogale thetis* (large individual) |
| QML1311H_Thy6 | QML1311H | 280 - 330 | *Thylogale* | Molar | M2/ |  | 0.708785 | 0.000031 | <7kg, *Thylogale (stigmatica/billardierii)* |
| QML1384LU_Thy7 | QML1384LU | > 330 | *Thylogale* | Molar | M2/ |  | 0.708895 | 0.000077 | <7kg, *Thylogale thetis* (large individual) |
| QML1384LU_Thy8a | QML1384LU | > 330 | *Thylogale* | Incisor | i/1 | Base | 0.708786 | 0.000034 | <7kg, *Thylogale (stigmatica/billardierii)* |
| QML1384LU_Thy8b | QML1384LU | > 330 | *Thylogale* | Incisor | i/1 | Top | 0.708817 | 0.000057 | <7kg, *Thylogale (stigmatica/billardierii)* |
| QML1384LU_Thy9 | QML1384LU | > 330 | *Thylogale* | Molar | M4/ |  | 0.708049 | 0.000088 | <7kg, *Thylogale thetis* (large individual) |
| QML1384LU_Thy10 | QML1384LU | > 330 | *Thylogale* | Incisor | i/1 |  | 0.709139 | 0.000128 | <7kg, *Thylogale (stigmatica/billardierii)* |

Age ranges for QML1311H, QML1311C/D and QML1384LU are based on those proposed in Laurikainen Gaete et al. (2025)

Age range for QML1312 based on ages from Hocknull et al. (2007)

**Table S2.** Measured ^87^Sr/^86^Sr isotope ratios in plant samples collected from an 8,000 km^2^ region surround Mt Etna Caves (Denoted in Supplementary Figure 3).

| Sample name | Dominant Rock | Substrate Name | Location | Longitude | Latitude | Vegetation Type | 87Sr/86Sr | 2SE |
| --- | --- | --- | --- | --- | --- | --- | --- | --- |
| MtE_V19 | Basalt | Alton Downs Basalt | Etna Creek Rd | 150.46341 | -23.21681 | *Casurina* sp | 0.706728 | 0.000022 |
| MtE_V12 | Arenite-Mudrock | Back Creek Group | Riverslea Rd | 149.96127 | -23.5931 | *Eucalyptus* sp. | 0.713402 | 0.000006 |
| MtE_V39 | Mafites | Balnagowan Volcanic Member | Limestone Creek | 150.68951 | -23.16904 | *Eucalyptus* sp. | 0.708562 | 0.000006 |
| MtE_V31 | Granitoid | Bayfield Granite | Byfield Rd | 150.68599 | -22.947 | *Eucalyptus* sp. | 0.709826 | 0.000007 |
| MtE_V11 | Arenite-Mudrock | Biloela Formation | Riverslea Rd | 149.97316 | -23.60166 | *Eucalyptus* sp. | 0.710593 | 0.000006 |
| MtE_V3 | Granitoid | Bundaleer Tonalite | Kabra Rd | 150.42341 | -23.50868 | *Eucalyptus* sp. | 0.705094 | 0.000005 |
| MtE_V47 | Mixed Sedimentary and Felsites | Chalmers Formation | Thompson Point Rd | 150.67901 | -23.42442 | *Eucalyptus* sp. | 0.706395 | 0.000014 |
| MtE_V33B | Sand | Coastal Dunes | Kemp Beach | 150.79158 | -23.17405 | *Terminalia catappa* (Leaf) | 0.709393 | 0.000014 |
| MtE_V33 | Sand | Coastal Dunes | Kemp Beach | 150.79158 | -23.17405 | *Terminalia catappa* (Fruit) | 0.709178 | 0.000005 |
| MtE_V16 | Mixed Sedimentary and Mafites | Craiglee Beds | Glenroy Rd | 150.11215 | -23.18598 | *Eucalyptus* sp. | 0.708733 | 0.000006 |
| MtE_V5 | Basalt | Dalma Basalt | Stanwell | 150.31561 | -23.48901 | Pea Sp. Unknown | 0.704704 | 0.000012 |
| MtE_V7 | Arenite-Rudite | Dinner Creek Conglomerate | Rosewood Wycarbah Rd | 150.13593 | -23.50298 | *Eucalyptus* sp. | 0.705906 | 0.000011 |
| MtE_V35 |  | Doonside Formation | Camms Rd | 150.79158 | -23.17405 | Paperbark sp | 0.706860 | 0.000011 |
| MtE_V49 | Felsites | Ellrott Rhyolite | Broadmount Rd | 150.74017 | 23.48489 | *Eucalyptus* sp. | 0.708443 | 0.000007 |
| MtE_VB11 | Gabbroid | Gracemere Gabbro | Watts Rd | 150.4716 | -23.4476 | *Eucalyptus* sp. | 0.706729 | 0.000007 |
| MtE_V29 | Arenite-Mudrock | Lakes Creek Formation | Rossmoya Rd | 150.45792 | -23.14538 | Unknown | 0.706486 | 0.000007 |
| MtE_V37 | Arenite-Mudrock | Lakes Creek Formation | Artillery Rd | 150.58864 | -23.23589 | *Eucalyptus* sp. | 0.709549 | 0.000006 |
| MtE_VB14 |  | Limestone | Capricorn Caves Walking Track | 150.4920768 | -23.1677008 | *Asplenium* sp. | 0.707336 | 0.000033 |
| MtE_V13 | Carbonates | Lion Creek Limestone | Broughton Rd | 150.25421 | -23.38099 | Unknown | 0.707161 | 0.000007 |
| MtE_V23 | Arenite-Mudrock | Mount Alma Formation | B Thomassons Rd | 150.4912467 | -23.1581361 | *Eucalyptus* sp. | 0.708254 | 0.000007 |
| MtE_V14 | Arenite-Mudrock | Mount Alma Formation | Glenroy Rd | 150.17264 | -23.26269 | *Eucalyptus* sp. | 0.707902 | 0.000005 |
| MtE_V21 | Arenite-Mudrock | Mount Alma Formation | Olsens Caves Rd | 150.4851574 | -23.1662095 | *Eucalyptus* sp. | 0.707193 | 0.000010 |
| MtE_V30 | Arenite-Mudrock | Mount Alma Formation | Bruce Hwy | 150.40319 | -23.09834 | Unknown | 0.708993 | 0.000011 |
| MtE_VB13 | Arenite-Mudrock | Mount Alma Formation | Capricorn Caves Walking Track | 150.4920768 | -23.1677008 | *Ficus* sp. | 0.706638 | 0.000023 |
| MtE_VB1 | Arenite-Mudrock | Mount Alma Formation | Camoo Caves | 150.4661 | -23.16373 | Unknown | 0.707965 | 0.000019 |
| MtE_VB5 | Arenite-Mudrock | Mount Alma Formation | Western ridge | 150.4493647 | -23.1576509 | *Ficus* sp. | 0.708597 | 0.000016 |
| MtE_VB6 | Arenite-Mudrock | Mount Alma Formation | Western Ridge | 150.4494404 | -23.157547 | *Polyscias elegans* | 0.708634 | 0.000011 |
| MtE_VB9 | Arenite-Mudrock | Mount Alma Formation |  | 150.45385 | -23.15789 | Bauhemia | 0.709169 | 0.000008 |
| MtE_VB12 | Carbonates | Mount Alma Formation Limestone | Capricorn Caves Walking Track | 150.492327 | -23.165769 | *Eucalyptus* sp. | 0.707221 | 0.000014 |
| MtE_VB17 | Carbonates | Mount Alma Formation Limestone | Capricorn Caves | 150.4897544 | -23.1640558 | *Ficus* sp. | 0.707912 | 0.000039 |
| MtE_VB18 | Carbonates | Mount Alma Formation Limestone | Capricorn Caves | 150.4897544 | -23.1640558 | Vine Sp | 0.708060 | 0.000044 |
| MtE_VB2B* | Carbonates | Mount Alma Formation Limestone | Camoo Caves | 150.46538 | -23.16334 | Grass | 0.713937 | 0.000022 |
| MtE_VB2A | Carbonates | Mount Alma Formation Limestone | Camoo Caves | 150.46538 | -23.16334 | *Ficus* sp. | 0.708476 | 0.000024 |
| MtE_VB3 | Carbonates | Mount Alma Formation Limestone | Etna Ridge | 150.4503105 | -23.1563435 | White Cedar | 0.708889 | 0.000017 |
| MtE_VB4 | Carbonates | Mount Alma Formation Limestone | Western Ridge | 150.4502578 | -23.1564548 | *Ficus* sp. | 0.708633 | 0.000016 |
| MtE_V34A | Felsites | Mount Hedlow Trachyte | Capricorn Coast NP | 150.79373 | -23.16374 | *Ficus* sp. | 0.709084 | 0.000006 |
| MtE_V34B* | Felsites | Mount Hedlow Trachyte | Capricorn Coast NP | 150.79341 | -23.16293 | *Xanthorrhoea* sp | 0.707321 | 0.000007 |
| MtE_V38 | Felsites | Mount Hedlow Trachyte | Baga NP | 150.6289738 | -23.2169732 | Conifer/Cycad sp | 0.709395 | 0.000010 |
| MtE_V43 | Felsites | Mount Hedlow Trachyte |  | 150.57845 | -23.1211 | *Eucalyptus* sp. | 0.707608 | 0.000007 |
| MtE_V6 | Mafites | Native Cat Andesite | Native Cat Rd | 150.1666 | -23.4962 | *Eucalyptus* sp. | 0.704561 | 0.000006 |
| MtE_VB15 | Ultramafic Rock | PLPzx-YARROL/SCAG | Pershouse Rd | 150.4327611 | -23.139359 | *Eucalyptus* sp. | 0.707647 | 0.000016 |
| MtE_V28 | Ultramafic Rock | PLPzx-YARROL/SCAG | Pershouse Rd | 150.43418 | -23.14108 | *Eucalyptus* sp. | 0.709102 | 0.000014 |
| MtE_V8 | Arenite | Precipice Sandstone | Sandy Creek | 150.26309 | -23.53427 | *Eucalyptus* sp. | 0.705993 | 0.000006 |
| MtE_V46 | Granitoid | Prg/g-YARROL/SCAG | Yeppoon Rd | 150.60123 | -23.25603 | *Eucalyptus* sp. | 0.706839 | 0.000008 |
| MtE_V2 | Alluvium | Qa-QLD | Somerset Rd | 150.39729 | -23.46671 | *Eucalyptus* sp. | 0.705000 | 0.000011 |
| MtE_V4 | Alluvium | Qa-QLD | Power Station Rd | 150.35372 | -23.48546 | *Eucalyptus* sp. | 0.706722 | 0.000013 |
| MtE_V17 | Alluvium | Qa-QLD | Stirling Drive | 150.51212 | -23.28076 | *Eucalyptus* sp. | 0.706456 | 0.000006 |
| MtE_V42 | Alluvium | Qa-QLD | Hedlow Creek | 150.58464 | -23.10693 | *Eucalyptus* sp. | 0.706794 | 0.000008 |
| MtE_V48 | Alluvium | Qa-QLD | Balnagowan Rd | 150.73406 | -23.45626 | Casuarina sp | 0.706919 | 0.000011 |
| MtE_V50 | Alluvium | Qa-QLD | Fitzroy River | 150.7362 | -23.48839 | *Eucalyptus* sp. | 0.708358 | 0.000007 |
| MtE_V1_  Replicate | Alluvium | Qha-QLD | Fitzroy River | 150.51636 | -23.37453 | *Eucalyptus* sp. | 0.707641 | 0.000007 |
| MtE_V1 | Alluvium | Qha-QLD | Fitzroy River | 150.51636 | -23.37453 | *Eucalyptus* sp. | 0.707646 | 0.000008 |
| MtE_V32_  Replicate | Colluvium | Qr-QLD | Stoney Creek Rd | 150.67546 | -22.92269 | Pine tree sp? | 0.711022 | 0.000011 |
| MtE_V32 | Colluvium | Qr-QLD | Stoney Creek Rd | 150.67546 | -22.92269 | Pine tree sp? | 0.711016 | 0.000011 |
| MtE_V15 | Granitoid | Ridgeland Granodiorite | Glenroy Rd | 150.11423 | -23.18798 | *Eucalyptus* sp. | 0.706323 | 0.000007 |
| MtE_VB10 | Sedimentary Rock | Rockhampton Group | Rodeo Grounds | 150.4583 | -23.17226 | *Heptapleurum actinophyllum* | 0.707548 | 0.000027 |
| MtE_V18 | Sedimentary Rock | Rockhampton Group | Meldrum Rd | 150.47259 | -23.25481 | *Ficus* sp. | 0.707998 | 0.000007 |
| MtE_V24 | Sedimentary Rock | Rockhampton Group | The Caves Quarry | 150.4470482 | -23.1632365 | *Eucalyptus* sp. | 0.705898 | 0.000010 |
| MtE_V25 | Sedimentary Rock | Rockhampton Group | Old Water Station | 150.43919 | -23.16751 | *Eucalyptus* sp. | 0.705817 | 0.000010 |
| MtE_VB16 |  | Rockhampton Group | The Caves Quarry | 150.4470482 | -23.1632365 | Grass sp | 0.706367 | 0.000016 |
| MtE_V10 | Mafites | Rookwood Volcanics | Bridgeman Rd | 150.15404 | -23.635 | *Eucalyptus* sp. | 0.705269 | 0.000007 |
| MtE_VB8 |  | Siltstone |  | 150.45367 | -23.15941 | Python Tree | 0.709099 | 0.000007 |
| MtE_V8_B |  | Siltstone |  | 150.45367 | -23.15941 | Python Tree - Bark | 0.706591 | 0.000006 |
| MtE_V9 | Granitoid | Umbrella Granodiorite | Sandy Creek Road | 150.26476 | -23.556059 | *Ficus* sp. | 0.705005 | 0.000006 |
| MtE_VB7 |  | Volcanics | Western Bench 9/10 | 150.45397 | -23.15887 | White Cedar | 0.709083 | 0.000007 |
| MtE_V40 | Arenite-Mudrock | Wandilla Formation | Old Byfield Rd | 150.66198 | -23.13191 | *Eucalyptus* sp. | 0.710278 | 0.000005 |
| MtE_V41 | Arenite-Mudrock | Wandilla Formation | Lake Mary Rd | 150.64262 | -23.09854 | *Eucalyptus* sp. | 0.707963 | 0.000006 |

* Indicates samples with potential anthropogenic modification - These values have been excluded from further analyses

**Table S3.** Mean drift corrected δ^13^C isotope values (± StdDev) measured in a series of IAEA Standards.

| Standard name | δ13C, ‰ on the VPDB δ13C scale | Combined standard uncertainty at 1σ-level | Reference | Average of Normalised δ^13^C | StdDev of Normalised δ^13^C | Count of Normalised δ^13^C |
| --- | --- | --- | --- | --- | --- | --- |
| IAEA-603 Calcite * | 2.46 | 0.01 | IAEA (2016) | 2.51 | 0.05 | 16 |
| IAEA-611 Carbonate* | -30.795 | 0.04 | IAEA | -30.79 | 0.03 | 12 |
| NBS19 Limestone* | 1.95 | not reported | IAEA | 1.92 | 0.04 | 26 |
| NBS18 Calcite | -5.014 | 0.035 | IAEA | -5.01 | 0.06 | 16 |
| Marble | NA | NA | NA | 2.10 | 0.06 | 26 |

*Denotes reference material used for data normalisation
